# DNA G-quadruplexes for native mass spectrometry in potassium: a database of validated structures in electrospray-compatible conditions

**DOI:** 10.1101/2020.11.06.371393

**Authors:** Anirban Ghosh, Eric Largy, Valérie Gabelica

## Abstract

G-quadruplex DNA structures have become attractive drug targets, and native mass spectrometry can provide detailed characterization of drug binding stoichiometry and affinity, potentially at high throughput. However, the G-quadruplex DNA polymorphism poses problems for interpreting ligand screening assays. In order to establish standardized MS-based screening assays, we studied 28 sequences with documented NMR structures in (usually 100 mM) K+, and report here their circular dichroism (CD), melting temperature (*T*_m_), NMR spectra and electrospray mass spectra in 1 mM KCl/100 mM TMAA. Based on these results, we make a short-list of sequences that adopt the same structure in the MS assay as reported by NMR, and provide recommendations on using them for MS-based assays. We also built an R-based open-source application to build and consult a database, wherein further sequences can be incorporated in the future. The application handles automatically most of the data processing, and allows generating custom figures and reports. The database is included in the *g4dbr* package (https://github.com/EricLarG4/g4dbr) and can be explored online (https://ericlarg4.github.io/G4_database.html).

## INTRODUCTION

Nucleic acids constitute the fundamental biomolecular machinery to transfer genetic information, but are also involved in the regulation of gene expression (1). Besides the canonical double helix, nucleic acids can adopt various non-canonical structures i.e. triplexes, slipped hairpins, four-way junctions, left-handed Z-form, cruciform, G-quadruplexes or i-motifs (2). G-quadruplexes (G4s) have been the subject of intense structural and biological research, given their roles in gene regulation and other related cellular processes (3,4). G4s indeed have important biological effects in replication, transcription, translation, mutagenesis, genome damage repair, telomere maintenance, RNA splicing (3,5,6). Their key role in different cellular processes makes them crucial drug targets for diseases (6–8). Besides biological roles, G4s have also numerous other applications in theranostics, supramolecular chemistry, nanotechnology, etc (6,9–12).

The building block is a G-quartet wherein four guanines adopt a square planar arrangement stabilized by eight Hoogsteen hydrogen bonds (4). The stacking of adjacent G-quartets is further stabilized by coordination with monovalent or divalent cations positioned in-between G-quartets (4,13). The guanine repeats are connected by loops. Consequently, the G-quadruplex topologies are conveniently categorized as parallel (4 strands in the same direction), antiparallel (2 strands in one direction and 2 in the other direction) and hybrid (3 strands in one direction and 1 strand in the reverse direction). Further sub-classes can be defined depending on the number of G-quartets, or which loop is lateral, diagonal or propeller. These topologies are themselves linked with the conformation of the glycosidic bond angle between the guanine base and sugars, to the stacking arrangement giving rise to specific circular dichroism signals, and to the groove width distribution (14,15).

Different analytical methods are routinely used to characterize G-quadruplex structures and their interaction with ligands (small molecule, proteins, cations, co-solvents etc.). Each of the techniques has their own experimental limitations in terms of analytes (e.g. oligonucleotide size, concentration, thermodynamic stability, labeling), buffers (e.g. cation nature and concentration, ionic strength, pH, volatility), and conditions (e.g. temperature, pressure). Studies on human telomeric sequences (TTAGGG repeats), in particular, have revealed that minor changes in the oligonucleotide sequence or in the buffer conditions can alter the structure. At least eight different types of intramolecular G4 topologies were identified to date and some sequences are inherently polymorphic (13,16,17). In the absence of external factors (i.e. co-solvents, proteins), the nature and concentration of cations predominantly affect the quadruplex conformation of a given sequence (18,19). A seminal example of this issue is the 22-mer human telomeric sequence (22AG in this manuscript), which has been assigned as a parallel quadruplex by crystallography in potassium conditions (PDB: 1KF1), an antiparallel quadruplex by NMR in sodium conditions (PDB: 143D), and a mixture of topologies (hybrid & antiparallel) in circular dichroism of potassium-containing solutions (20–22). The comparison of results obtained by different groups, using different methods and/or experimental conditions might therefore not always be directly possible, and should always be questioned.

In this manuscript, we attempt to facilitate such comparisons by native electrospray mass spectrometry (ESI-MS) (23–25). Native MS of nucleic acid is often performed in ammonium acetate, which is compatible with G-quadruplex formation (24,26). Potassium is, however, more physiologically relevant than ammonium, and more quadruplex structures have been solved in K^+^ solution in comparison to other cations. To directly compare native MS data to the literature and to work with the physiologically relevant cation, it is therefore desirable to perform ESI-MS of potassium-containing samples. Therefore, since 2014 we use ESI-MS solutions can containing up to 1 mM KCl while the ionic strength is ensured by (typically 100 mM) trimethylammonium acetate (TMAA) which can more efficiently suppress nonspecific alkali adducts than NH_4_OAc (23). The number *n* of K^+^ ions bound to the sequence indicates the number *n*+1 of G-quartet in the observed sub-ensemble, and thus the MS-derived K^+^ binding constants are linked to the G-quadruplex folding constants (27). This was exploited for equilibrium, kinetics, thermal denaturation and ligand binding studies (22,28,29).

These electrospray-compatible solution conditions were not yet used for systematic studies such as ligand screening. One possible limitation is that in 1 mM K^+^ all the quadruplexes are less stable than in high salt (~100 mM K^+^) and as a result formation of misfolded/alternative folded & non folded species can occur to a significant extent. Alternative sample preparation methods including co-solvents (hexafluoroisopropanol, isopropanol) have been proposed to increase signal-to-noise, but the risk is to induce conformational changes (25,30,31). Therefore, in order to interpret the MS results in the light of an NMR-derived structure in K^+^ we need systematically verify by solution spectroscopy (CD, UV melting, and ^1^HNMR) that the conformation reported in high salt conditions is also the one present in native MS conditions. The objective of the present study is to build a database of G-quadruplex sequences with sufficient stability and validated folds (based on UV melting, circular dichroism and NMR spectroscopies) in 1 mM KCl+100 mM TMAA assay conditions. This will help short-listing sequences for future MS-based ligand screening studies in terms of structural selectivity.

## MATERIAL AND METHODS

### Materials

Oligonucleotides were purchased in lyophilized form with RP cartridge purification from Eurogentec (Seraing, Belgium). They were dissolved in nuclease-free grade water from Ambion (Fisher Scientific, Illkirch, France/ (Ambion, Life technologies SAS, Saint-Aubin, France). The concentration of the stock solutions was determined using absorbance at 260 nm and molar extinction coefficients calculated using the nearest-neighbor model in its traditional format (eq. 1), where ε_*i*_ is the molar extinction coefficient (in M^−1^cm^−1^) of the nucleotide in position *i* (in the 5’to 3’ direction), ε_i,i+1_ is the extinction coefficients for doublets of nucleotides in positions *i* and *i* + 1, and *N*_b_ is the number of nucleotides in the oligonucleotide (32,33).

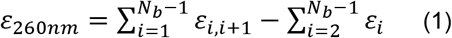

The implementation of Eq. (1) in the g4dbr application is provided in the g4dbr manual (Supporting information). All molar extinction coefficient values are provided in supporting information.

All sequences used in this study are listed in Table 1. Trimethylammonium acetate (TMAA, Ultra for UPLC, Fluka) and potassium chloride solution (1M concentration) (KCl, >99.999%), KH_2_PO_4_, K_2_HPO_4_, and D_2_O were purchased from Sigma-Aldrich (Saint-Quentin Fallavier, France). The stock oligonucleotide solutions were diluted to 100 μM in 100 mM TMAA supplemented with 1 mM KCl in water (pH 7.0) and kept at least 72 hours at 4°C to ensure G-quadruplex formation. For telomeric sequences, no annealing was done, but previous studies showed that the end point will be reached after 72h (22). For non-telomeric sequences, the solutions were annealed at 85 °C for 3-4 minutes in a water bath then let at room temperature for 24 hours before use. For UV melting and CD two buffers were used: (1) 100 mM TMAA supplemented with 1 mM KCl (MS-compatible buffer) and (2) 5-25 mM K_2_HPO_4_/KH_2_PO_4_ buffer supplemented with KCl (pH 7.0-7.1) (Table S1).

**Table 1.**
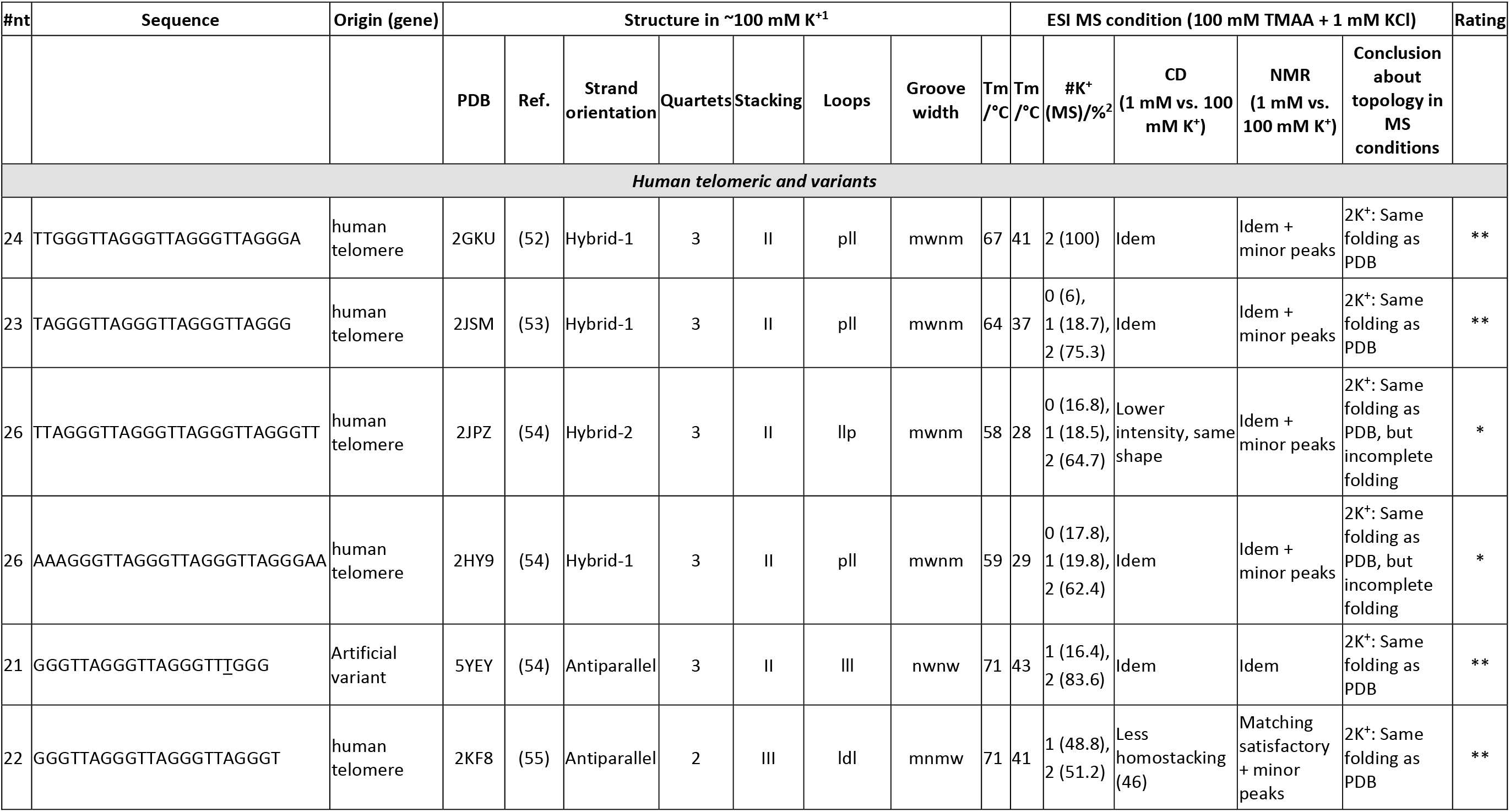
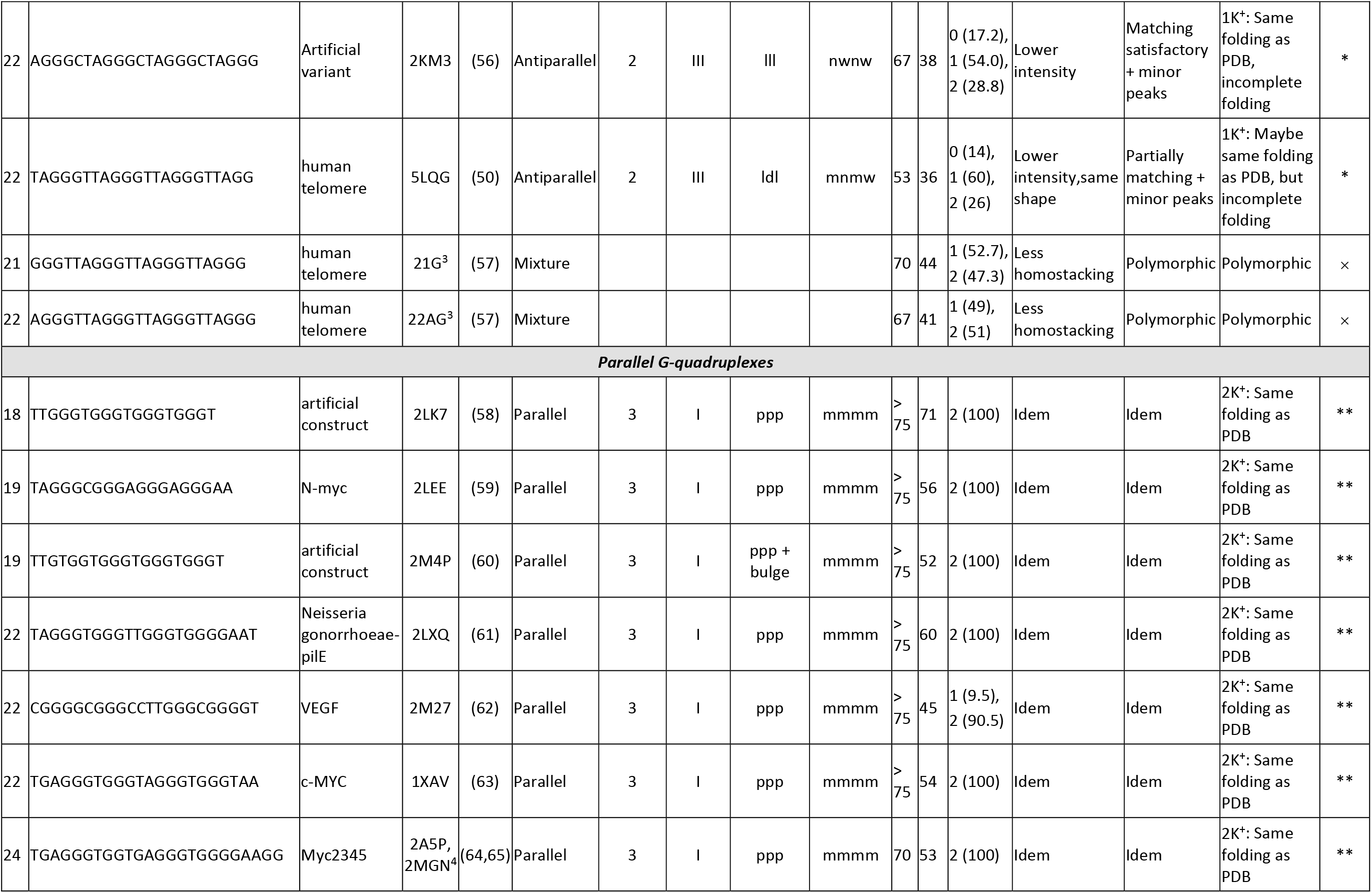
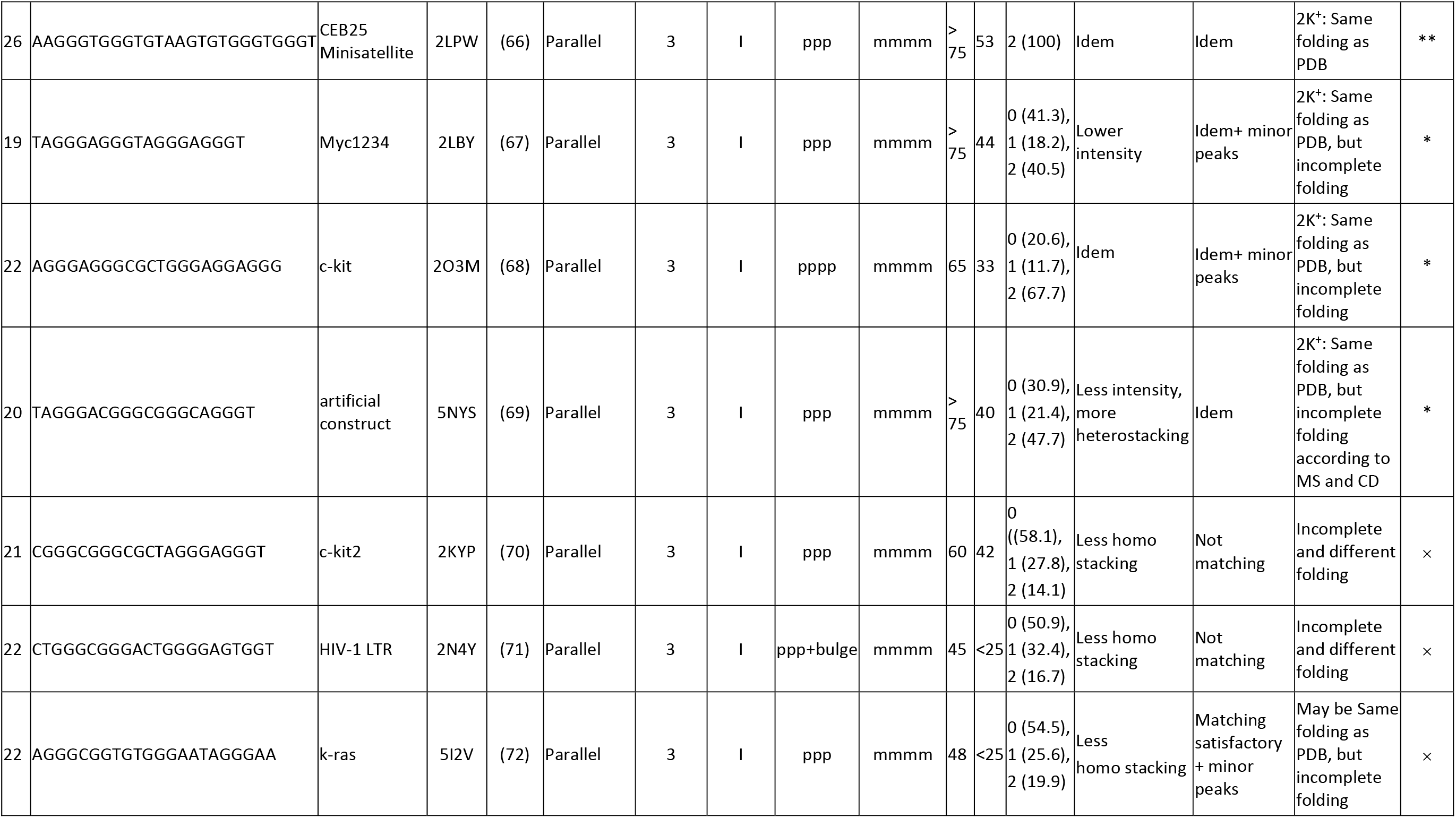
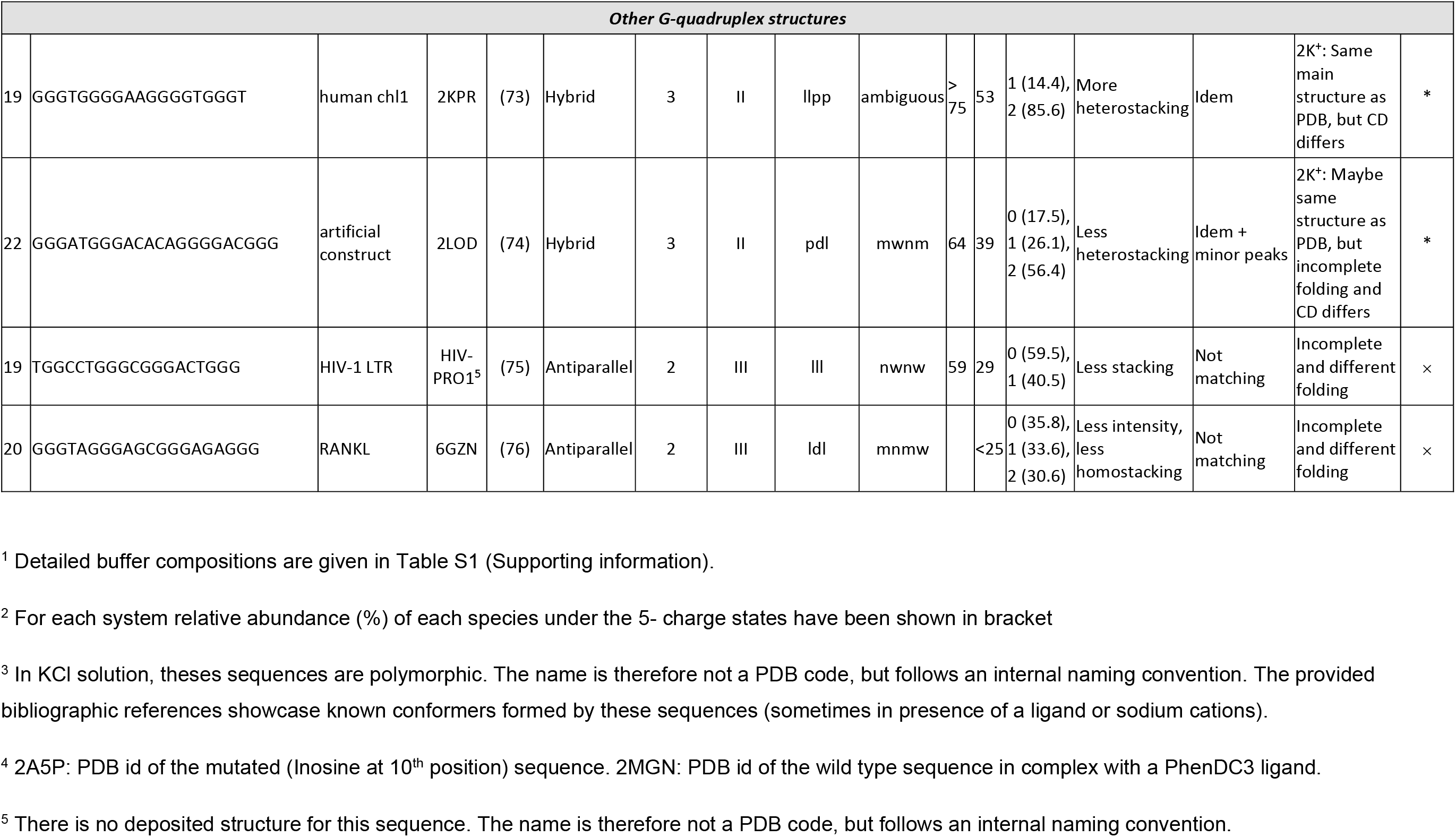
Oligonucleotides used in this work and described in detail in the database. The respective sequence, origin, structural features in both conditions and rating are summarized below.

### Circular dichroism (CD)

All circular dichroism experiments were performed on a Jasco J-815 spectrophotometer equipped with a JASCO CDF 426S Peltier temperature controller using a quartz cuvette (2 mm path length) at 25 °C. The DNA concentration was 10 μM for all the measurements. The scanning range was 220-320 nm with 0.2 nm data pitch, 2 nm bandwidth, and 0.5-sec response. For each sample, 3 accumulations were acquired with a scan speed of 50 nm/min. For each average spectrum of each sample, baseline subtraction was performed with the corresponding buffer without the DNA. The subtracted spectra were normalized to molar ellipticity coefficient (Δ*∊*) according to the following Equation (2):

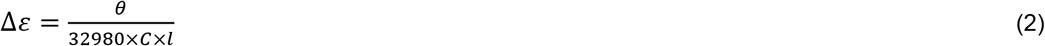

where θ = CD signal in millidegrees, *C* = DNA concentration in mol/L and *l* = path length in cm.

### Melting monitored by UV absorbance (UV-melting)

Melting temperatures were determined by measuring the changes in absorbance at 295 nm as a function of temperature, using a UVmc2 double-beam spectrophotometer (SAFAS, Monte Carlo, Monaco) equipped with a high-performance Peltier temperature controller and a thermostatable 10-cell holder, with 400-μL, 1-cm pathlength quartz cuvettes (115B-QS, Hellma GmbH & Co. KG, Müllheim, Germany). The samples contained the oligonucleotide (10 μM) in potassium phosphate or TMAA buffers, supplemented or not by potassium chloride, and were cooled to 4 °C. The absorbance was monitored at 260, 295, and 335 nm on a cycle composed of a heating to 90°C at a rate of 0.2 °C min^−1^, then cooling to 4°C at the same rate.

The raw absorbance data was buffer subtracted, and converted to molar extinction coefficient ε (in M^−1^ cm^−1^) using ε = *A*/*lC*, where *l* is a path length (in cm) and *C* the oligonucleotide concentration (in M). The melting temperatures (*T*_m_), determination for a 2-state equilibrium, and the conversion of the temperature-dependent absorbances *A*_T_ into folded fractions θ_T_, were carried out based on a non-linear fitting-based implementation of the baseline method, using eq. 3 where *a* and *b* are the slopes and intercepts, respectively, of the folded (*F*) and unfolded (*U*) baselines, *R* is the gas constant (in J.K^−1^.mol^−1^), and *T* is the temperature (in K) (34).

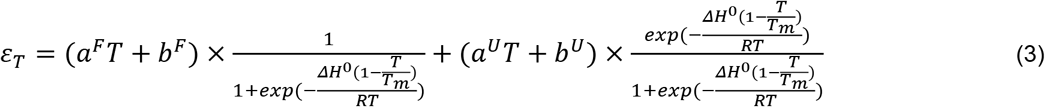

θ_T_ gives a direct access to the extent of folding of an oligonucleotide (1: all molecules entirely folded, 0: all molecules entirely unfolded), allows to visually assess the *T*_m_ (θ_t_ = 0.5), and normalize the data of different samples (and therefore different absorbances) to a common y-scale (34). For the non-linear fitting and the folded fraction calculation to be carried out, the data must contain both *lower* and *higher* baselines. When this was not the case (the oligonucleotide is too stable or unstable), the melting curves were simply normalized to [0;1], and no thermodynamic quantities were determined.

The derivation of Eq. (3) and its implementation in the g4dbr application are provided in the g4dbr manual (Supporting information).

### Nuclear magnetic resonance (NMR)

All ^1^HNMR experiments were carried out on a Bruker 700 MHz spectrometer (Bruker biospin) equipped with 5 mm TXI probe at 25 °C. The jump-and-return water suppression is used in all experiments (35). The sweep widths were 20 ppm with a 3-sec relaxation delay with a size of 32K data points per 1D spectra. The number of scans and dummy scans was 128 and 16 respectively. The 1D raw data were processed and analyzed with Topspin 4.06 software inbuilt with the instrument. All the quadruplex sequences were 100 μM strand concentration in 100 mM TMAA+1mM KCl in a 5 mm NMR tube (Wilmad from CortecNet, France).

### Electrospray mass spectrometry (ESI-MS)

All ESI-MS experiments were performed in negative ion mode on an Agilent 6560 IMS-Q-TOF (Agilent Technologies, Santa Clara, CA, USA) with a dual ESI source and soft tuning conditions (36). The experiments were performed in ion mobility mode (*p*He = 3.89 ± 0.01 torr, *T* = 296 ± 1 K). The source gas temperature was 200°C with fragmentor voltage at 350 V (soft conditions, by default). The injected DNA concentrations were 10 μM G-quadruplex in 100 mM TMAA and 1 mM KCl (180 uL/h flow rate with a syringe pump). The ESI-MS spectra are described herein, and the ion mobility results will be described in a separate paper.

### Data treatment, app and database

Circular dichroism, UV-melting, NMR, and native ESI-MS data filtering, normalization, fitting, and labeling was performed in g4db, an in-house Shiny application included in the g4dbr package, written in RStudio 1.3.1056,^1^ running R 4.1(37,38). The database is included in the *g4dbr* package (https://github.com/EricLarG4/g4dbr) and can be explored online (https://ericlarg4.github.io/G4_database.html) (39).The application documentation is provided in supporting information.

## RESULTS

### 1. Database composition and building

We examined 28 sequences (among which 10 human telomeric variants) for which a specific G-quadruplex structure had been solved by NMR in high KCl concentration (usually ~100 mM), with the exception of two polymorphic sequences 22AG and 21G. The sequences will be referred to by their PDB code as listed in Table 1 (see also Tables S2 to S29). The CD, ^1^HNMR, ESI-MS spectra and UV thermal denaturation profiles are all shown in the supporting information (Figures S1 to S168). Selected results are discussed in the following sections.

Our goal was to gather diverse topologies from the literature, with NMR data in potassium. The published solution structures in potassium include many parallel topologies with Type I base stacking, some hybrid topologies (in particular, among human telomeric sequences) with Type II stacking, some antiparallel topologies with 2 G-quartets (Type III stacking), one antiparallel topology with 3-quartet but a Type II stacking (5YEY) (Figure 1A-E). There is no documented antiparallel topology with 3-quartets and a purely Type III stacking in potassium; such structures are documented only in sodium (Figure 1F) (40).

**Figure 1:**
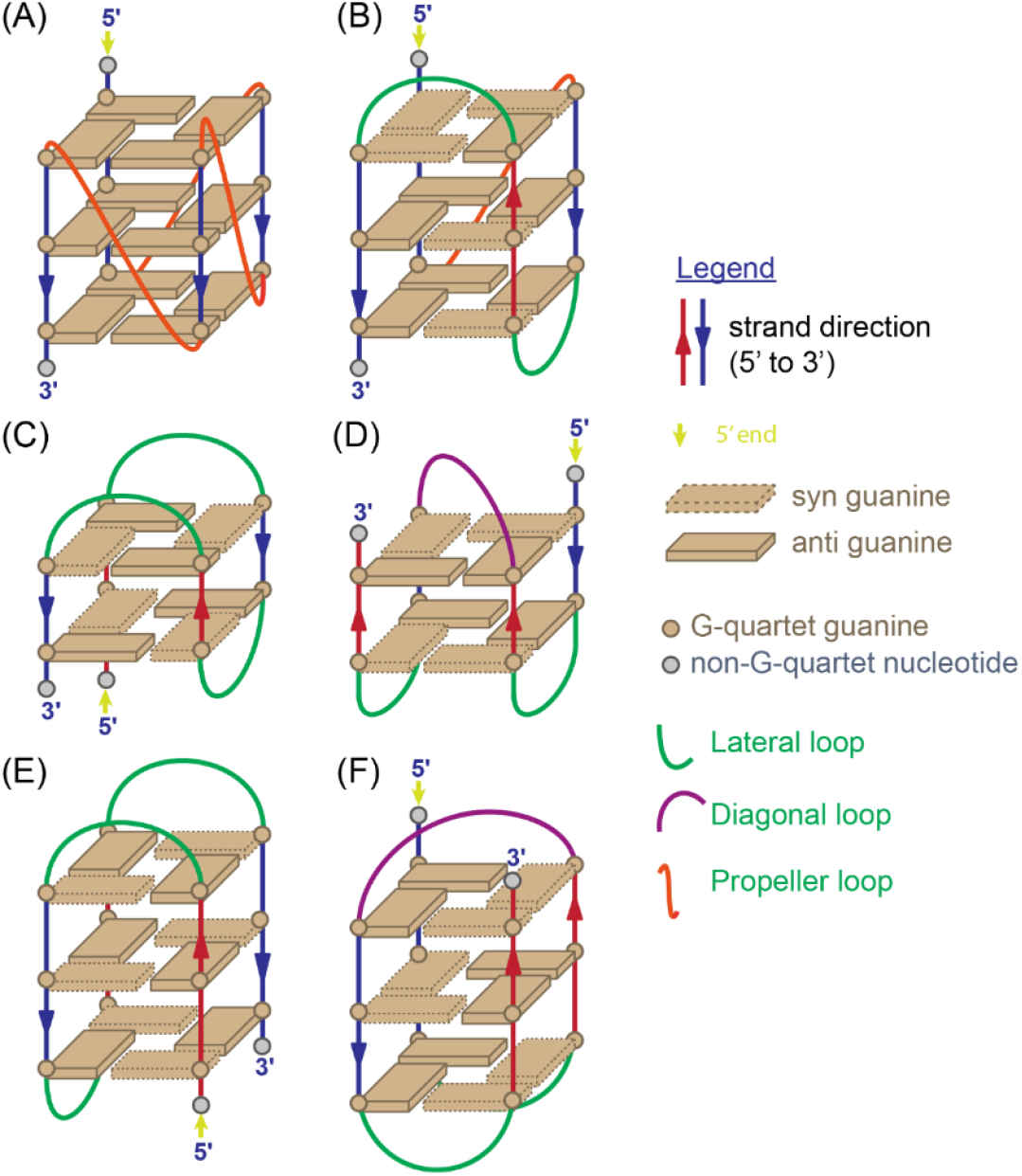
Schematic representation of different G4 topologies. (A) Parallel (Type I stacking), (B) Hybrid (Type II stacking), (C) Antiparallel 2-quartet (Type III stacking with chair conformation), (D) Antiparallel 2-quartet (Type III stacking with basket conformation), (E) Antiparallel 3-quartet (Type II stacking with chair conformation), (F) Antiparallel 3-quartet (Type III stacking with basket conformation).

Among bimolecular G quadruplex sequences, (12TAG)_2_ and (G_4_T_4_G_3_)_2_ are not stable enough in 1 mM K^+^, instead they exists as a single strand (41,42). In ESI MS (G_4_T_4_G_4_)_2_ and (G_4_T_3_G_4_)_2_ shows (M+3K)^n−^ (n=4,5,6) as major peaks corresponding to 4 quartet antiparallel conformation while (G3T4G4)_2_ form (M+2K)^n-^ (n=4,5,6) in 1 mM KCl (3 quartet antiparallel conformation) solution, in agreement with the solution NMR studies (24,41,43,44). The other possibility is to not stick to the 1 mM KCl/100 mM TMAA preparation. The propensity of cations to favor parallel structures follows the trend: K^+^ > NH_4_^+^ > Na^+^(45). In this context, instead of KCl/TMMA pair one can also use Ammonium acetate (NH_4_Oac) for the bimolecular quadruplexes where there is sufficient evidence of conformation and NH4 binding site from solution NMR studies. The 4 quartet antiparallel sequences i.e. (G_4_T_4_G_4_)_2_ & (G_4_T_3_G_4_)_2_ have 3 NH_4_ ^+^ binding sites with strong affinity while (G_3_T_4_G_4_)_2_ has 2 NH_4_^+^ binding sites (24). Such sequences will be added to the database in the future.

The database was built using an in-house, open-source R package, *g4dbr* (https://github.com/EricLarG4/g4dbr). Specifically, the *g4db* function is dedicated to the processing, tidying, storing, visualization, and reporting of CD, ^1^H-NMR, UV-melting and native mass spectrometry (MS) data from oligonucleotide samples. Although developed for the G-quadruplex forming sequences characterized in this manuscript, *g4db* can be used with any nucleic acid sequence. The long-term goal is to provide open-source tools for the deposition of oligonucleotide biophysics data (raw and processed), while allowing for easy and versatile visualization and reporting.

In practice, users can employ the app to visualize a database previously generated by *g4db* in the *database* module, or visualize and process raw data, then import it into a new or existing database using the *importR* module (Figure S169). To read raw data in *g4db*, it must be first pasted into a templated Excel file provided in the package. Several tools are included in the package for the automated processing of data, and can also be used outside of the database scope. The most notable are:

- Data filtering by oligonucleotide name, sequence, topology, buffer, cation, and selective writing to databases;
- Labeling of MS spectra performed from user-supplied species names, for which the expected *m/z* are calculated by the application;
- Calculation of molar extinction coefficients at 260 nm from the oligonucleotide sequence, using eq. 1. This can be used independently from g4db, using the *epsilon*.*calculator* function (included in the g4dbr package);
- Conversion of CD data in mdeg to molar extinction coefficient, using eq. 2;
- Determination of folded fraction vs. temperature, and *T*_m_, from UV-melting data (*meltR* module), using eq. 3. This will be released as a standalone application, and its performance will be discussed in a separate publication;
- MS noise reduction by intensity filtering, which can be carried out independently with the *mass.diet* function (included in the package).

The processed data is consolidated into.Rda files, which can either be consulted in *g4db*, or can be loaded in base R for uses outside the application scope. In *g4db*, the data can be visualized with several customizable plots, and exported in reports in word, pdf, or HTML formats. The reports generated for the oligonucleotides characterized herein are collated in supporting information, and are accessible online as well (https://ericlarg4.github.io/G4_database.html). The use of g4db is described in supporting information (Figures S170–S192) and and whole application deposited to zenodo repository (39).

### 2. Stability in ESI-MS conditions (1 mM KCl/100 mM TMAA) compared to ~100 mM KCl

The melting temperatures are systematically lower in 1 mM KCl than in 100 mM KCl for all the sequences studied. As a result, for some molecular systems, the decrease is such that, at room temperature, a fraction of the oligonucleotide is not folded. In the ESI-MS spectra, this translates into the appearance of peaks with lower-than-predicted number of K^+^ ions bound. The number of K^+^ ions predicted to bind in the G-quadruplex core is *n-1*, with *n* the number of stacked G-quartets. However, whether all sites are fully filled depends on the K^+^ binding equilibrium constants and on the KCl concentration.

For the 3-quartet G-quadruplexes, there is a good correlation between the melting temperature, the fraction folded at 25°C and the relative intensity of the 2-K+ complex in the ESI-MS conditions. The 2-K^+^ abundance increase with the folded fraction or Tm for 3-quartet G4s. Usually, if *T*_m_ > 40°C (which corresponds to fraction folded at 25°C > 80%), there is no 0K (non-folded) in ESI-MS conditions (Figure 2). The exceptions are 2M27, 2KPR and 5YEY where there is still ~9%,14% & 16% 1-K+ complex with a Tm of 45 °C, 53°C & 43°C respectively (Fraction folded at 25°C~99%, 100%, 100% respectively). The three outliers are 2LBY, 5NYS and 2LOD (“*” rating in Table 1) for which the combined abundance of 0-K and 1-K complex is ≥42% even though they have a flat low-temperature baseline in UV-melting. This suggests that under such flat baselines, several species can coexist, if the transition is due only to one of these species. This would require further exploration by temperature-resolved mass spectrometry.

**Figure 2.**
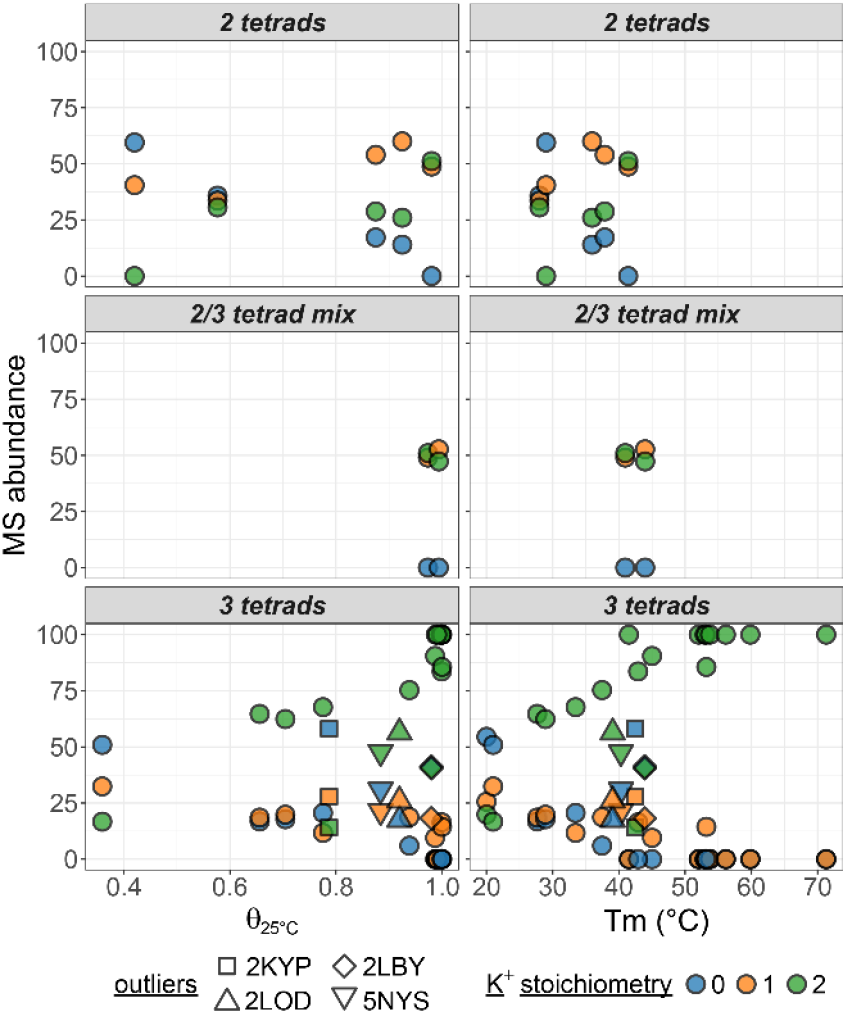
Species abundance in native ESI-MS experiments against their folded fraction at 25°C (left) or melting temperature (right) determined by UV-melting. The oligonucleotides are grouped in panels by their number of G-quartets, and colored by their number of potassium adducts. Outlying sequences are shown with a specific shape.

The trend is similar with 1-K^+^ (2-quartet), but few data points are available. Note that the interpretation of the K^+^ distribution in antiparallel 2-quartet structures is peculiar. The case of the 2-quartet 2KF8 was discussed in detail previously (where it was named 22GT) (46). In 2KF8, the main stoichiometry is 1K^+^, and the 2K^+^ complex reflects a second lower-affinity binding site between a quartet and a triplet, which upon stacking also changes the CD spectrum. We can thus imagine that in the 1K^+^ complex the G-triple is not structured, while it is present in the 2K^+^ complex. In contrast, sequences that do not involves stable base pairing (triplex or G-C-G-C) above the quartet at 1mM K^+^(2KM3, 5LQG) do not take up so much of a second K^+^ ion.

### 3. Solution structure in ESI-MS conditions (1 mM KCl/100 mM TMAA) compared to 100 mM K^+^

The ^1^H-NMR spectra recorded in 1 mM KCl and 100 mM TMAA were compared with those published in the literature, and all peaks (in the imino region) that were matching are labeled according to the published base number assignment. In some cases minor peaks were present (as indicated in the table), but note that in 100 mM K^+^ minor other conformations were also noticed for the wild type sequences. In such case we would still conclude that the main topology in ESI-MS conditions are the same as in the PDB.

We also compared the CD spectra obtained in 100 mM KCl and in 1 mM KCl + 100 mM TMAA, and noted when the shapes were identical, showed more homo-stacking (larger relative signal at 260 nm) or more hetero-stacking (larger relative signal at 290 nm). Parallel topologies show exclusively anti-anti guanine stacking (type I stacking) characterized in CD spectra by positive maximum at ~265 nm and negative maxima at 245 nm. Hybrid G-quadruplexes combine syn/anti and anti/syn with anti/anti stacking (type II stacking) in the topology which is represented by two positive peaks at 270 and 290 nm and 1 negative minimum at 245 nm. Antiparallel topologies usually display alternative stacking of syn/anti and anti/syn guanines (type III stacking) leading to the CD positive maxima at 290 nm and CD negative minima at 260 nm (47). Shape changes can hint at different structural populations, and when noticed, the confidence in having the same structure in ESI-MS conditions as in the published conditions was lower.

In Table 1 we rated each sequence. Two stars (**) means that the folding is >90% complete in the same topology as formed by NMR. One star (*) means that either the folding is incomplete, or that there are doubts that the topologies formed are the same in 1 mM and 100 mM KCl. Sequences with incomplete folding can be problematic for ligand screening by ESI-MS, because ligands that bind to the folded fraction also have to displace the folding equilibrium, and therefore the apparent binding affinity is the convoluted result of folding and binding equilibria. Nevertheless, it is still possible that the folded fraction has the same structure as the 100 mM K+ (NMR) structure, and that these sequences might thus still be of use for structural or specificity studies. Below we discuss the behavior of different G-quadruplex families in more detail.

#### 3.1. Human telomeric sequences and variants

This group includes one 3-quartet antiparallel topology (5YEY), four 3-quartet hybrid topologies (2JSM, 2GKU, 2HY9, and 2JPZ), three 2-quartet antiparallel topologies (2KF8, 2KM3, 5LQG), and two polymorphic sequences (21G, 22AG). The polymorphic sequences show mixed conformational topology from CD and broad imino proton signal in ^1^HNMR spectra in the ESI-MS buffer as well. Their mass spectra show both [M+1K]^n−^ and [M+2K]^n−^ (n=4,5,6) stoichiometries, indicating the presence of 2-quartet topologies in the mixture (Figure 3).

**Figure 3:**
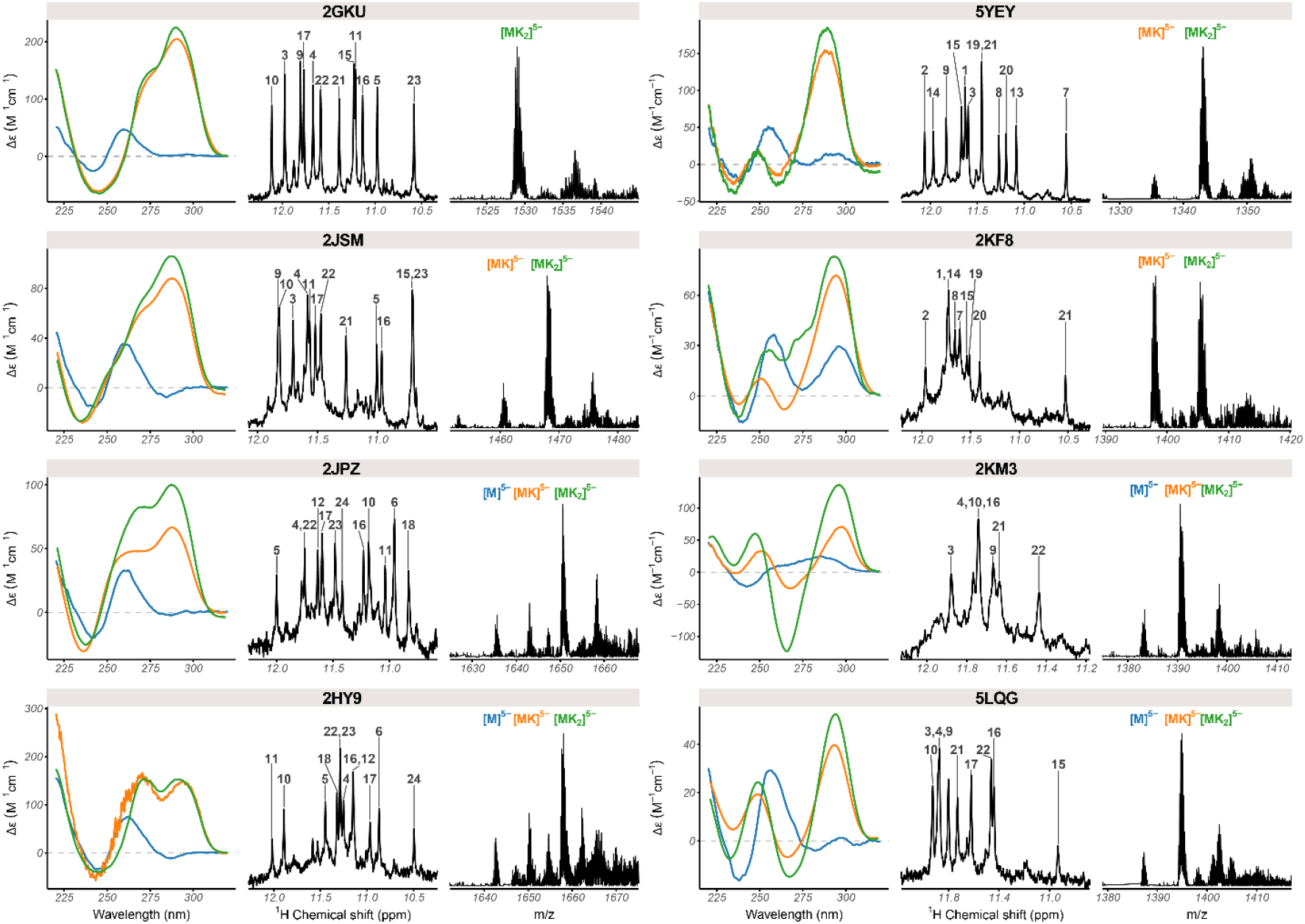
Circular dichroism (left; 10 μM DNA in blue: 100 mM TMAA, orange: 100 mM TMAA + 1 mM KCl, green: potassium phosphate + KCl), ^1^H NMR (center; 100 μM DNA in 100 mM TMAA + 1 mM KCl), and native ESI-MS (right; 10 μM DNA in 100 mM TMAA + 1 mM KCl; M: monomer, K: potassium) data of selected human telomeric quadruplex-forming oligonucleotides.

For all hybrid topologies, we observe the 3-quartet [M+2K]^n−^ complex as the major species in 1 mM K^+^. However, for most of the hybrid G4 sequences (except 2GKU) we observe a substantial population of 2-quartet [M+1K]^n−^ and non-folded species [M+0K]^n−^, indicating an incomplete folding in 1mM KCl (2JPZ, 2HY9 & 2JSM) (Figure 3). The subpopulation of minor species is also evident from the unassigned peaks in the imino region of ^1^HNMR spectra for all the sequences. For hybrid-1 telomeric sequences, the preferred sequences for ligand screening should be 2GKU and 2JSM (the first is more stable and less polymorphic, but the overhangs are not fully faithful to telomeric repeats). For hybrid-2 we have only one sequence representative, 2JPZ, which is incompletely folded (θ_25_ = 0.65). Therefore, it would be desirable to find another suitable hybrid-2 telomeric sequences, stable enough in ESI-MS conditions for ligand screening. Note that there is also evidence of coexistence of hybrid-1 and hybrid-2 topologies (48,49), including in our ESI-MS conditions as will be detailed in the forthcoming paper describing ion mobility results.

In antiparallel sequences we have two antiparallel 2-quartet sequences with basket topology (2KF8, 5LQG) and one chair topology (2KM3). For 2KM3 and 5LQG, [M+1K]^n−^ complexes predominate in ESI MS while for 2KF8 [M+1K]^n−^ and [M+2K]^n−^ peaks coexist (Figure 3). The second K^+^ binding site is enabled by the formation of guanine triplet composed of G9, G13, and G21 in the diagonal loop (46). 5LQG can form two different antiparallel basket-type G-quadruplexes depending upon K^+^ concentration and pH as shown previously (50), and is incompletely folded, which makes it unsuitable for ligand screening in MS. Finally, for 5YEY we observe a major population of [M+2K]^n−^ and mostly one conformation according to ^1^HNMR spectra. In summary, for antiparallel sequences, the best sequences for ligand screening are 2KF8 (antiparallel 2-quartet, basket) and 5YEY (antiparallel 3-quartet, chair). 2KM3 is the only antiparallel 2-quartet with a chair conformation, but is incompletely folded (θ_25_ = 0.87).

#### 3.2 Parallel G-quadruplexes

A second group of non-telomeric (mainly promoter sequences) with validated folds are parallel-stranded with type I stacking. Eight sequences with length ranging from 18 to 26 nucleotides are fully folded (>90%) in the ESI-MS conditions. These include two artificial constructs (PDB: 2LK7, 2M4P), two c-myc promoter variants (PDB: 1XAV, 2A5P/2MGN), as well as n-myc (PDB: 2LEE), VEGF (PDB: 2M27), Neisseria gonorrhoeae-pilE promoters (PDB: 2LXQ) and the CEB25 minisatellite (PDB: 2LPW) (Figure 4). Another three sequences (2LBY, another c-myc promoter variant; 2O3M, the c-kit promoter; 5NYS, an artificial construct) also form the same parallel structure as reported by NMR, but with a significant fraction unfolded (0K). These three sequences should not be used for testing ligand preference to parallel versus other topologies, but can be used with cautious interpretation of the apparent binding constants if one is interested in these specific structures. Finally, the c-kit2 (PDB: 2KYP), HIV-1 LTR (HIV-PRO1), and k-ras (PDB: 5I2V) do not fold in the desired topology in 1 mM KCl.

**Figure 4:**
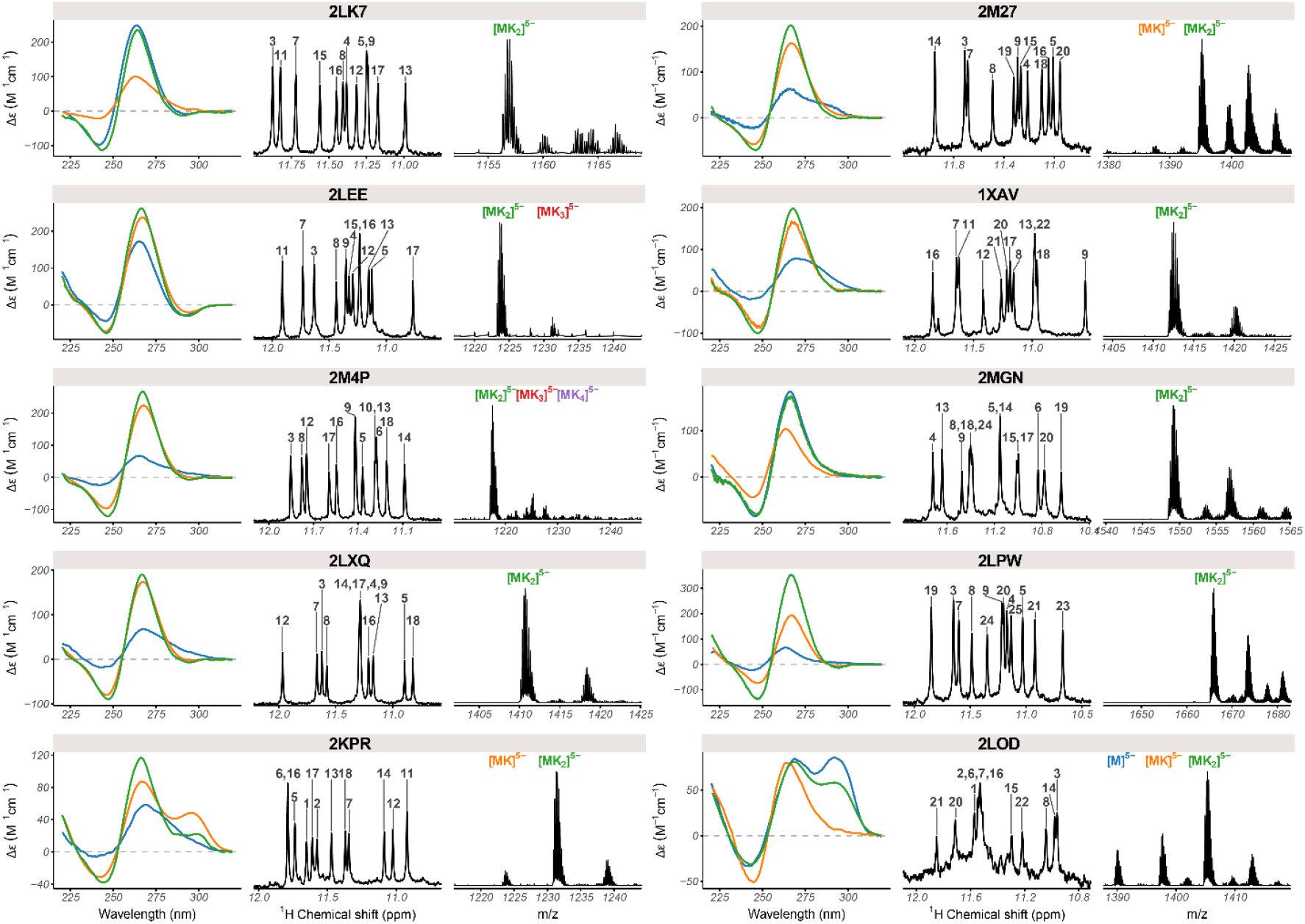
Circular dichroism (left; 10 μM DNA in blue: 100 mM TMAA, orange: 100 mM TMAA + 1 mM KCl, green: potassium phosphate + KCl), ^1^H NMR (center; 100 μM DNA in 100 mM TMAA + 1 mM KCl), and native ESI-MS (right; 10 μM DNA in 100 mM TMAA + 1 mM KCl; M: monomer, K: potassium) data of selected parallel and hybrid quadruplex-forming oligonucleotides.

#### 3.3. Other model structures

To have a few other topologies adequate for ligand screening in ESI-MS, we tested two other 3-quartet hybrid structures (2KPR and 2LOD) and two 2-quartet antiparallel structures (6GZN, HIV-PRO1). The two hybrid folds present both similarities (NMR) and differences (CD) between 100 mM and 1 mM KCl conditions (Figure 4). Both sequences have a major population of [M+2K]^n−^, but also [M+1K]^n−^ as a minor species in 1 mM K^+^. Therefore the confidence rating is low (“*”) compared to the representatives of Type I folding. The two antiparallel structures did not form the same fold in ESI-MS conditions as evident from ^1^HNMR, CD and ESI-MS.

#### 3.4. Recommendations for native MS screening

In order to short-list sequences for biophysical and ligand screening studies by native mass spectrometry, we would like to summarize the following points:

1. Pre-select sequences that can potentially be of interest due to its origin or biological function, based on the following factors:
  a. A high-resolution structure is available (probably by solution NMR in K^+^), obtained from a non-modified sequence.
  b. The structure does reflect a single or predominant/major conformation in solution.
  c. Sequences of the same length but different topology can always be an interesting point to screen for a similar group of ligands to observe “selectivity”.
2. Perform the experiments and the data treatment in a consistent manner to ensure the validity of comparison across techniques and oligonucleotides, and minimize the variability of results.
  a. Establish and use standard protocols in each experiment type for all sequences concerned,
  b. Where applicable, normalize data to streamline comparisons. For instance, CD data obtained for different oligonucleotide concentrations and/or cuvette path lengths can be more easily compared if the data is converted to Δε (eq. 2).
  c. As far as possible, eliminate human biases from the data treatment, by e.g. automation. For instance, the ‘manual’ baseline subtraction of UV-melting data is notoriously imprecise, and was automated herein (eq. 3) (34).
3. To compare with the high-resolution structure (NMR derived) in high K^+^ containing buffer it is necessary to perform additional solution spectroscopic experiments (CD, UV melting) in both conditions (1mM K^+^ and ≥ 1 mM K^+^). Finally, by comparing imino proton signals in ^1^HNMR with the published assignment, one can conclude whether a particular sequence retains its same tetrad arrangement or not. ^1^HNMR can also give some information on the formation of triads/base pairs apart from the quartet formation. Since G-quadruplexes are inherently polymorphic, at 1 mM K^+^ we also found minor populations for other misfolded species/alternative conformations as evident from the unassigned peaks in 1HNMR for most of the sequences. However, we tried to focus on comparing the major species observed in ESI-MS (3 quartet for Type I and II and 2 quartets for type-III folding) with the major species in 1HNMR. In-depth structural analysis for each of the conformers for every sequence in native MS buffer by solution NMR is, however, beyond the scope of the manuscript. We recommend to discard sequences for which a significant amount of unfolded folded species is found (^1^HNMR not matching with literature, different CD profile, θ_25_ << 1 in UV melting, and high abundance of the 0-K complex in ESI-MS).
  a. A similar CD signature (in terms of positive and negative maxima) demonstrates the same stacking of nucleotides in both conditions for a particular sequence. However, lower intensity points towards incomplete folding or less homostacking in 1mM K+. This is also supplemented by the relative abundance of each species in ESI-MS and comparison of Tm(s)/Fraction folded from UV melting in both conditions. Indeed, we have found a good correlation between the presence of a 2-K+ peak in native MS, Tm, and folded fraction (θ25) for most of the Parallel quadruplex sequences (Figure 1).
  b. The coexistence of several predominant K+ stoichiometries in native MS hints toward the formation of multiple coexisting conformers in equilibrium (e.g., 21G & 22AG). However, one has to cautious about the presence of alternate K+ binding sites other than G-quartet planes (e.g., K^+^ binding site in G triad of 5’ end of 2KF8). The K^+^ stoichiometry determined by MS, which often reflects the number of G-quartet, should be used judiciously, that is considering the other analytical techniques.
  c. The presence of predominant 1K^+^ peak in native MS, together with a significant 290-nm band in CD suggests the formation of an antiparallel quadruplex with two G-quartets (e.g., 2KF8, 5LQG, & 2KM3). On the other hand, few human telomere hybrid (e.g., 2HY9, 2JPZ, 2JSM) and other hybrid sequences (e.g., 2KPR, 2LBY, 2LOD) show the presence of non-folded [(M+0K])^n−^] and misfolded [(M+0K])^n−^] species due to the low thermodynamic stability in 1mM K+.
4. Consolidate the data in a way that provides the community with open and easy access to raw and processed data, as well as all necessary contextual elements (e.g. oligonucleotide sequence, extinction coefficient, and concentration, buffer and cation nature and concentration, literature references). A database/software suite was developed to automatize the gathering, data treatment, storage, and reporting in a completely open way. In the future, sequences with different topology and validated fold (Table 1) will be used as a target for ligand screening by native MS.

## CONCLUSIONS

Parallel 3-quartet structures are over-represented among the intramolecular structures solved to date in potassium. For non-parallel structures, the best models for ESI-MS screening are variants of the human telomeric sequence, with the known caveat that due to inherent polymorphism, ligand-induced conformational changes can occur even for sequences that have one very predominant fold in absence of ligand(28,41,51). Finding suitable candidates with intramolecular Type II & Type III folding among non-telomeric sequences for MS-based ligand screening in 1 mM K^+^ remains challenging because there are not many such structures solved already in 100 mM K^+^. Note that for now we limited our database to quadruplex sequences with all natural nucleotides.

In the future, we will continue to update the database and, in particular, we will incorporate more sequences with unusual quadruplex folding (e.g., Left-handed quadruplex, quadruplex-duplex hybrid) and bimolecular/tetra molecular quadruplexes in our online database to have a robust library of G4s. Another possibility would be to include bimolecular G-quadruplex structures in the screening panel. On careful choosing of instrumental tuning setting it is possible to visualize the solution phase equilibrium in Ammonium acetate (24).

## AVAILABILITY

The reports generated for the oligonucleotides characterized herein are accessible online as well (https://ericlarg4.github.io/G4_database.html). The whole application is deposited to Zenodo repository (DOI: 10.5281/zenodo.4200176).

## SUPPLEMENTARY DATA

Supplementary Data are available at NAR online.

## ACKNOWLEDGEMENT

This work was financially supported by the European Union (H2020-MSCA-IF-2017-799695-CROWDASSAY and ERC-2013-CoG-616551-DNAFOLDIMS), and benefited from access to NMR, MS and CD at the Plateforme de BioPhysico-Chimie Structurale of the IECB, which staff is acknowledged for their support. Prof. J. L. Mergny and his group are acknowledged for access to SAFAS spectrophotometer. The authors acknowledge Dr. Samir Amrane and Dr. Frédéric Rosu for useful discussions.

