## Supplementary Data for "DNA G-quadruplexes for native mass spectrometry in potassium: a database of validated structures in electrospray-compatible conditions"

*SUPPORTING INFORMATION*

#### *Table of content*

NMR buffers ____ 3
2GKU ____ 5
2JSM ____ 6
2JPZ ____ 7
2HY9 ____ 8
5YEY ____ 9
2KF8 ____ 10
2KM3 ____ 11
5LQG ____ 12
21G ____ 13
22AG ____ 14
2LK7 ____ 15
2LEE ____ 16
2M4P ____ 17
2LXQ ____ 18
2M27 ____ 19
1XAV ____ 20
2MGN ____ 21
2LPW ____ 22
2LBY ____ 23
2O3M ____ 24
5NYS ____ 25
2KYP ____ 26
2N4Y ____ 27
5I2V ____ 28
2KPR ____ 29
2LOD ____ 30
HIV-PRO1 ____ 31
6GZN ____ 32
g4dbr manual ____ 33

### NMR buffers

*Table S1. NMR buffer composition used to determine the structure deposited in the PDB*

| Oligonucleotide | [Potassium phosphate] (mM) | [KCl] (mM) | pH | reference |
|---|---|---|---|---|
| 2GKU | 20 | 70 | 7.0 | (1) |
| 2JSM | 20 | 70 | 7.0 | (2) |
| 2JPZ | 25 | 70 | 7.0 | (3) |
| 2HY9 | 25 | 70 | 7.0 | (3) |
| 5YEY | 20 | 70 | 7.0 | (3) |
| 2KF8 | 20 | 70 | 7.0 | (4) |
| 2KM3 | 20 | 70 | 7.0 | (5) |
| 5LQG | 5 | 70 | 7.0 | (6) |
| 21G | 20 | 70 | 7.0 | * |
| 22AG | 20 | 70 | 7.0 | * |
| 2LK7 | 10 | 0 | 7.0 | (7) |
| 2LEE | 20 | 100 | 6.5 | (8) |
| 2M4P | 20 | 30 | 7.0 | (9) |
| 2LXQ | 5 | 25 | 7.0 | (10) |
| 2M27 | 20 | 70 | 7.0 | (11) |
| 1XAV | 25 | 70 | 7.0 | (12) |
| 2MGN | 10 | 35 | 7.0 | (13, 14) |
| 2LPW | 20 | 70 | 7.5 | (15) |
| 2LBY | 25 | 70 | 7.0 | (16) |
| 2O3M | 20 | 70 | 7.0 | (17) |
| 5NYS | 20 | 100 | 7.0 | (18) |
| 2KYP | 5 | 20 | 7.0 | (19) |
| 2N4Y | 20 | 70 | 7.0 | (20) |
| 5I2V | 20 | 70 | 6.5 | (21) |
| 2KPR | 5 | 50 | 6.8 | (22) |
| 2LOD | 10 | 50 | 6.8 | (23) |
| HIV-PRO1 | 20 | 70 | 7.0 | (24) |
| 6GZN | 15 | 70 | 7.0 | (25) |

* These oligonucleotides do not have a high-resolution NMR structure deposited in the PDB. The buffer composition was defaulted to 20 mM potassium phosphate supplemented with 70 mM KCl.

---

#### 2GKU

*Table S2. Species information*

| Sequence (5' to 3') | $\epsilon_{260nm}$ $(M^{-1}cm^{-1})$ | DOI |
|---|---|---|
| TTGGGTTAGGGTTAGGGTTAGGGA | 244300 | 10.1021/ja062791w |

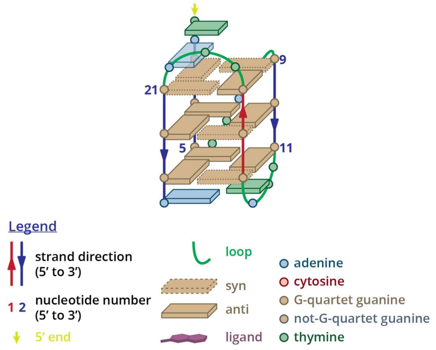

*Figure S1. Structure diagram of 2GKU*

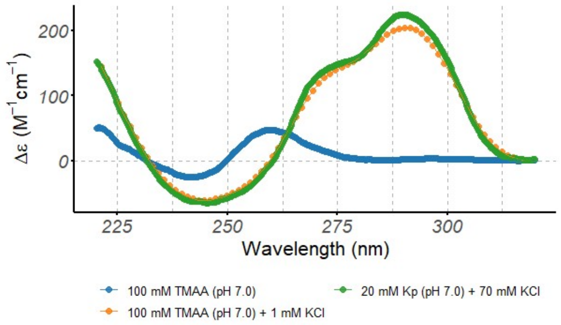

*Figure S2. Circular dichroism spectra of the 2GKU oligonucleotide (10 µM), acquired at 25°C in 0.4-cm path-length cuvettes*

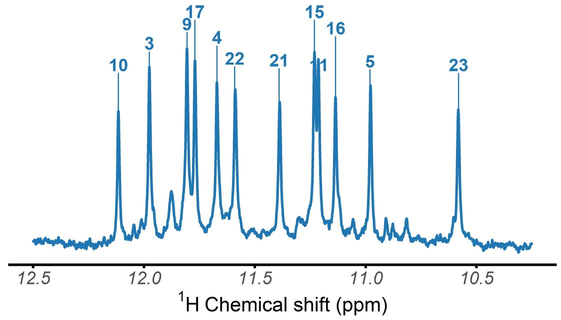

*Figure S 3. ¹H-NMR spectrum of the 2GKU oligonucleotide, acquired at 25°C in 100 mM TMAA (pH 7.0) + 1 mM KCl*

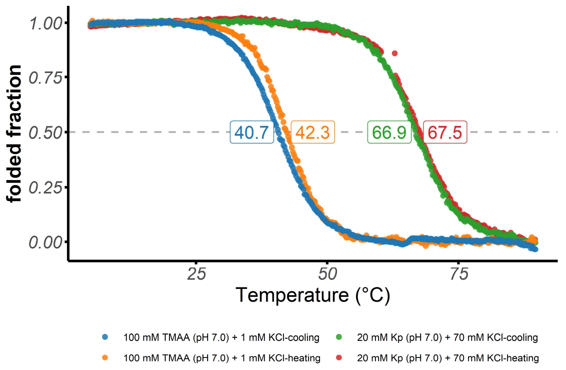

*Figure S4. Folded fraction of the 2GKU oligonucleotide as a function of temperature, determined by UV-melting (λ = 295 nm)*

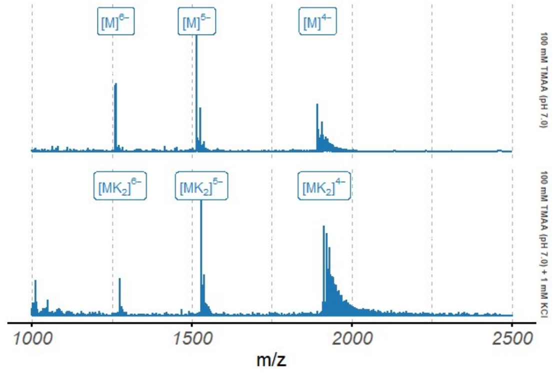

*Figure S5. Native ESI-MS spectra of the 2GKU oligonucleotide (10 µM)*

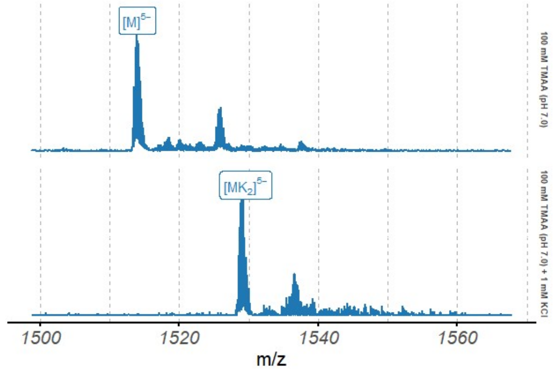

*Figure S6. Native ESI-MS spectra of the 2GKU oligonucleotide (10 µM), focused on the 5⁻ charge state*

#### 2JSM

*Table S3. Species information*

| Sequence (5' to 3') | $\epsilon_{260nm}$ $(M^{-1}cm^{-1})$ | DOI |
|---|---|---|
| TAGGGTTAGGGTTAGGGTTAGGG | 236500 | 10.1093/nar/gkm706 |

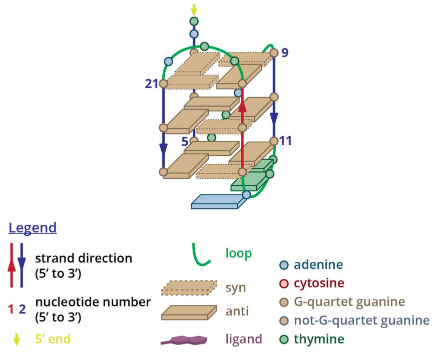

*Figure S7. Structure diagram of 2JSM*

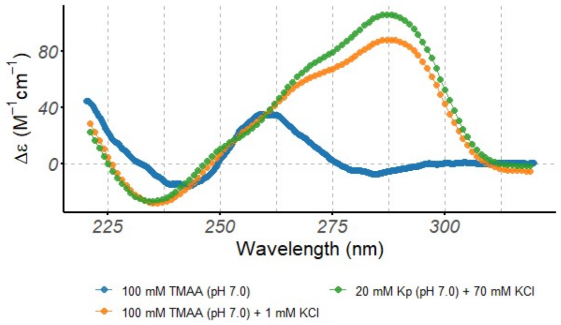

*Figure S8. Circular dichroism spectra of the 2JSM oligonucleotide (10 μM), acquired at 25°C in 0.4-cm path-length cuvettes*

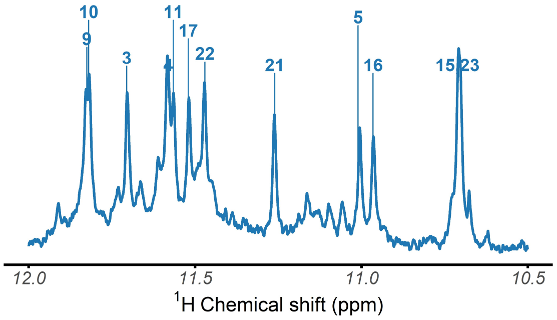

*Figure S9. $^{1}$H-NMR spectrum of the 2JSM oligonucleotide, acquired at 25°C in 100 mM TMAA (pH 7.0) + 1 mM KCl*

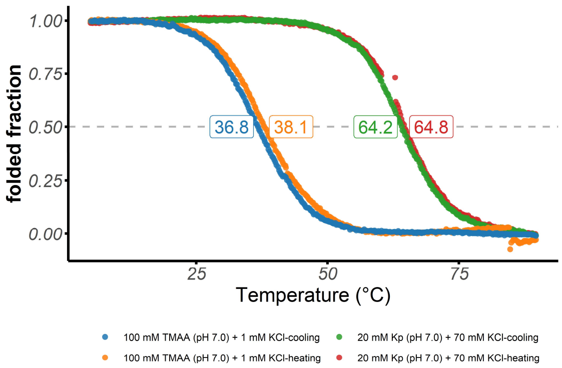

*Figure S10. Folded fraction of the 2JSM oligonucleotide as a function of temperature, determined by UV-melting (λ = 295 nm)*

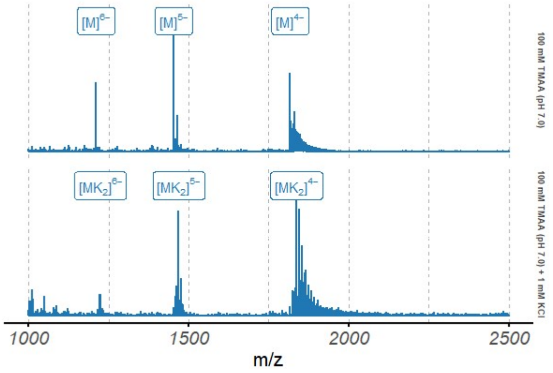

*Figure S11. Native ESI-MS spectra of the 2JSM oligonucleotide (10 μM)*

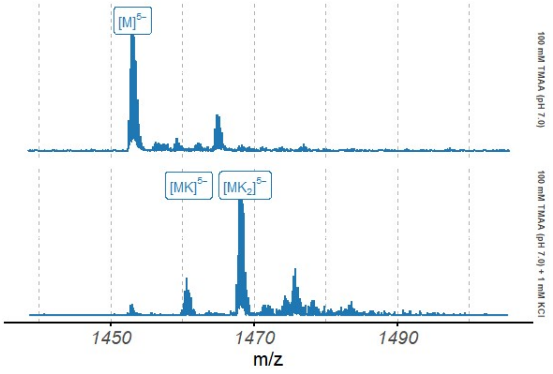

*Figure S12. Native ESI-MS spectra of the 2JSM oligonucleotide (10 μM), focused on the 5⁻ charge state*

#### 2JPZ

*Table S4. Species information*

| Sequence (5' to 3') | $\epsilon_{260nm}$ $(M^{-1}cm^{-1})$ | DOI |
|---|---|---|
| TTAGGGTTAGGGTTAGGGTTAGGGTT | 261200 | 10.1093/nar/gkm522 |

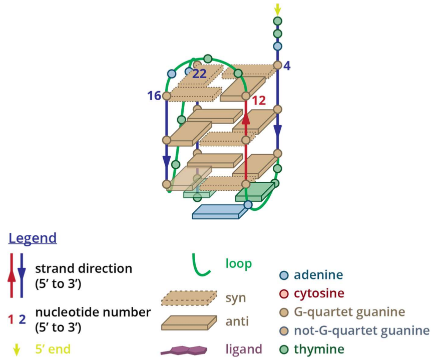

*Figure S13. Structure diagram of 2JPZ*

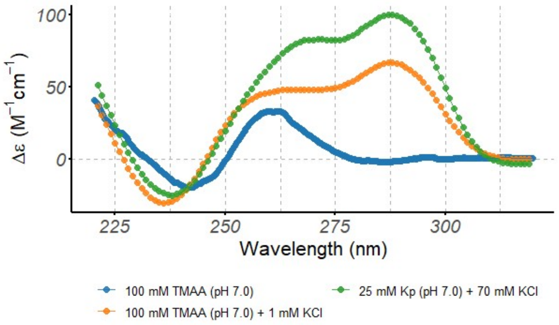

*Figure S14. Circular dichroism spectra of the 2JPZ oligonucleotide (10 µM), acquired at 25°C in 0.4-cm path-length cuvettes*

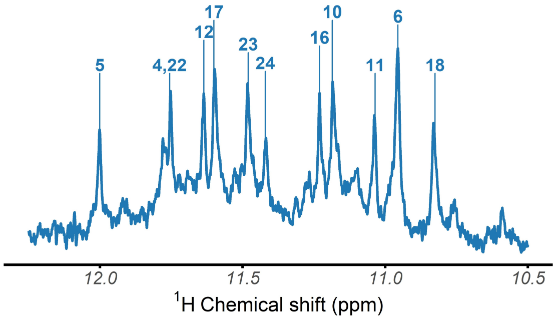

*Figure S15. $^{1}$H-NMR spectrum of the 2JPZ oligonucleotide, acquired at 25°C in 100 mM TMAA (pH 7.0) + 1 mM KCl*

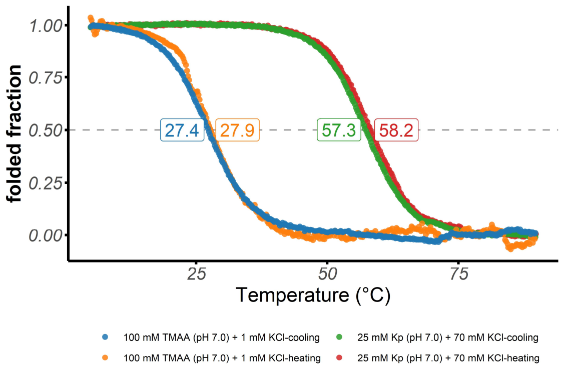

*Figure S16. Folded fraction of the 2JPZ oligonucleotide as a function of temperature, determined by UV-melting (λ = 295 nm)*

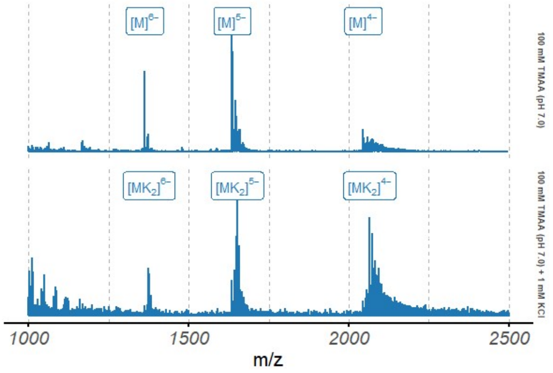

*Figure S17. Native ESI-MS spectra of the 2JPZ oligonucleotide (10 µM)*

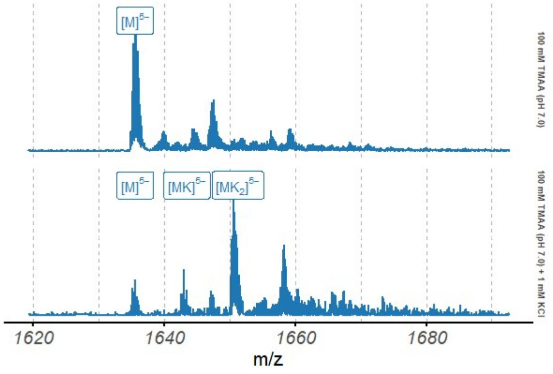

*Figure S18. Native ESI-MS spectra of the 2JPZ oligonucleotide (10 µM), focused on the 5⁻ charge state*

## 2HY9

*Table S5. Species information*

| Sequence (5' to 3') | $\epsilon_{260nm}$ $(M^{-1}cm^{-1})$ | DOI |
|---|---|---|
| AAAGGGTTAGGGTTAGGGTTAGGGAA | 278200 | 10.1093/nar/gkm009 |

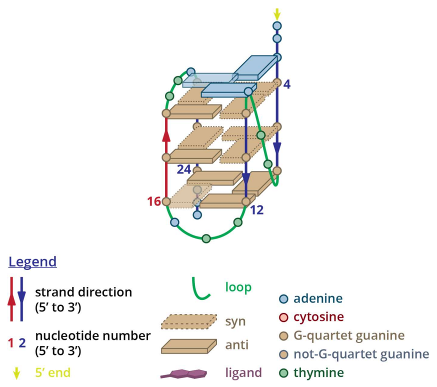

*Figure S19. Structure diagram of 2HY9*

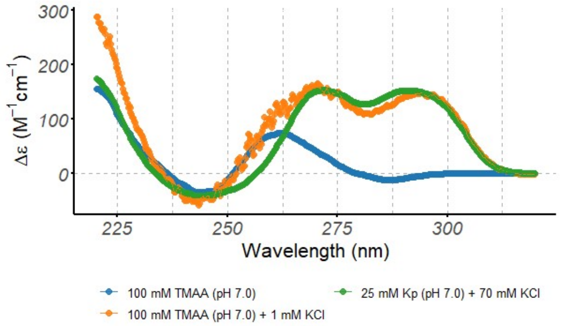

*Figure S20. Circular dichroism spectra of the 2HY9 oligonucleotide (10 μM), acquired at 25°C in 0.4-cm path-length cuvettes*

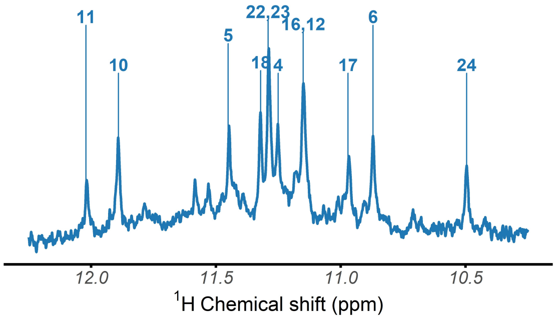

*Figure S21. $^1$H-NMR spectrum of the 2HY9 oligonucleotide, acquired at 25°C in 100 mM TMAA (pH 7.0) + 1 mM KCl*

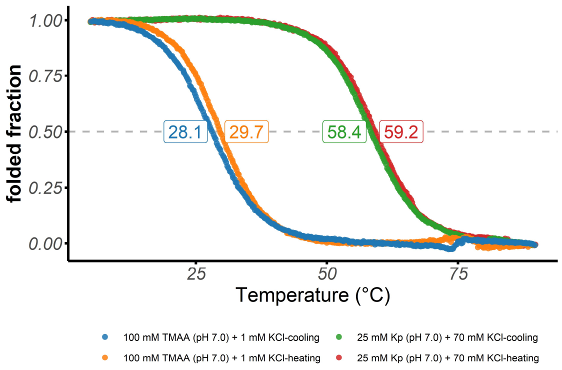

*Figure S22. Folded fraction of the 2HY9 oligonucleotide as a function of temperature, determined by UV-melting (λ = 295 nm)*

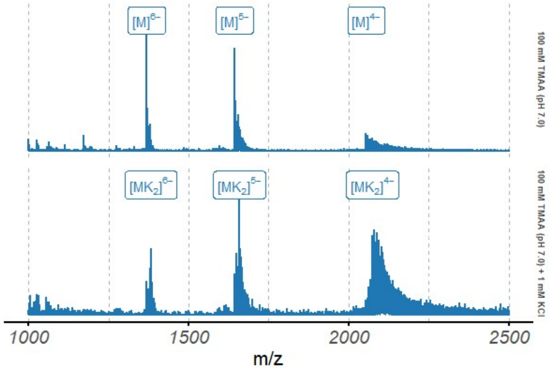

*Figure S23. Native ESI-MS spectra of the 2HY9 oligonucleotide (10 μM)*

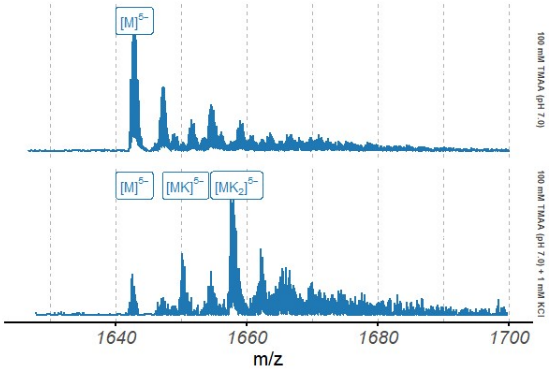

*Figure S24. Native ESI-MS spectra of the 2HY9 oligonucleotide (10 μM), focused on the 5⁻ charge state*

### 5YEY

*Table S6. Species information*

| Sequence (5' to 3') | $\epsilon_{260nm}$ $(M^{-1}cm^{-1})$ | DOI |
|---|---|---|
| GGGTTAGGGTTAGGGTTTGGG | 209100 | 10.1039/c8sc03813a |

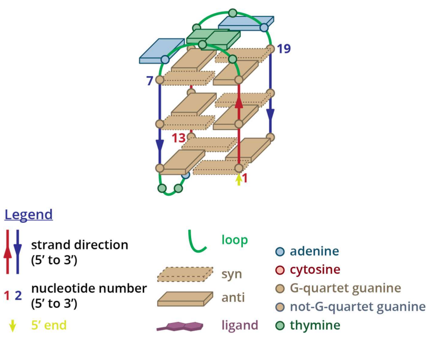

*Figure S25. Structure diagram of 5YEY*

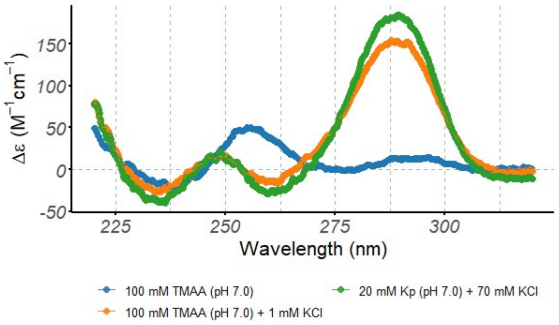

*Figure S26. Circular dichroism spectra of the 5YEY oligonucleotide (10 µM), acquired at 25°C in 0.2-cm path-length cuvettes*

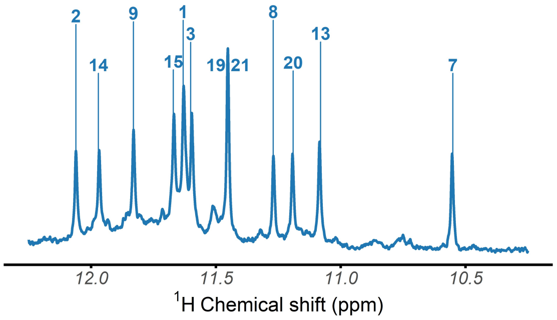

*Figure S27. $^{1}$H-NMR spectrum of the 5YEY oligonucleotide, acquired at 25°C in 100 mM TMAA (pH 7.0) + 1 mM KCl*

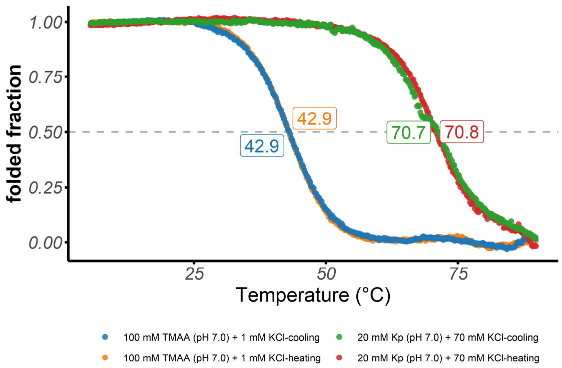

*Figure S28. Folded fraction of the 5YEY oligonucleotide as a function of temperature, determined by UV-melting (λ = 295 nm)*

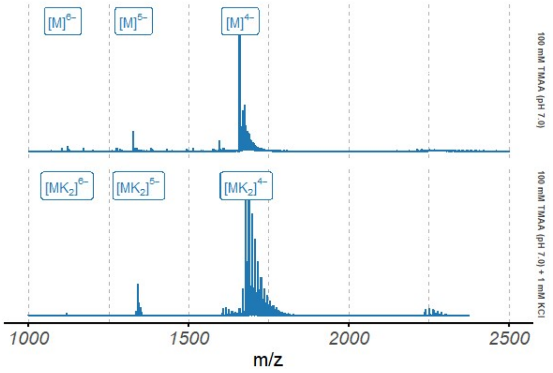

*Figure S29. Native ESI-MS spectra of the 5YEY oligonucleotide (10 µM)*

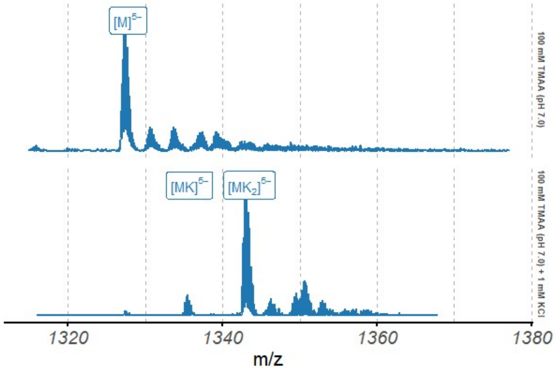

*Figure S30. Native ESI-MS spectra of the 5YEY oligonucleotide (10 µM), focused on the 5⁻ charge state*

# 2KF8

*Table S7. Species information*

| Sequence (5' to 3') | $\epsilon_{260nm}$ $(M^{-1}cm^{-1})$ | DOI |
|---|---|---|
| GGGTTAGGGTTAGGGTTAGGGT | 223500 | 10.1021/ja807503g |

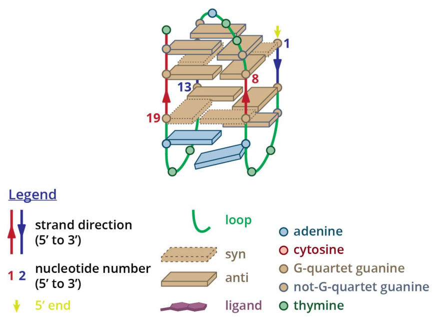

*Figure S31. Structure diagram of 2KF8*

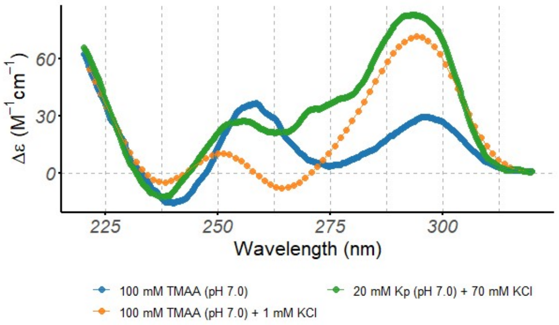

*Figure S32. Circular dichroism spectra of the 2KF8 oligonucleotide (10 µM), acquired at 25°C in 0.4-cm path-length cuvettes*

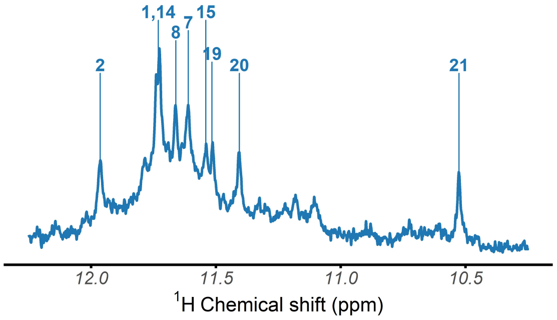

*Figure S33. $^{1}$H-NMR spectrum of the 2KF8 oligonucleotide, acquired at 25°C in 100 mM TMAA (pH 7.0) + 1 mM KCl*

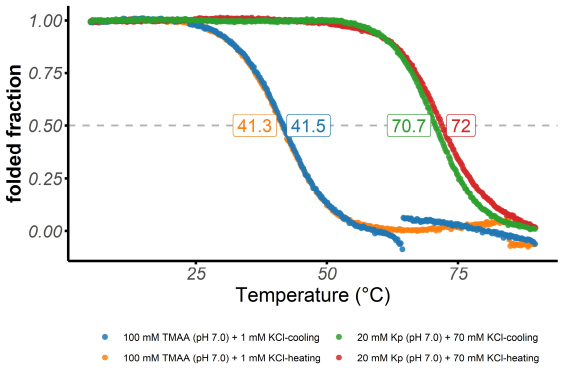

*Figure S34. Folded fraction of the 2KF8 oligonucleotide as a function of temperature, determined by UV-melting (λ = 295 nm)*

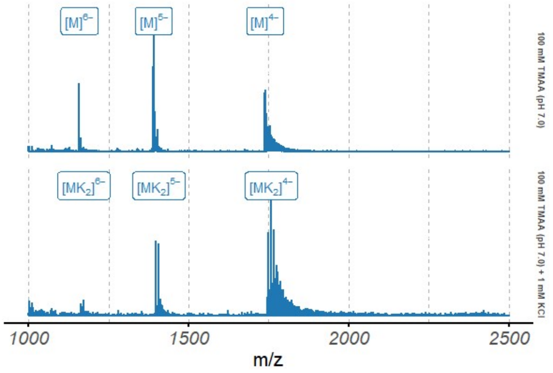

*Figure S35. Native ESI-MS spectra of the 2KF8 oligonucleotide (10 µM)*

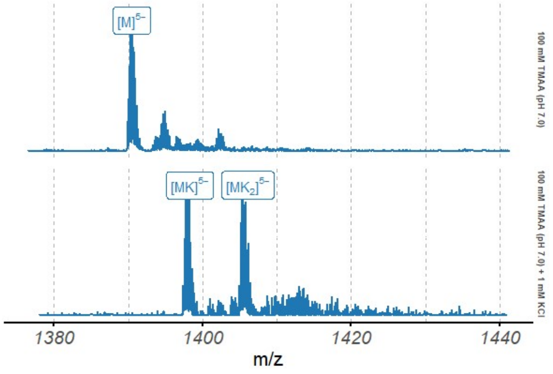

*Figure S36. Native ESI-MS spectra of the 2KF8 oligonucleotide (10 µM), focused on the $5^{-}$ charge state*

# 2KM3

*Table S8. Species information*

| Sequence (5' to 3') | $\epsilon_{260nm}$ $(M^{-1}cm^{-1})$ | DOI |
|---|---|---|
| AGGGCTAGGGCTAGGGCTAGGG | 220400 | 10.1093/nar/gkp630 |

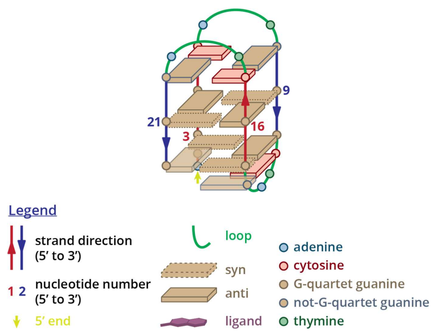

*Figure S37. Structure diagram of 2KM3*

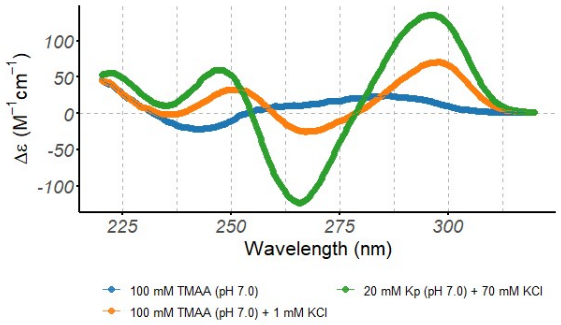

*Figure S38. Circular dichroism spectra of the 2KM3 oligonucleotide (10 μM), acquired at 25°C in 0.4-cm path-length cuvettes*

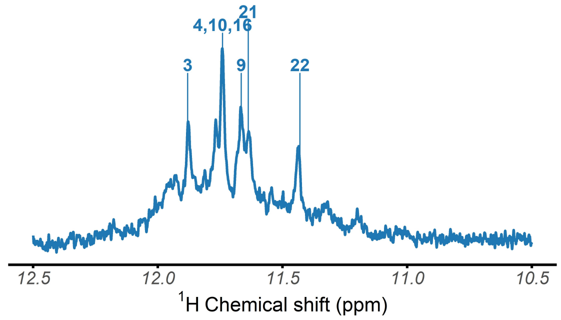

*Figure S39. $^{1}$H-NMR spectrum of the 2KM3 oligonucleotide, acquired at 25°C in 100 mM TMAA (pH 7.0) + 1 mM KCl*

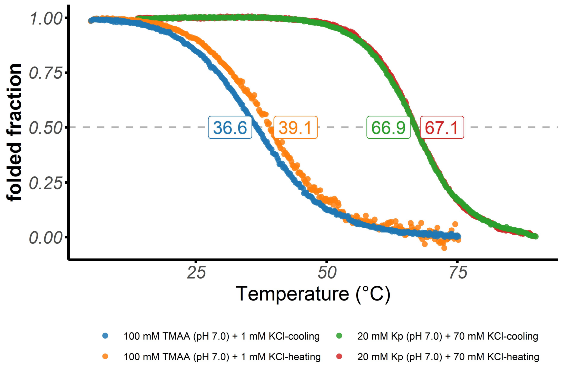

*Figure S40. Folded fraction of the 2KM3 oligonucleotide as a function of temperature, determined by UV-melting (λ = 295 nm)*

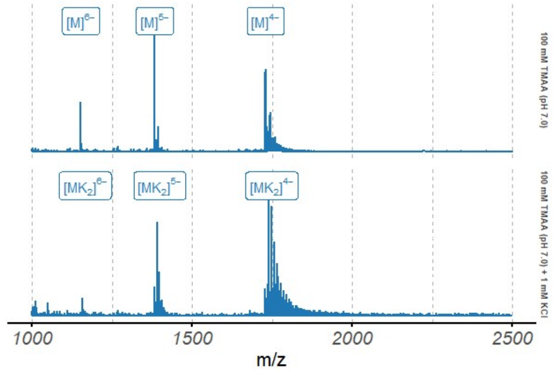

*Figure S41. Native ESI-MS spectra of the 2KM3 oligonucleotide (10 μM)*

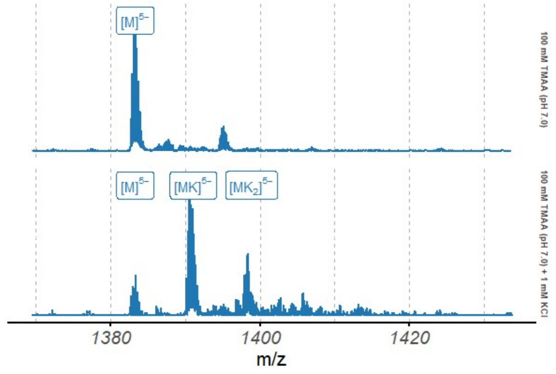

*Figure S42. Native ESI-MS spectra of the 2KM3 oligonucleotide (10 μM), focused on the 5$^{-}$ charge state*

### 5LQG

*Table S9. Species information*

| Sequence (5' to 3') | $\epsilon_{260nm}$ $(M^{-1}cm^{-1})$ | DOI |
|---|---|---|
| TAGGGTTAGGGTTAGGGTTAGG | 226400 | 10.1002/anie.201507569 |

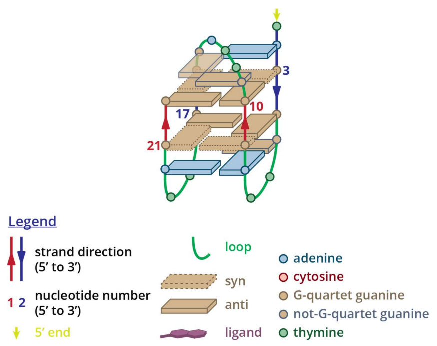

*Figure S43. Structure diagram of 5LQG*

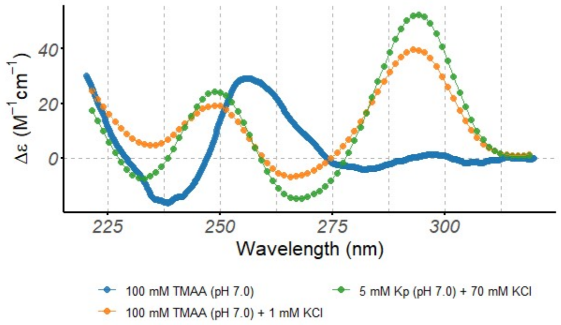

*Figure S44. Circular dichroism spectra of the 5LQG oligonucleotide (10 µM), acquired at 25°C in 0.4-cm path-length cuvettes*

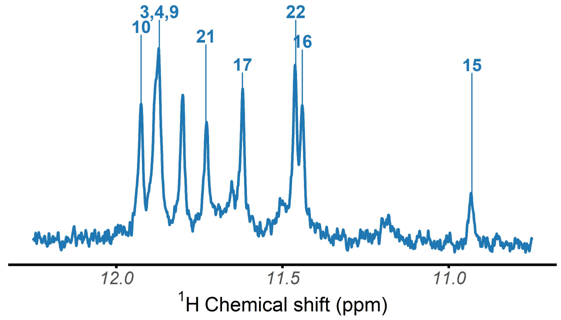

*Figure S45. $^{1}$H-NMR spectrum of the 5LQG oligonucleotide, acquired at 25°C in 100 mM TMAA (pH 7.0) + 1 mM KCl*

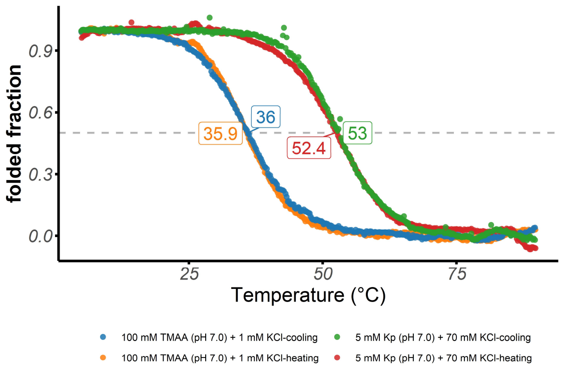

*Figure S46. Folded fraction of the 5LQG oligonucleotide as a function of temperature, determined by UV-melting (λ = 295 nm)*

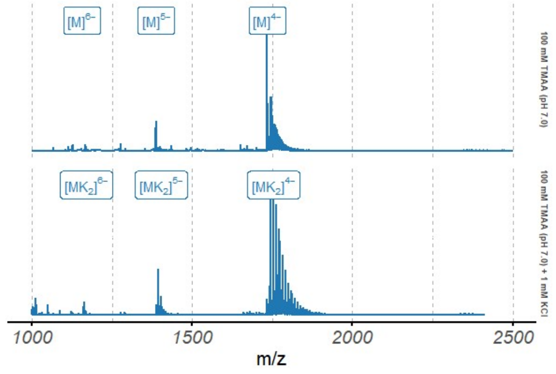

*Figure S47. Native ESI-MS spectra of the 5LQG oligonucleotide (10 µM)*

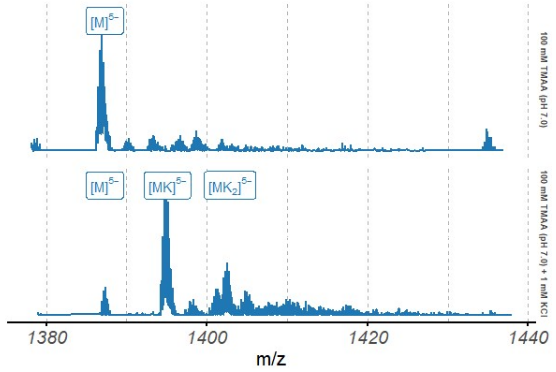

*Figure S48. Native ESI-MS spectra of the 5LQG oligonucleotide (10 µM), focused on the 5⁻ charge state*

# 21G

| Sequence (5' to 3') | $\epsilon_{260nm}$ $(M^{-1}cm^{-1})$ | DOI |
|---|---|---|
| GGGTTAGGGTTAGGGTTAGGG | 215000 | 10.1021/jm301899y |

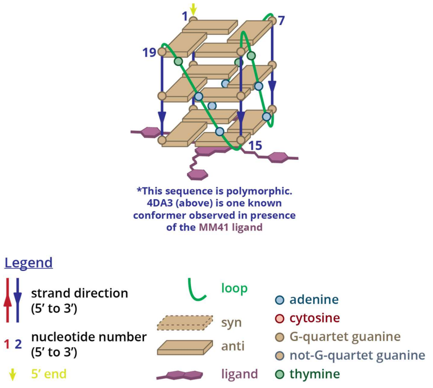

*Figure S49. Structure diagram of 21G*

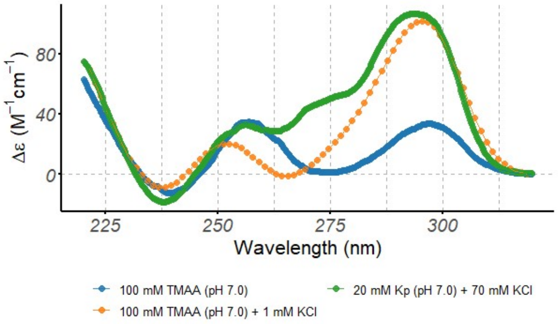

*Figure S50. Circular dichroism spectra of the 21G oligonucleotide (10 µM), acquired at 25°C in 0.4-cm path-length cuvettes*

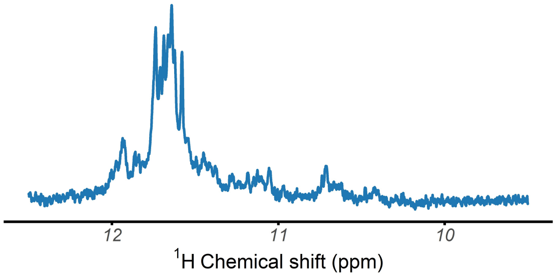

*Figure S51. $^{1}$H-NMR spectrum of the 21G oligonucleotide, acquired at 25°C in 100 mM TMAA (pH 7.0) + 1 mM KCl*

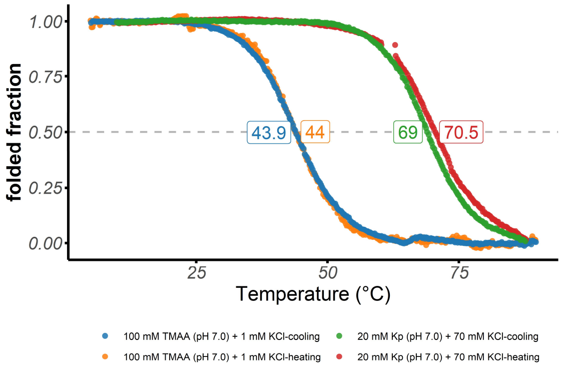

*Figure S52. Folded fraction of the 21G oligonucleotide as a function of temperature, determined by UV-melting (λ = 295 nm)*

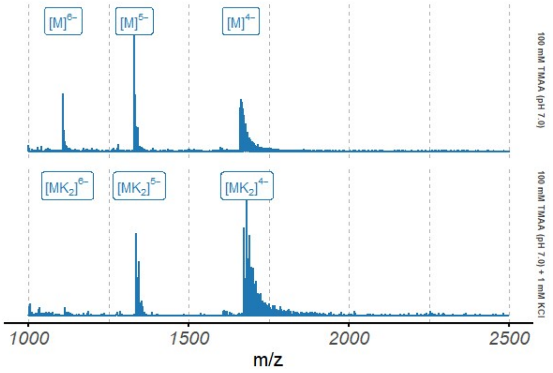

*Figure S53. Native ESI-MS spectra of the 21G oligonucleotide (10 µM)*

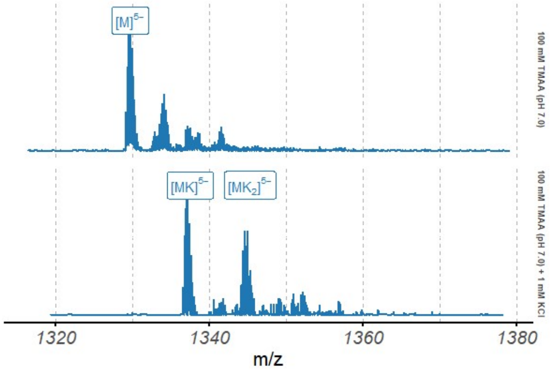

*Figure S54. Native ESI-MS spectra of the 21G oligonucleotide (10 µM), focused on the $5^-$ charge state*

## 22AG

| Sequence (5' to 3') | $\epsilon_{260nm}$ $(M^{-1}cm^{-1})$ | DOI |
|---|---|---|
| AGGGTTAGGGTTAGGGTTAGGG | 228500 | 10.1016/0969-2126(93)90015-9, 10.1038/nature755 |

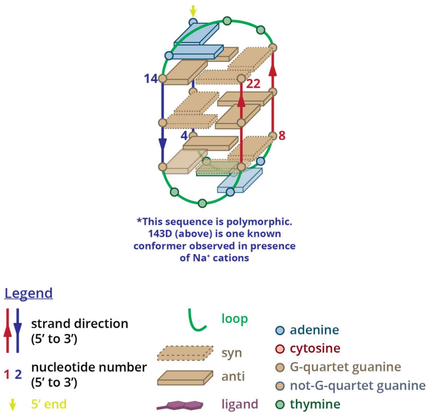

Figure S55. Structure diagram of 22AG

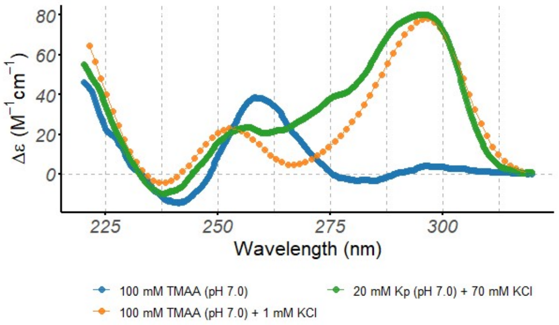

*Figure S56. Circular dichroism spectra of the 22AG oligonucleotide (10 μM), acquired at 25°C in 0.4-cm path-length cuvettes*

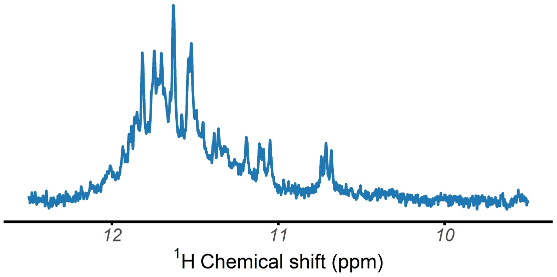

*Figure S57. ¹H-NMR spectrum of the 22AG oligonucleotide, acquired at 25°C in 100 mM TMAA (pH 7.0) + 1 mM KCl*

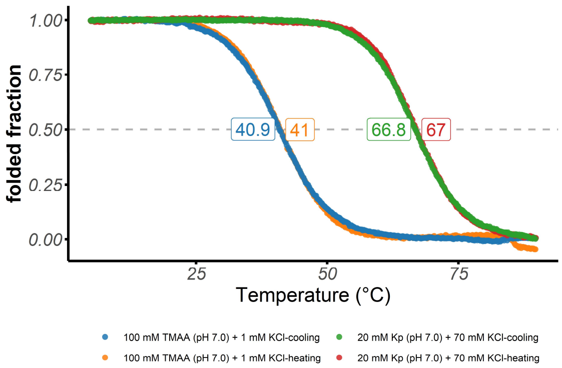

*Figure S58. Folded fraction of the 22AG oligonucleotide as a function of temperature, determined by UV-melting (λ = 295 nm)*

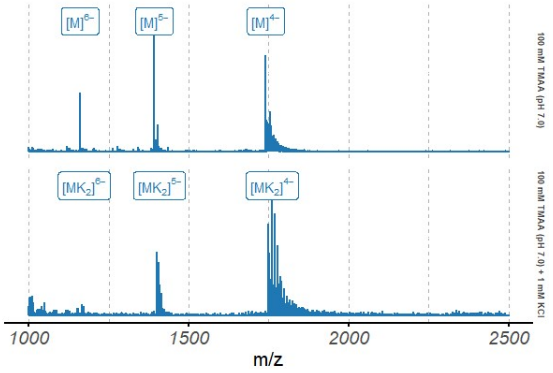

*Figure S59. Native ESI-MS spectra of the 22AG oligonucleotide (10 μM)*

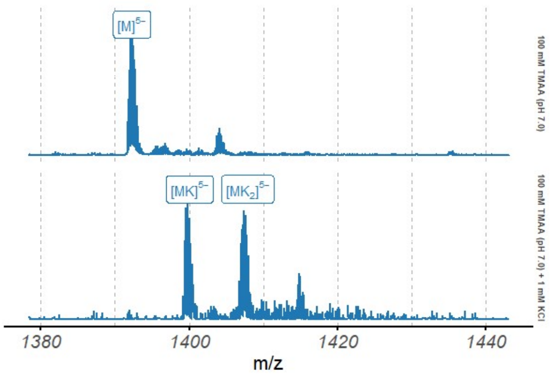

*Figure S60. Native ESI-MS spectra of the 22AG oligonucleotide (10 μM), focused on the 5⁻ charge state*

## 2LK7

*Table S10. Species information*

| Sequence (5' to 3') | $\epsilon_{260nm}$ $(M^{-1}cm^{-1})$ | DOI |
|---|---|---|
| TTGGGTGGGTGGGTGGGT | 172800 | 10.1002/chem.201103295 |

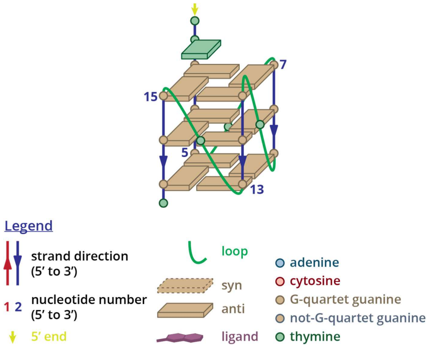

*Figure S61. Structure diagram of 2LK7*

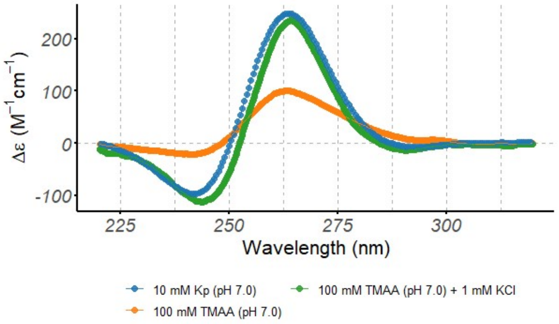

*Figure S62. Circular dichroism spectra of the 2LK7 oligonucleotide (10 µM), acquired at 25°C in 0.4-cm path-length cuvettes*

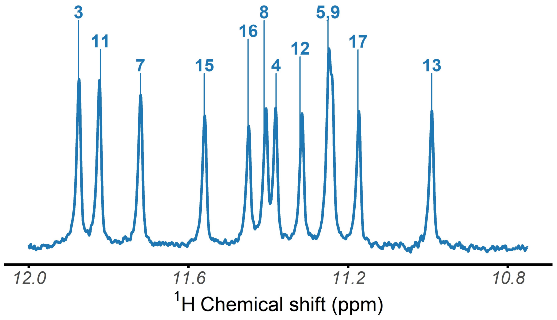

*Figure S63. $^{1}$H-NMR spectrum of the 2LK7 oligonucleotide, acquired at 25°C in 100 mM TMAA (pH 7.0) + 1 mM KCl*

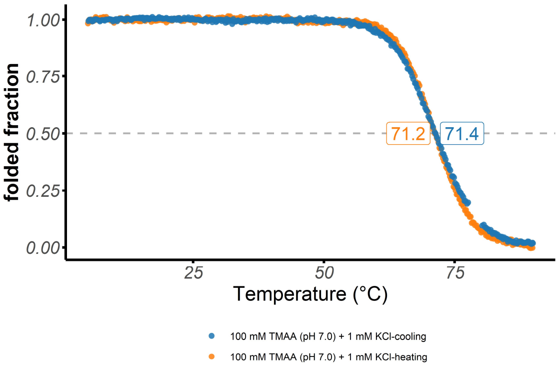

*Figure S64. Folded fraction of the 2LK7 oligonucleotide as a function of temperature, determined by UV-melting (λ = 295 nm)*

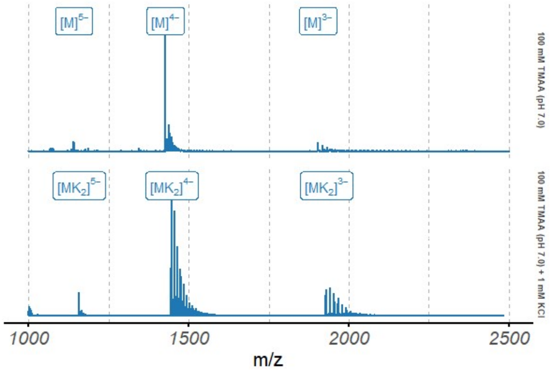

*Figure S65. Native ESI-MS spectra of the 2LK7 oligonucleotide (10 µM)*

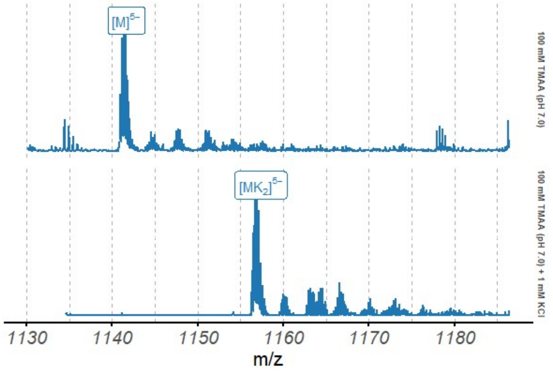

*Figure S66. Native ESI-MS spectra of the 2LK7 oligonucleotide (10 µM), focused on the 5⁻ charge state*

### 2LEE

*Table S11. Species information*

| Sequence (5' to 3') | $\epsilon_{260nm}$ $(M^{-1}cm^{-1})$ | DOI |
|---|---|---|
| TAGGGCGGGAGGGAGGGAA | 202800 | 10.1021/ja208483v |

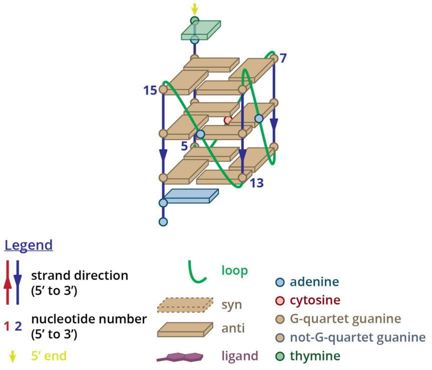

*Figure S67. Structure diagram of 2LEE*

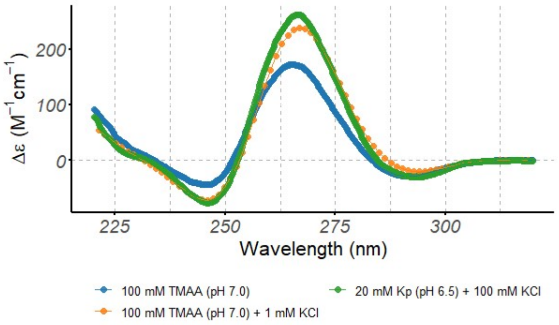

*Figure S68. Circular dichroism spectra of the 2LEE oligonucleotide (10 µM), acquired at 25°C in 0.4-cm path-length cuvettes*

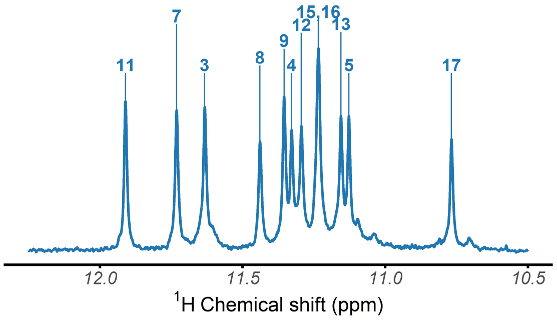

*Figure S69. $^1$H-NMR spectrum of the 2LEE oligonucleotide, acquired at 25°C in 100 mM TMAA (pH 7.0) + 1 mM KCl*

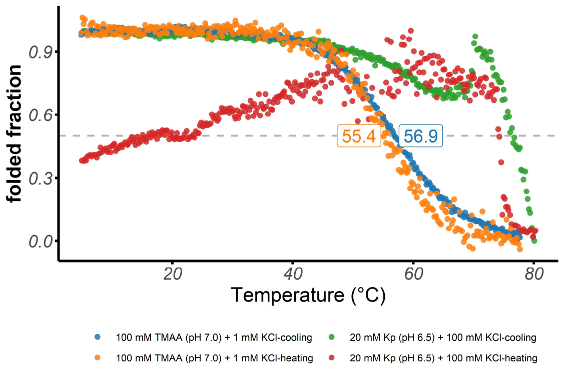

*Figure S70. Folded fraction of the 2LEE oligonucleotide as a function of temperature, determined by UV-melting (λ = 295 nm)*

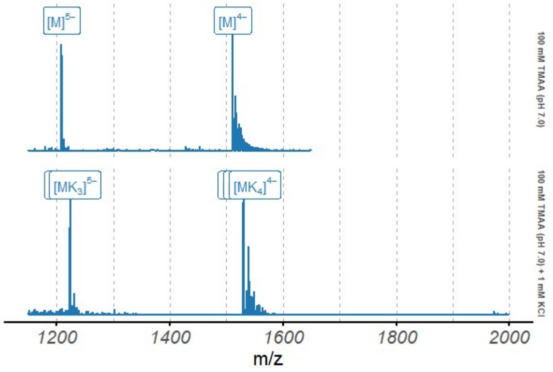

*Figure S71. Native ESI-MS spectra of the 2LEE oligonucleotide (10 µM)*

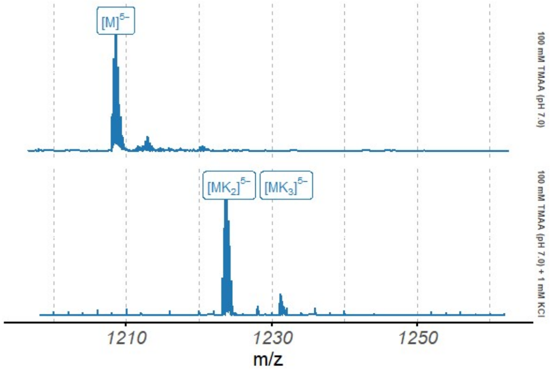

*Figure S72. Native ESI-MS spectra of the 2LEE oligonucleotide (10 µM), focused on the $5^-$ charge state*

# 2M4P

*Table S12. Species information*

| Sequence (5' to 3') | $\epsilon_{260nm}$ $(M^{-1}cm^{-1})$ | DOI |
|---|---|---|
| TTGTGGTGGGTGGGTGGGT | 181500 | 10.1038/nature755 |

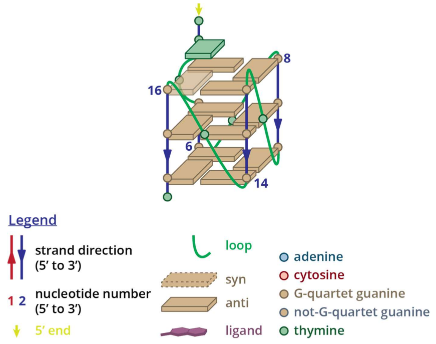

*Figure S73. Structure diagram of 2M4P*

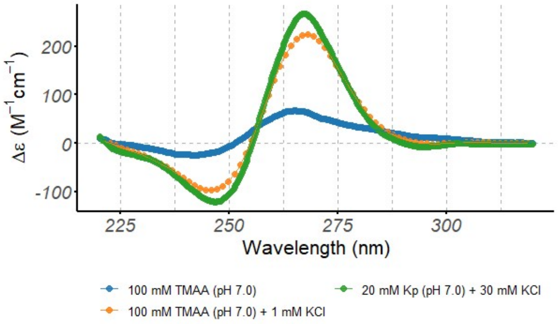

*Figure S74. Circular dichroism spectra of the 2M4P oligonucleotide (10 µM), acquired at 25°C in 0.4-cm path-length cuvettes*

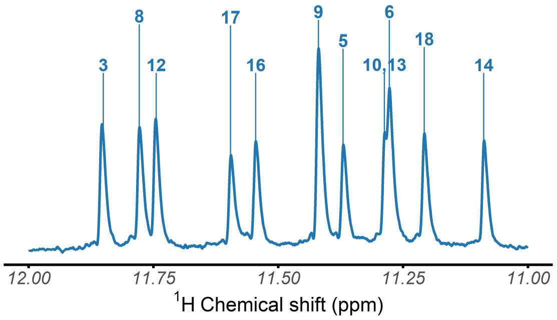

*Figure S75. $^{1}$H-NMR spectrum of the 2M4P oligonucleotide, acquired at 25°C in 100 mM TMAA (pH 7.0) + 1 mM KCl*

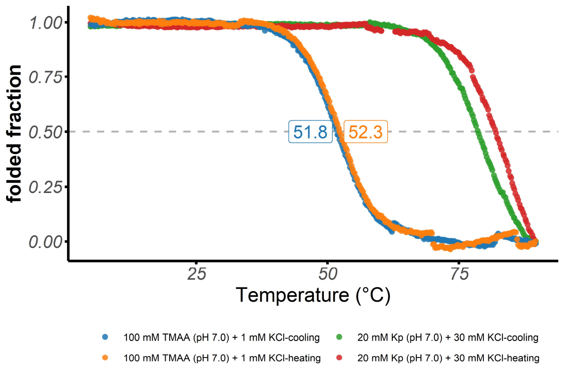

*Figure S76. Folded fraction of the 2M4P oligonucleotide as a function of temperature, determined by UV-melting (λ = 295 nm)*

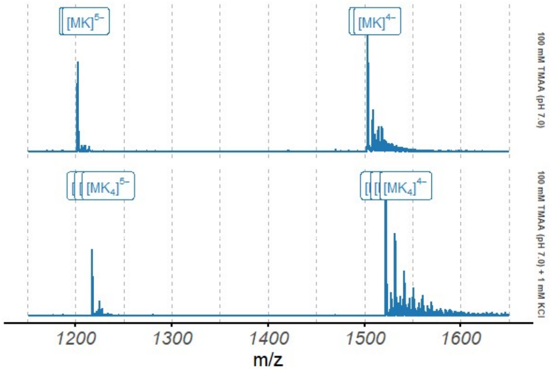

*Figure S77. Native ESI-MS spectra of the 2M4P oligonucleotide (10 µM)*

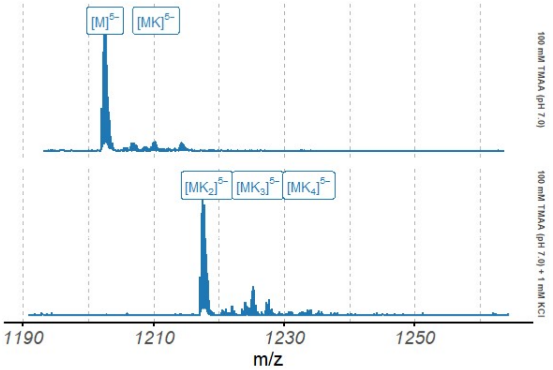

*Figure S78. Native ESI-MS spectra of the 2M4P oligonucleotide (10 µM), focused on the 5⁻ charge state*

#### 2LXQ

*Table S13. Species information*

| Sequence (5' to 3') | $\epsilon_{260nm}$ $(M^{-1}cm^{-1})$ | DOI |
|---|---|---|
| TAGGGTGGGTTGGGTGGGGAAT | 221500 | 10.1016/j.str.2012.09.013 |

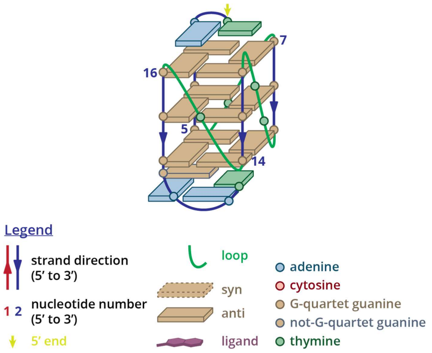

*Figure S79. Structure diagram of 2LXQ*

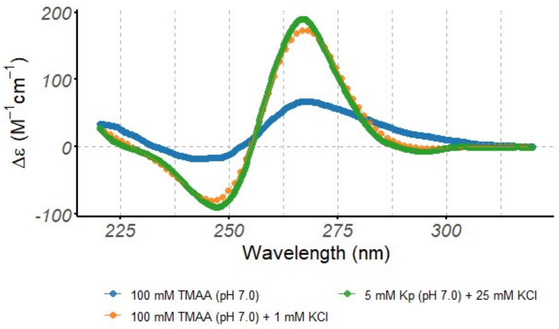

*Figure S80. Circular dichroism spectra of the 2LXQ oligonucleotide (10 μM), acquired at 25°C in 0.4-cm path-length cuvettes*

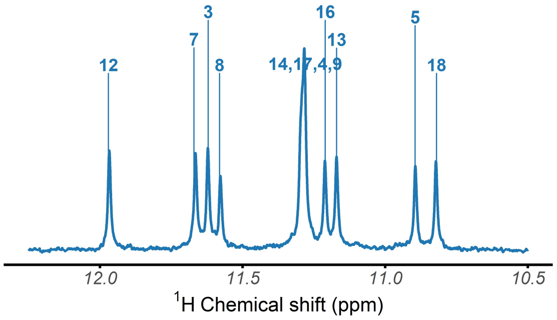

*Figure S81. ¹H-NMR spectrum of the 2LXQ oligonucleotide, acquired at 25°C in 100 mM TMAA (pH 7.0) + 1 mM KCl*

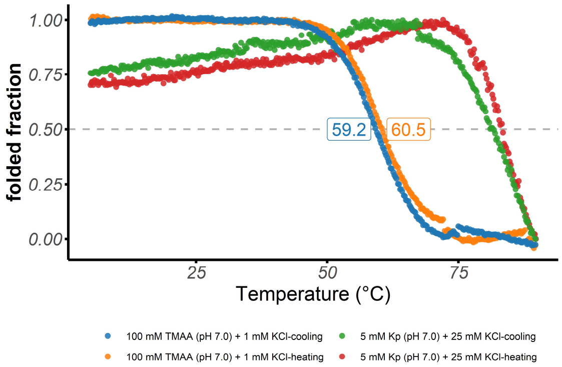

*Figure S82. Folded fraction of the 2LXQ oligonucleotide as a function of temperature, determined by UV-melting (λ = 295 nm)*

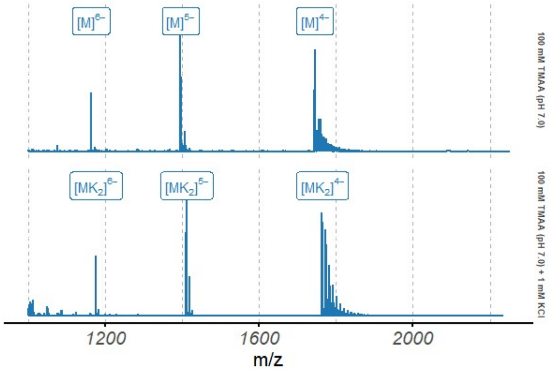

*Figure S83. Native ESI-MS spectra of the 2LXQ oligonucleotide (10 μM)*

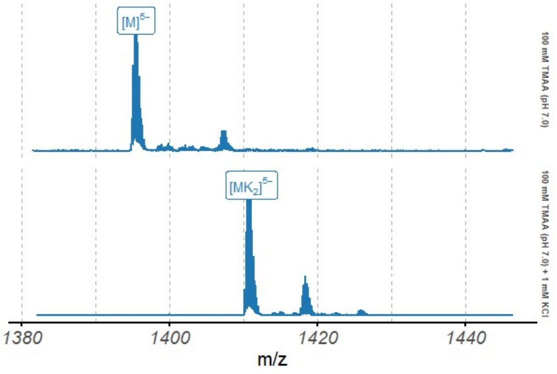

*Figure S84. Native ESI-MS spectra of the 2LXQ oligonucleotide (10 μM), focused on the 5⁻ charge state*

## 2M27

*Table S14. Species information*

| Sequence (5' to 3') | $\epsilon_{260nm}$ $(M^{-1}cm^{-1})$ | DOI |
|---|---|---|
| CGGGGCGGGCCTTGGGCGGGGT | 200400 | 10.1093/nar/gkt784 |

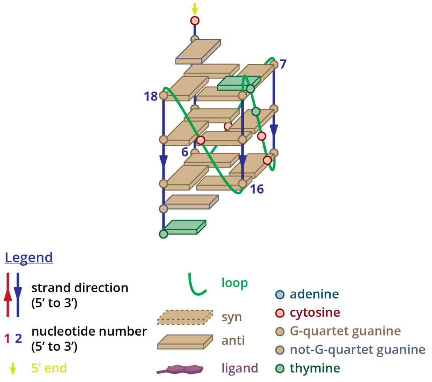

*Figure S85. Structure diagram of 2M27*

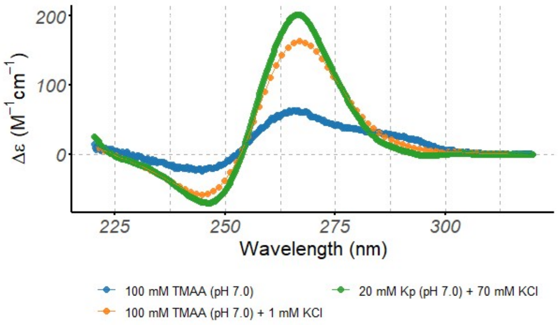

*Figure S86. Circular dichroism spectra of the 2M27 oligonucleotide (10 μM), acquired at 25°C in 0.4-cm path-length cuvettes*

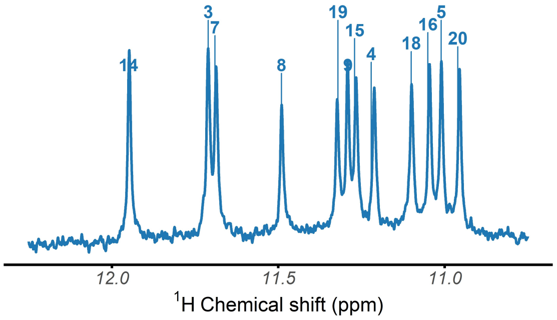

*Figure S87. $^1$H-NMR spectrum of the 2M27 oligonucleotide, acquired at 25°C in 100 mM TMAA (pH 7.0) + 1 mM KCl*

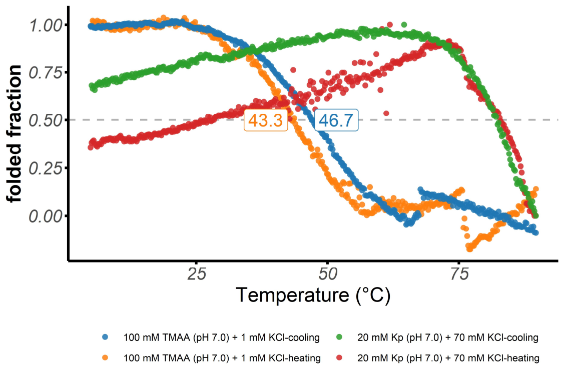

*Figure S88. Folded fraction of the 2M27 oligonucleotide as a function of temperature, determined by UV-melting (λ = 295 nm)*

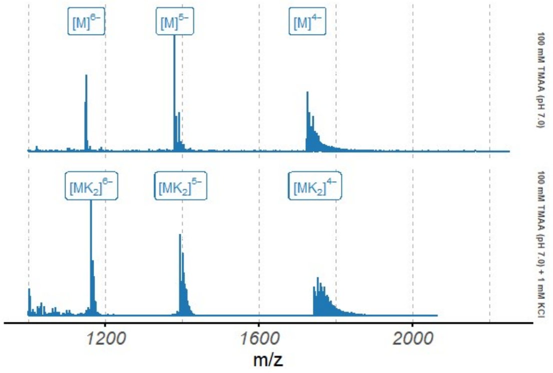

*Figure S89. Native ESI-MS spectra of the 2M27 oligonucleotide (10 μM)*

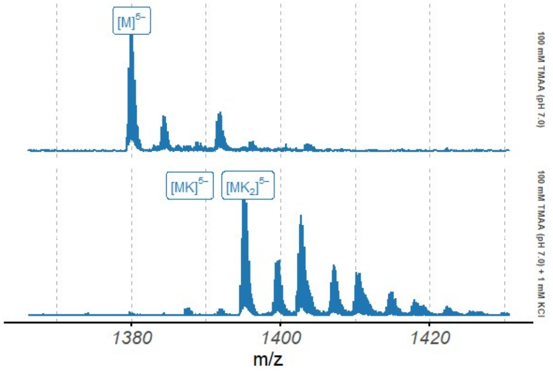

*Figure S90. Native ESI-MS spectra of the 2M27 oligonucleotide (10 μM), focused on the 5⁻ charge state*

#### 1XAV

*Table S15. Species information*

| Sequence (5' to 3') | $\epsilon_{260nm}$ $(M^{-1}cm^{-1})$ | DOI |
|---|---|---|
| TGAGGGTGGGTAGGGTGGGTAA | 228700 | 10.1021/bi048242p |

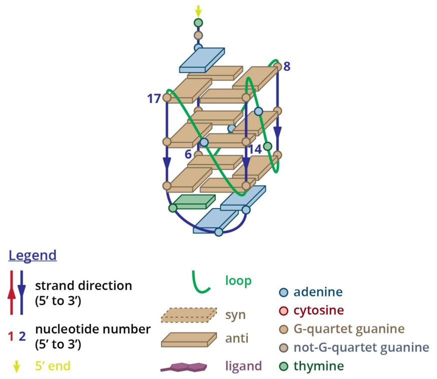

*Figure S91. Structure diagram of 1XAV*

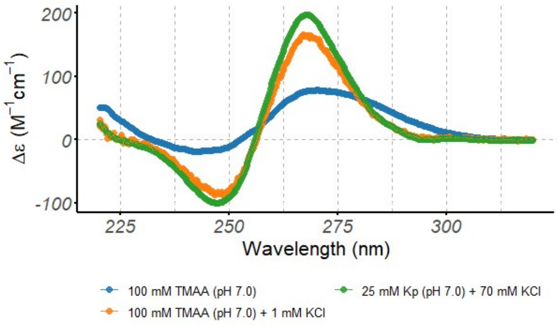

*Figure S92. Circular dichroism spectra of the 1XAV oligonucleotide (10 µM), acquired at 25°C in 0.4-cm path-length cuvettes*

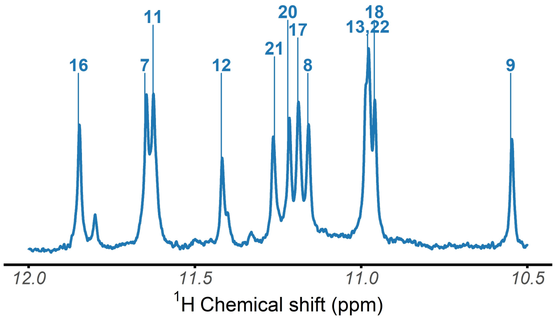

*Figure S93. $^{1}$H-NMR spectrum of the 1XAV oligonucleotide, acquired at 25°C in 100 mM TMAA (pH 7.0) + 1 mM KCl*

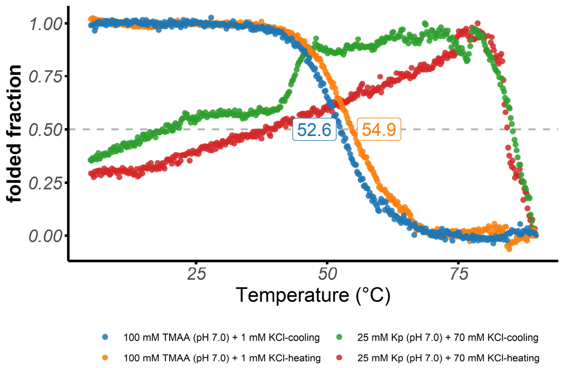

*Figure S94. Folded fraction of the 1XAV oligonucleotide as a function of temperature, determined by UV-melting (λ = 295 nm)*

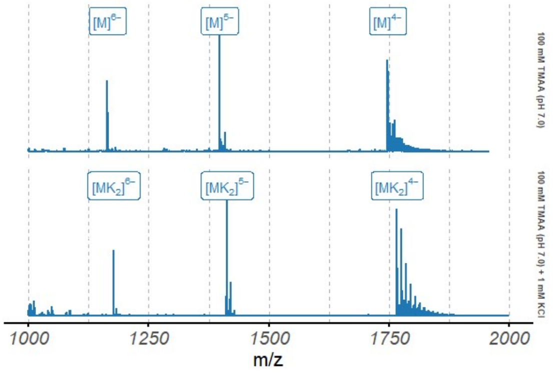

*Figure S95. Native ESI-MS spectra of the 1XAV oligonucleotide (10 µM)*

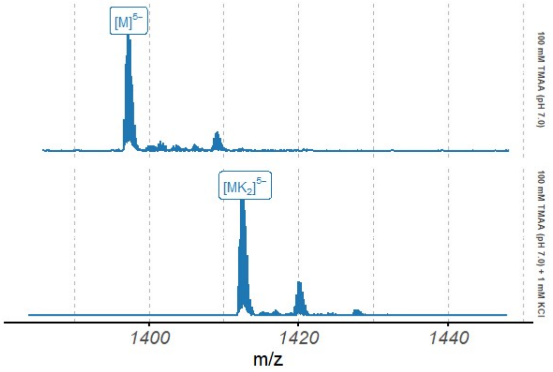

*Figure S96. Native ESI-MS spectra of the 1XAV oligonucleotide (10 µM), focused on the 5⁻ charge state*

### 2MGN

*Table S16. Species information*

| Sequence (5' to 3') | $\epsilon_{260nm}$ $(M^{-1}cm^{-1})$ | DOI |
|---|---|---|
| TGAGGGTGGTGAGGGTGGGGAAGG | 248200 | 10.1002/anie.201308063 |

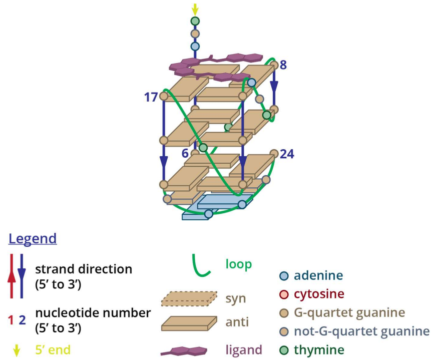

*Figure S97. Structure diagram of 2MGN*

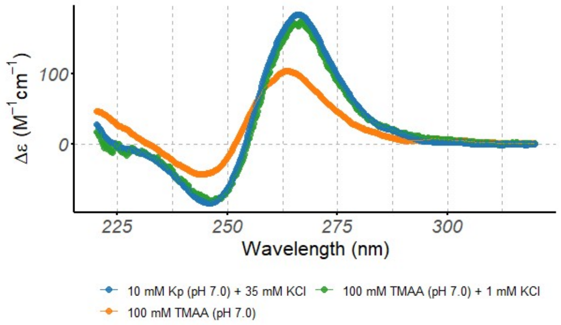

*Figure S98. Circular dichroism spectra of the 2MGN oligonucleotide (10 μM), acquired at 25°C in 0.4-cm path-length cuvettes*

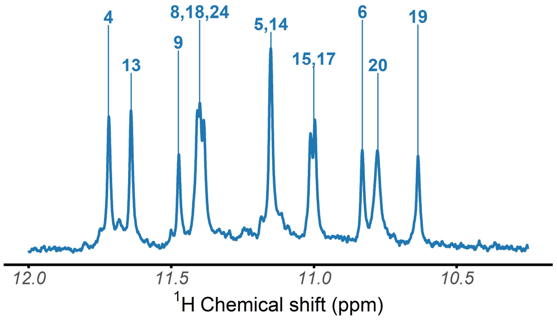

*Figure S99. ¹H-NMR spectrum of the 2MGN oligonucleotide, acquired at 25°C in 100 mM TMAA (pH 7.0) + 1 mM KCl*

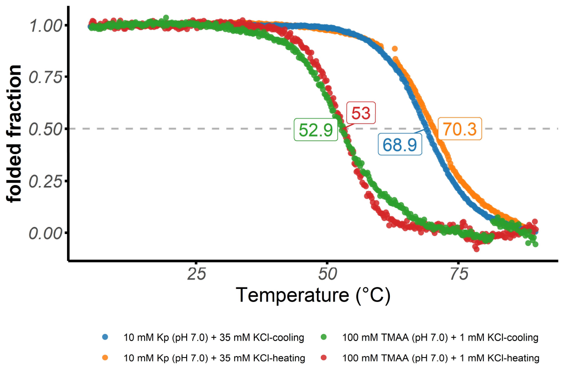

*Figure S100. Folded fraction of the 2MGN oligonucleotide as a function of temperature, determined by UV-melting (λ = 295 nm)*

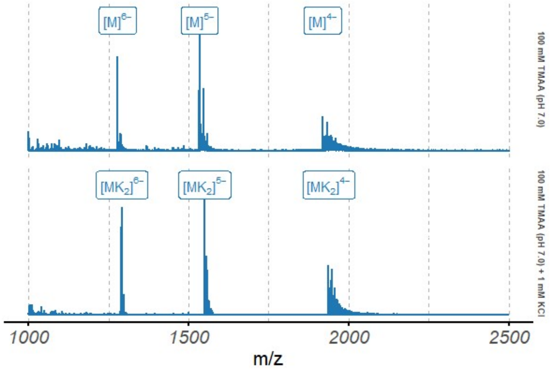

*Figure S101. Native ESI-MS spectra of the 2MGN oligonucleotide (10 μM)*

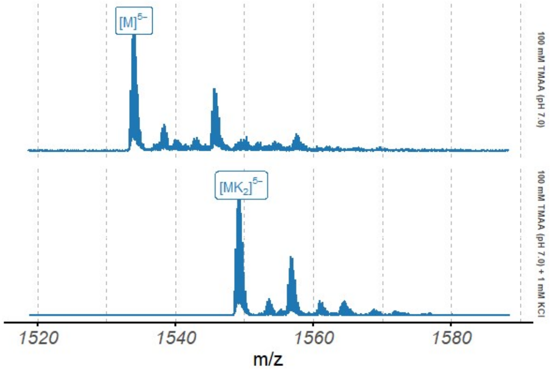

*Figure S102. Native ESI-MS spectra of the 2MGN oligonucleotide (10 μM), focused on the 5⁻ charge state*

### 2LPW

*Table S17. Species information*

| Sequence (5' to 3') | $\epsilon_{260nm}$ $(M^{-1}cm^{-1})$ | DOI |
|---|---|---|
| AAGGGTGGGTGTAAGTGTGGGTGGGT | 265100 | 10.1021/ja208993r |

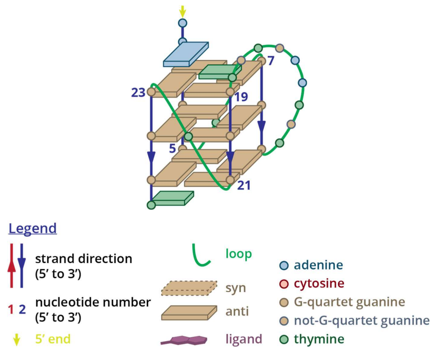

*Figure S103. Structure diagram of 2LPW*

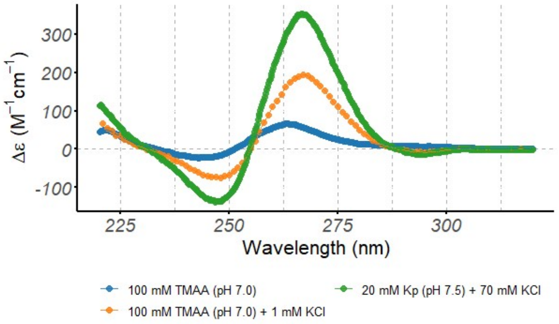

*Figure S104. Circular dichroism spectra of the 2LPW oligonucleotide (10 µM), acquired at 25°C in 0.4-cm path-length cuvettes*

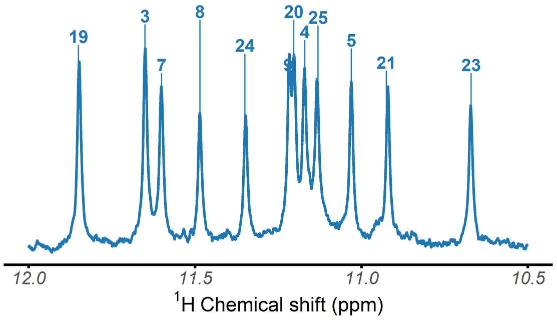

*Figure S105. $^{1}$H-NMR spectrum of the 2LPW oligonucleotide, acquired at 25°C in 100 mM TMAA (pH 7.0) + 1 mM KCl*

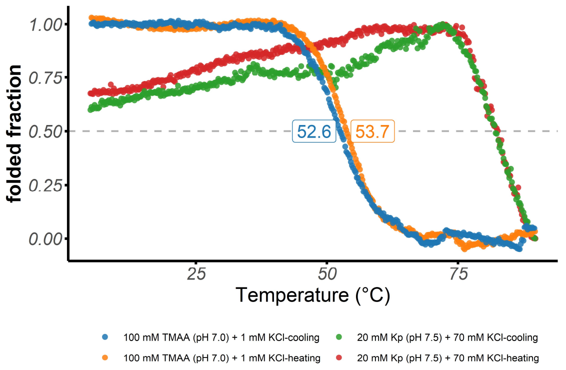

*Figure S106. Folded fraction of the 2LPW oligonucleotide as a function of temperature, determined by UV-melting (λ = 295 nm)*

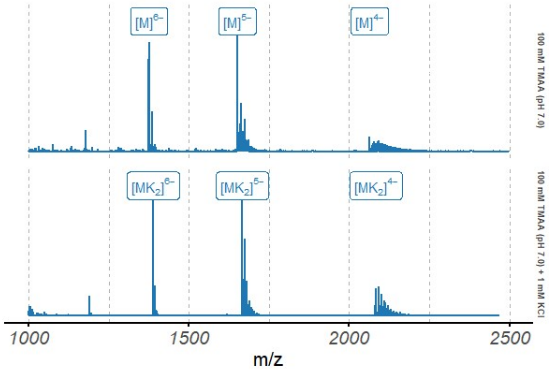

*Figure S107. Native ESI-MS spectra of the 2LPW oligonucleotide (10 µM)*

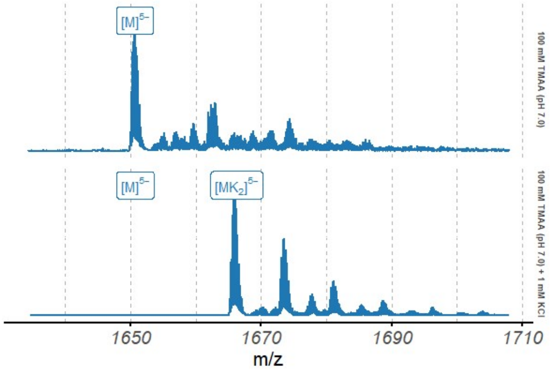

*Figure S108. Native ESI-MS spectra of the 2LPW oligonucleotide (10 µM), focused on the 5⁻ charge state*

#### 2LBY

*Table S18. Species information*

| Sequence (5' to 3') | $\epsilon_{260nm}$ $(M^{-1}cm^{-1})$ | DOI |
|---|---|---|
| TAGGGAGGGTAGGGAGGGT | 201700 | 10.1093/nar/gkr612 |

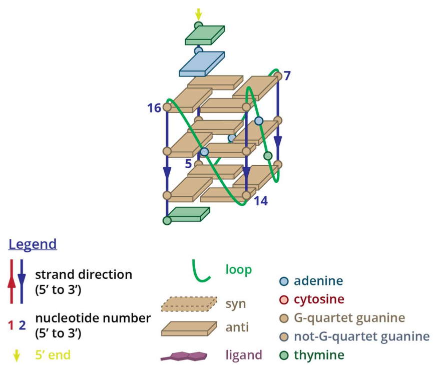

*Figure S109. Structure diagram of 2LBY*

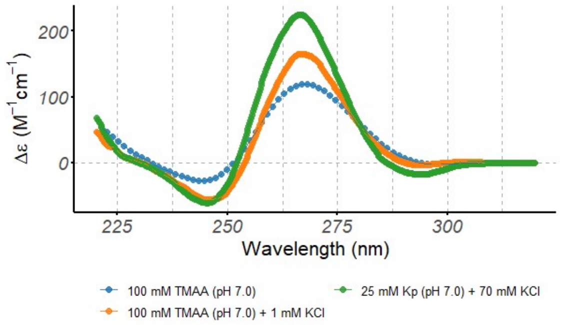

*Figure S110. Circular dichroism spectra of the 2LBY oligonucleotide (10 µM), acquired at 25°C in 0.4-cm path-length cuvettes*

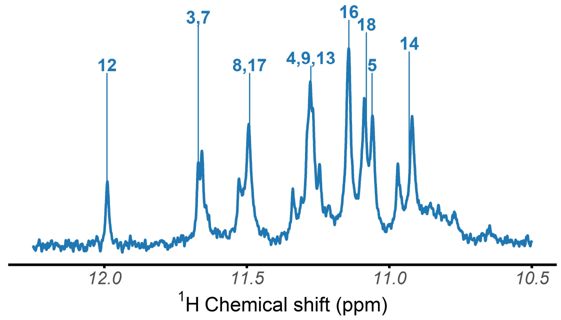

*Figure S111. ¹H-NMR spectrum of the 2LBY oligonucleotide, acquired at 25°C in 100 mM TMAA (pH 7.0) + 1 mM KCl*

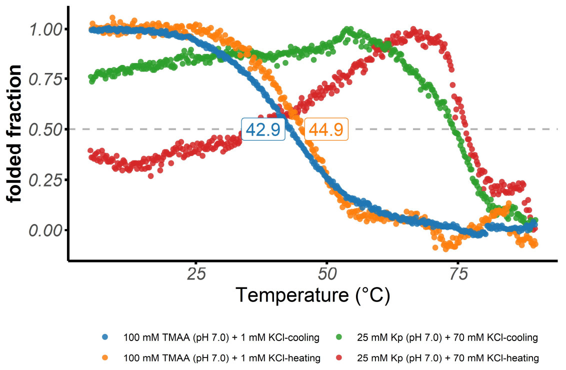

*Figure S112. Folded fraction of the 2LBY oligonucleotide as a function of temperature, determined by UV-melting (λ = 295 nm)*

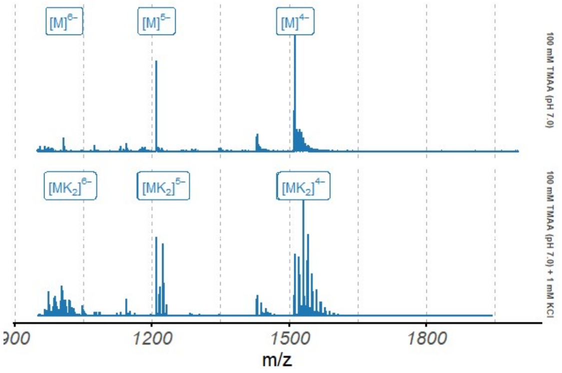

*Figure S113. Native ESI-MS spectra of the 2LBY oligonucleotide (10 µM)*

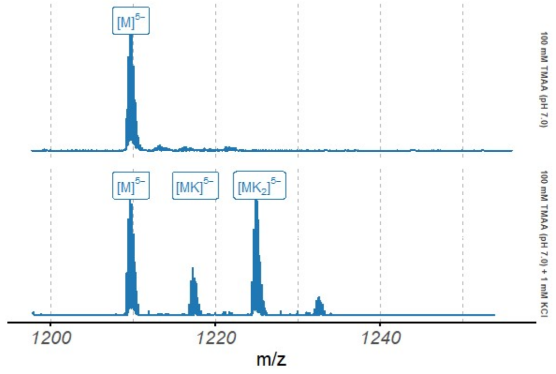

*Figure S114. Native ESI-MS spectra of the 2LBY oligonucleotide (10 µM), focused on the 5⁻ charge state*

# 2O3M

*Table S19. Species information*

| Sequence (5' to 3') | $\epsilon_{260nm}$ $(M^{-1}cm^{-1})$ | DOI |
|---|---|---|
| AGGGAGGGCGCTGGGAGGAGGG | 226700 | 10.1021/ja068739h |

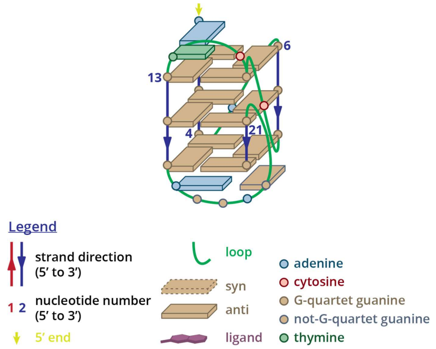

*Figure S115. Structure diagram of 2O3M*

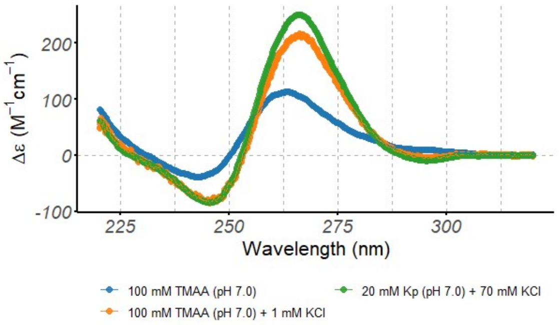

*Figure S116. Circular dichroism spectra of the 2O3M oligonucleotide (10 µM), acquired at 25°C in 0.4-cm path-length cuvettes*

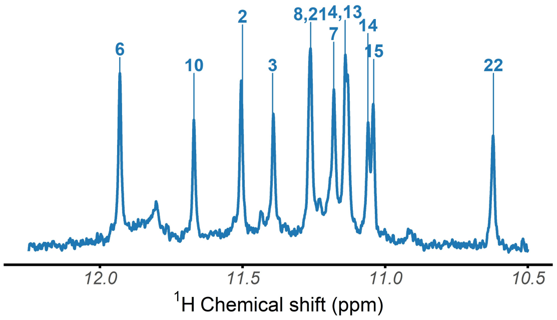

*Figure S117. ¹H-NMR spectrum of the 2O3M oligonucleotide, acquired at 25°C in 100 mM TMAA (pH 7.0) + 1 mM KCl*

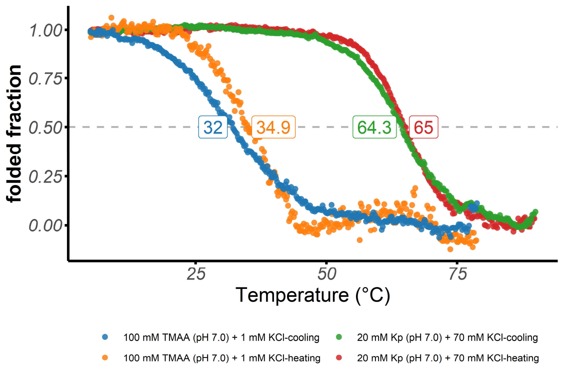

*Figure S118. Folded fraction of the 2O3M oligonucleotide as a function of temperature, determined by UV-melting (λ = 295 nm)*

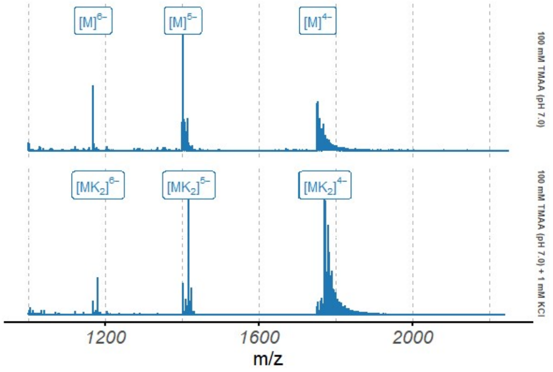

*Figure S119. Native ESI-MS spectra of the 2O3M oligonucleotide (10 µM)*

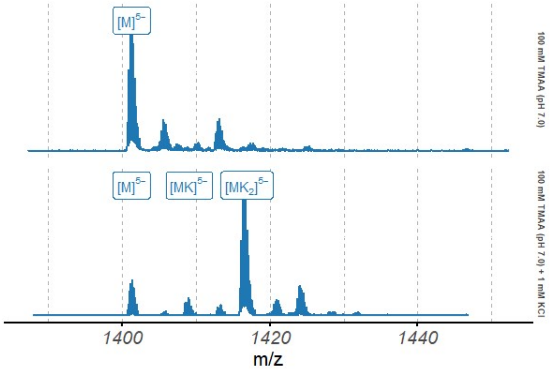

*Figure S120. Native ESI-MS spectra of the 2O3M oligonucleotide (10 µM), focused on the 5⁻ charge state*

#### 5NYS

*Table S20. Species information*

| Sequence (5' to 3') | $\epsilon_{260nm}$ $(M^{-1}cm^{-1})$ | DOI |
|---|---|---|
| TAGGGACGGGCGGGCAGGGT | 198600 | 10.1093/nar/gky250 |

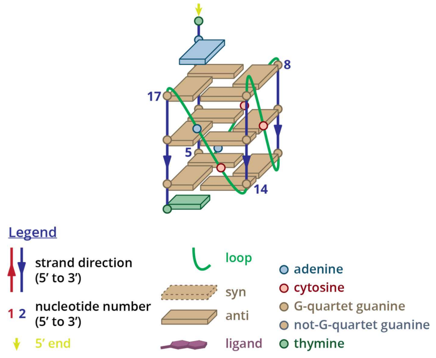

*Figure S121. Structure diagram of 5NYS*

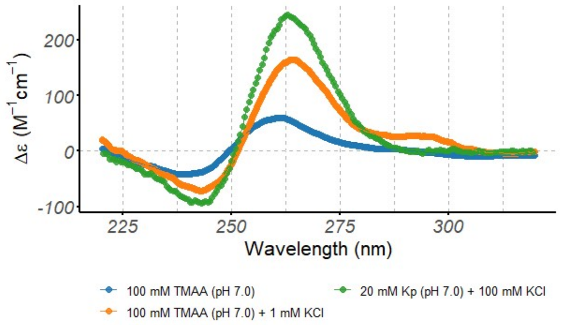

*Figure S122. Circular dichroism spectra of the 5NYS oligonucleotide (10 µM), acquired at 25°C in 0.4-cm path-length cuvettes*

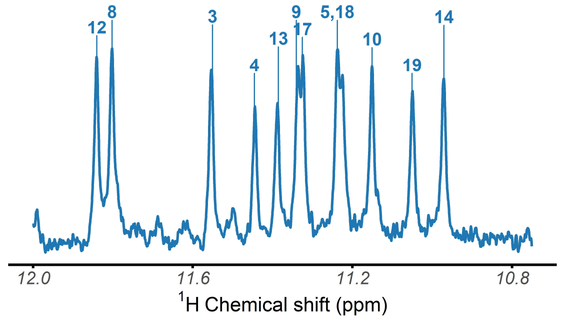

*Figure S123. ¹H-NMR spectrum of the 5NYS oligonucleotide, acquired at 25°C in 100 mM TMAA (pH 7.0) + 1 mM KCl*

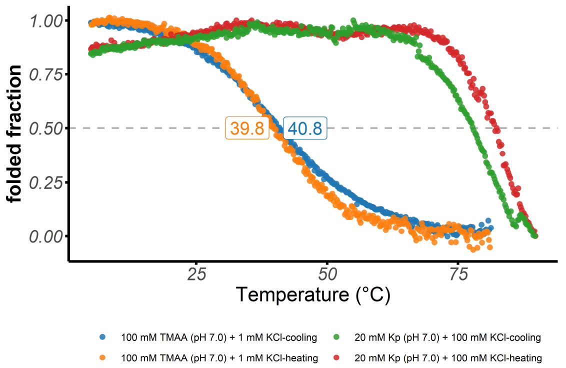

*Figure S124. Folded fraction of the 5NYS oligonucleotide as a function of temperature, determined by UV-melting (λ = 295 nm)*

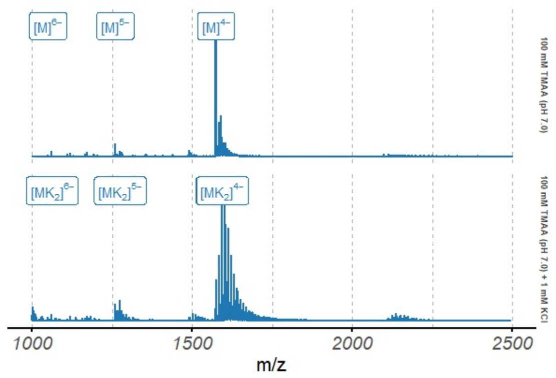

*Figure S125. Native ESI-MS spectra of the 5NYS oligonucleotide (10 µM)*

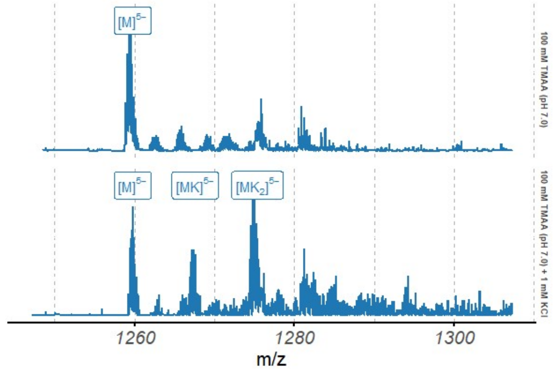

*Figure S126. Native ESI-MS spectra of the 5NYS oligonucleotide (10 µM), focused on the 5⁻ charge state*

### 2KYP

*Table S21. Species information*

| Sequence (5' to 3') | $\epsilon_{260nm}$ $(M^{-1}cm^{-1})$ | DOI |
|---|---|---|
| CGGGCGGGCGCTAGGGAGGGT | 202200 | 10.1093/nar/gkq558 |

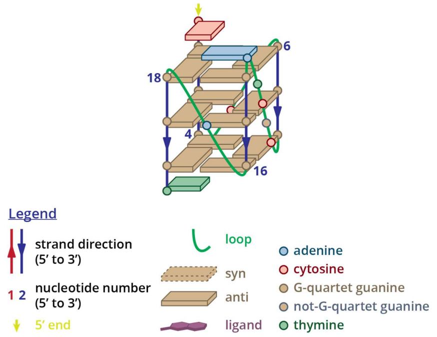

*Figure S127. Structure diagram of 2KYP*

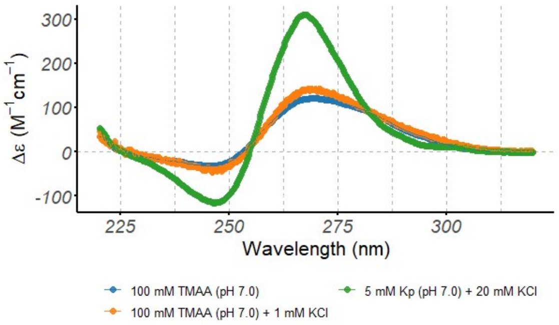

*Figure S128. Circular dichroism spectra of the 2KYP oligonucleotide (10 μM), acquired at 25°C in 0.4-cm path-length cuvettes*

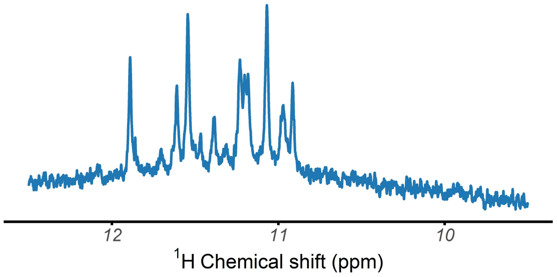

*Figure S129. ¹H-NMR spectrum of the 2KYP oligonucleotide, acquired at 25°C in 100 mM TMAA (pH 7.0) + 1 mM KCl*

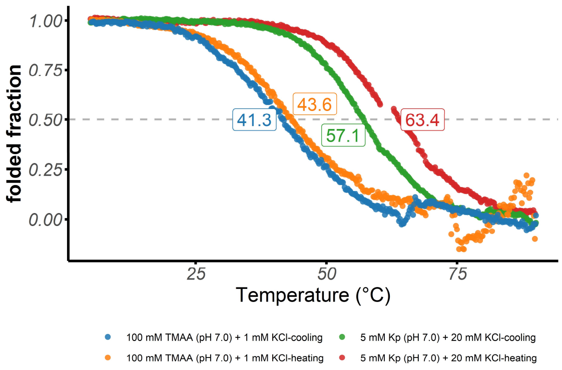

*Figure S130. Folded fraction of the 2KYP oligonucleotide as a function of temperature, determined by UV-melting (λ = 295 nm)*

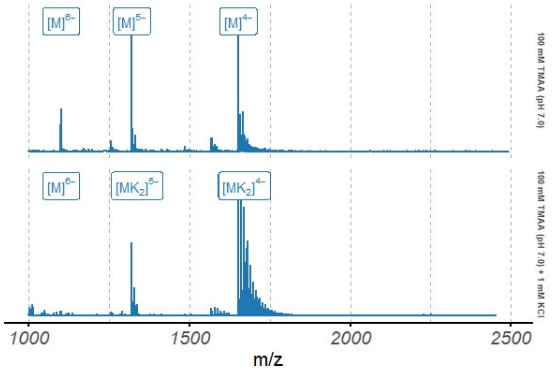

*Figure S131. Native ESI-MS spectra of the 2KYP oligonucleotide (10 μM)*

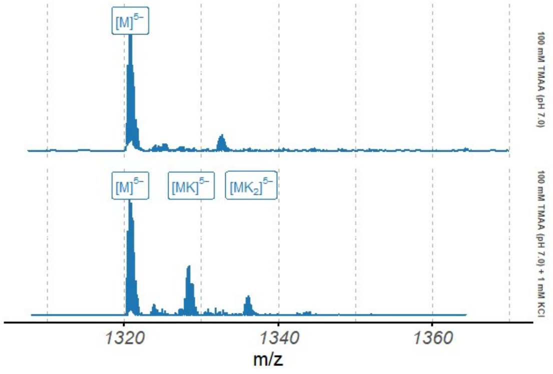

*Figure S132. Native ESI-MS spectra of the 2KYP oligonucleotide (10 μM), focused on the 5⁻ charge state*

# 2N4Y

*Table S22. Species information*

| Sequence (5' to 3') | $\epsilon_{260nm}$ $(M^{-1}cm^{-1})$ | DOI |
|---|---|---|
| CTGGGCGGGACTGGGGAGTGGT | 211200 | 10.1093/nar/gkw432 |

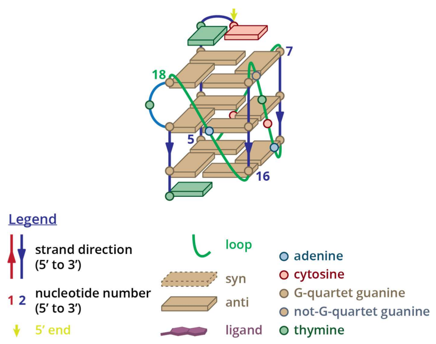

*Figure S133. Structure diagram of 2N4Y*

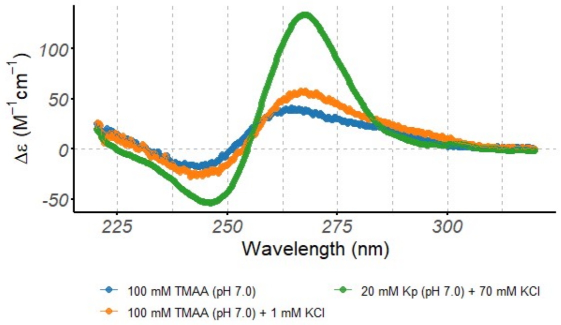

*Figure S134. Circular dichroism spectra of the 2N4Y oligonucleotide (10 µM), acquired at 25°C in 0.4-cm path-length cuvettes*

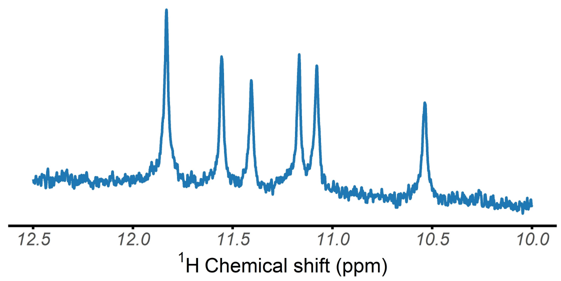

*Figure S135. $^{1}$H-NMR spectrum of the 2N4Y oligonucleotide, acquired at 25°C in 100 mM TMAA (pH 7.0) + 1 mM KCl*

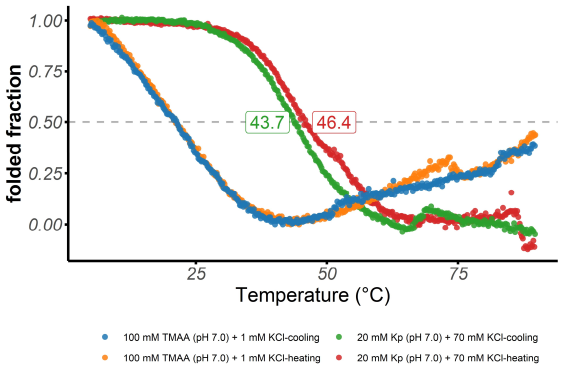

*Figure S136. Folded fraction of the 2N4Y oligonucleotide as a function of temperature, determined by UV-melting (λ = 295 nm)*

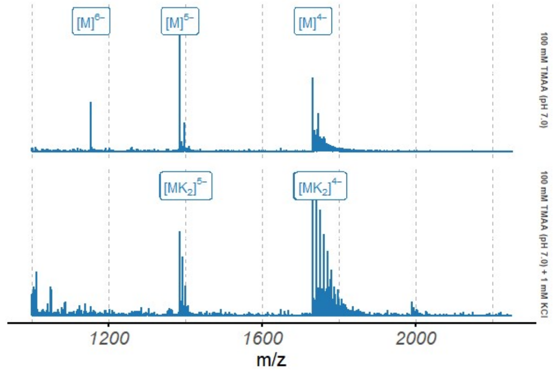

*Figure S137. Native ESI-MS spectra of the 2N4Y oligonucleotide (10 µM)*

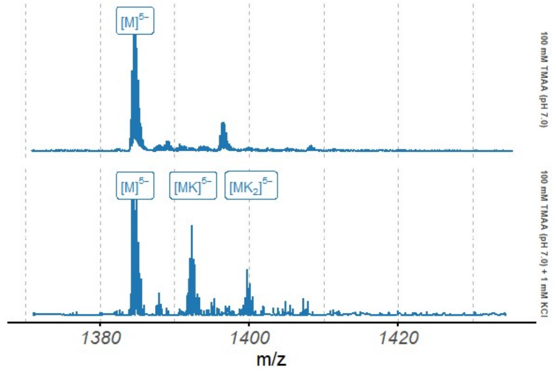

*Figure S138. Native ESI-MS spectra of the 2N4Y oligonucleotide (10 µM), focused on the 5⁻ charge state*

## 5I2V

*Table S23. Species information*

| Sequence (5' to 3') | $\epsilon_{260nm}$ $(M^{-1}cm^{-1})$ | DOI |
|---|---|---|
| AGGGCGGTGTGGGAATAGGGAA | 233100 | 10.1074/jbc.M117.781906 |

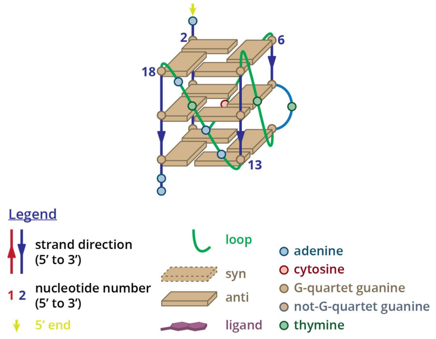

*Figure S139. Structure diagram of 5I2V*

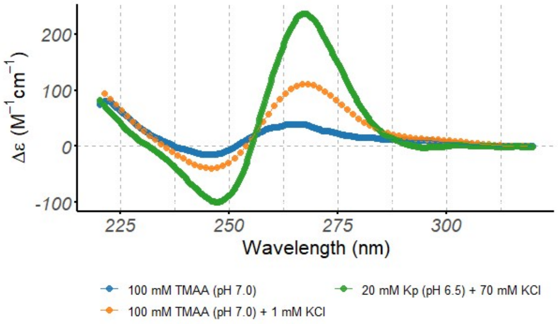

*Figure S140. Circular dichroism spectra of the 5I2V oligonucleotide (10 μM), acquired at 25°C in 0.4-cm path-length cuvettes*

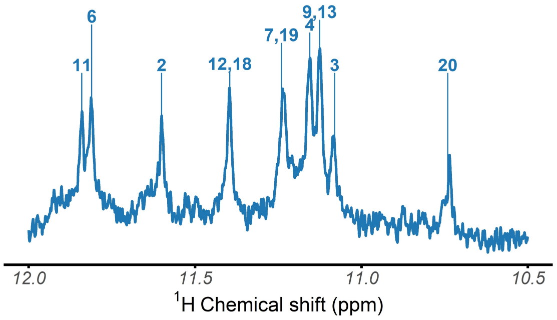

*Figure S141. $^{1}$H-NMR spectrum of the 5I2V oligonucleotide, acquired at 25°C in 100 mM TMAA (pH 7.0) + 1 mM KCl*

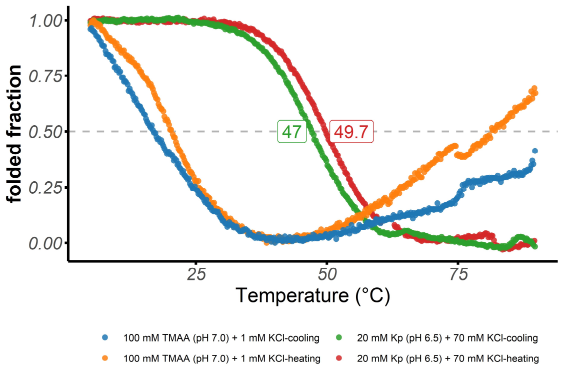

*Figure S142. Folded fraction of the 5I2V oligonucleotide as a function of temperature, determined by UV-melting (λ = 295 nm)*

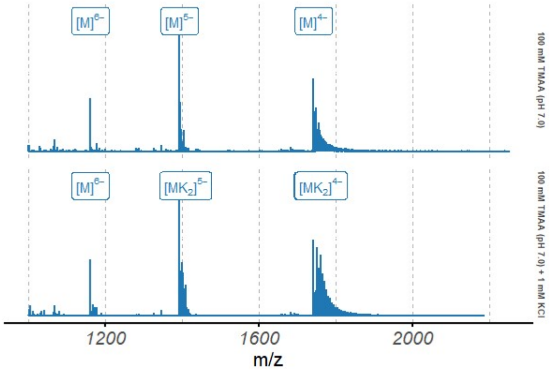

*Figure S143. Native ESI-MS spectra of the 5I2V oligonucleotide (10 μM)*

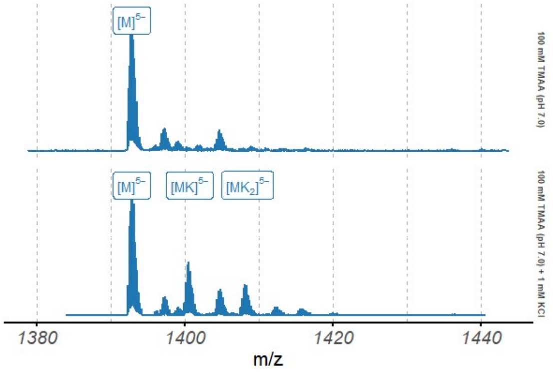

*Figure S144. Native ESI-MS spectra of the 5I2V oligonucleotide (10 μM), focused on the 5⁻ charge state*

### 2KPR

*Table S24. Species information*

| Sequence (5' to 3') | $\epsilon_{260nm}$ $(M^{-1}cm^{-1})$ | DOI |
|---|---|---|
| GGGTGGGGAAGGGGTGGGT | 193900 | 10.1016/j.str.2009.10.015 |

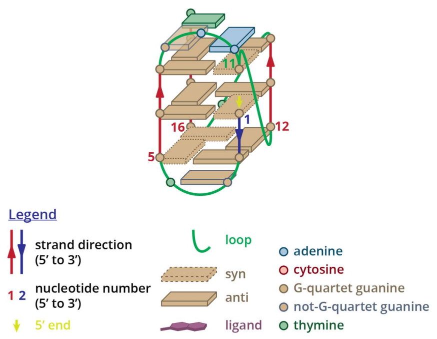

*Figure S145. Structure diagram of 2KPR*

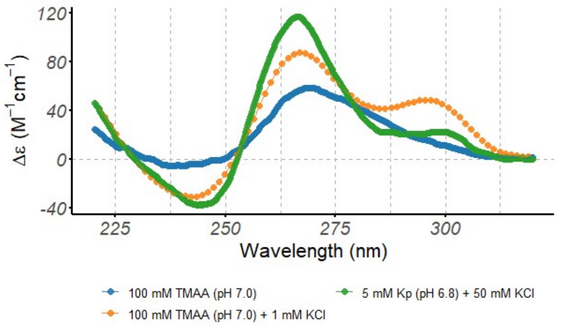

*Figure S146. Circular dichroism spectra of the 2KPR oligonucleotide (10 µM), acquired at 25°C in 0.4-cm path-length cuvettes*

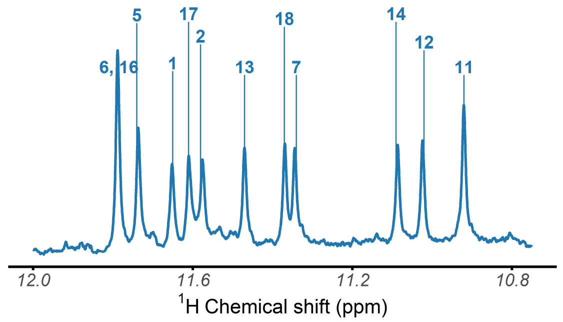

*Figure S147. $^1$H-NMR spectrum of the 2KPR oligonucleotide, acquired at 25°C in 100 mM TMAA (pH 7.0) + 1 mM KCl*

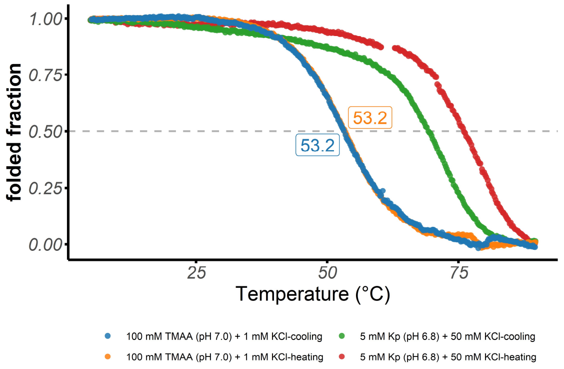

*Figure S148. Folded fraction of the 2KPR oligonucleotide as a function of temperature, determined by UV-melting (λ = 295 nm)*

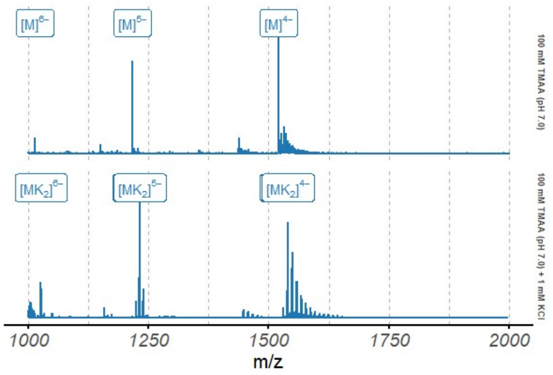

*Figure S149. Native ESI-MS spectra of the 2KPR oligonucleotide (10 µM)*

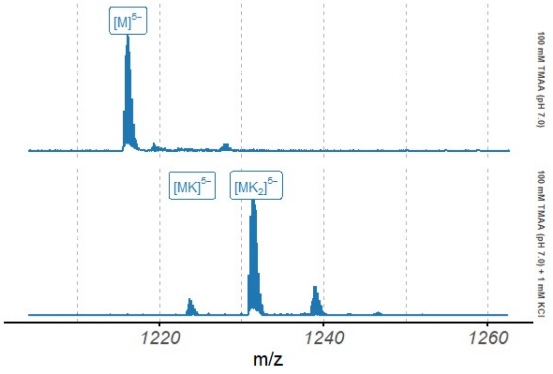

*Figure S150. Native ESI-MS spectra of the 2KPR oligonucleotide (10 µM), focused on the 5$^-$ charge state*

#### 2LOD

*Table S25. Species information*

| Sequence (5′ to 3′) | $\epsilon_{260nm}$ $(M^{-1}cm^{-1})$ | DOI |
|---|---|---|
| GGGATGGGACACAGGGGACGGG | 226400 | 10.1093/nar/gks329 |

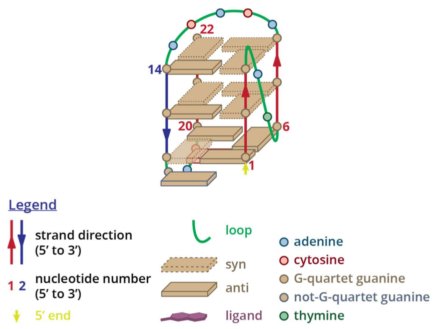

*Figure S151. Structure diagram of 2LOD*

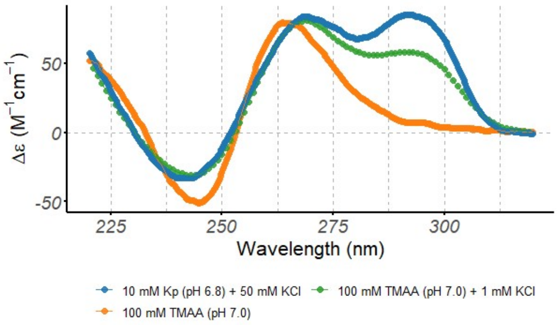

*Figure S152. Circular dichroism spectra of the 2LOD oligonucleotide (10 µM), acquired at 25°C in 0.4-cm path-length cuvettes*

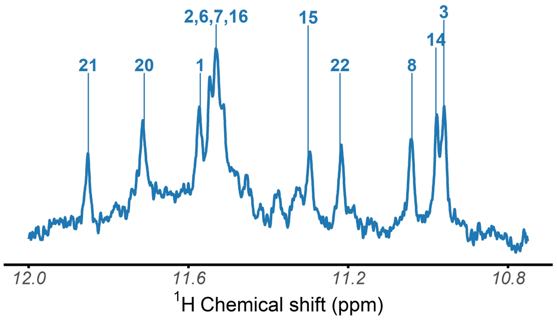

*Figure S153. $^{1}$H-NMR spectrum of the 2LOD oligonucleotide, acquired at 25°C in 100 mM TMAA (pH 7.0) + 1 mM KCl*

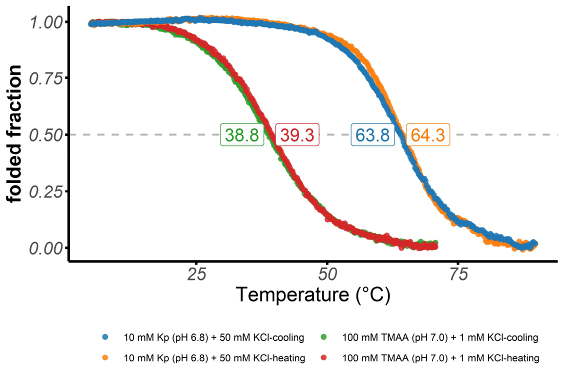

*Figure S154. Folded fraction of the 2LOD oligonucleotide as a function of temperature, determined by UV-melting (λ = 295 nm)*

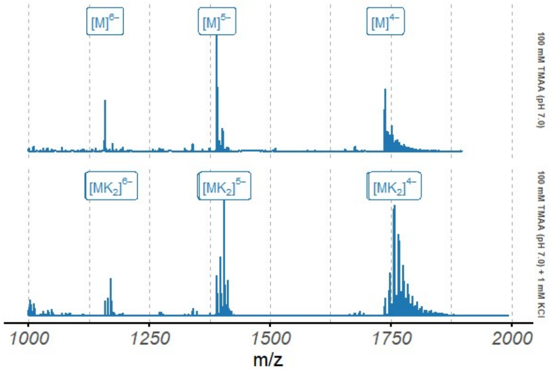

*Figure S155. Native ESI-MS spectra of the 2LOD oligonucleotide (10 µM)*

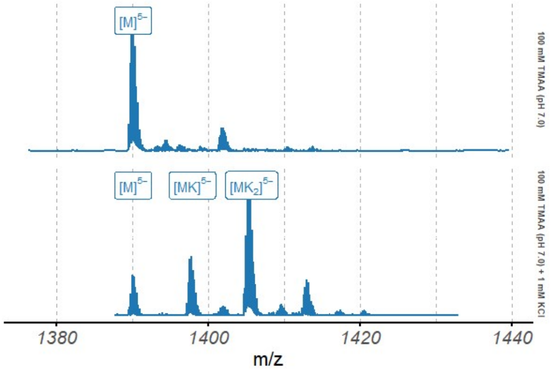

*Figure S156. Native ESI-MS spectra of the 2LOD oligonucleotide (10 µM), focused on the 5⁻ charge state*

### HIV-PRO1

*Table S26. Species information*

| Sequence (5’ to 3’) | $\epsilon_{260nm}$ $(M^{-1}cm^{-1})$ | DOI |
|---|---|---|
| TGGCCTGGGCGGGACTGGG | 163900 | 10.1021/ja501500c |

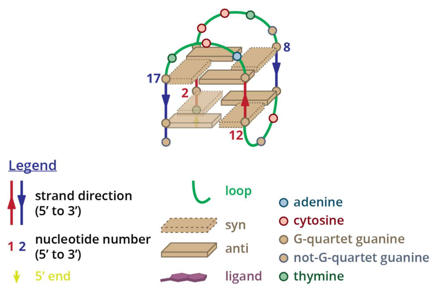

*Figure S157. Structure diagram of HIV-PRO1*

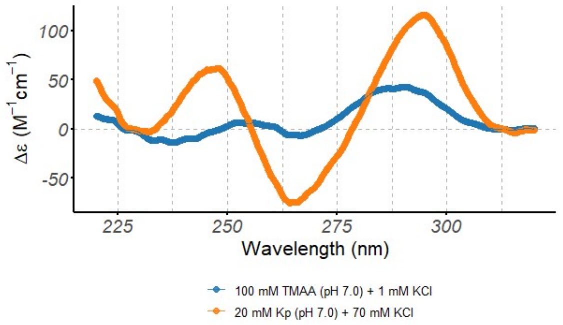

*Figure S158. Circular dichroism spectra of the HIV-PRO1 oligonucleotide (10 μM), acquired at 25°C in 0.4-cm path-length cuvettes*

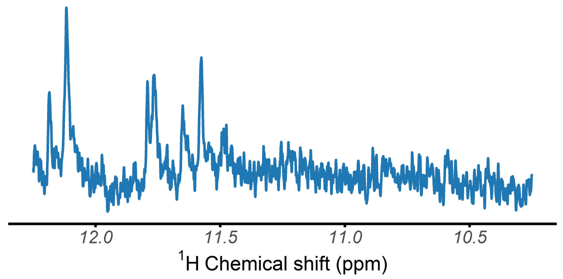

*Figure S159. $^{1}$H-NMR spectrum of the HIV-PRO1 oligonucleotide, acquired at 25°C in 100 mM TMAA (pH 7.0) + 1 mM KCl*

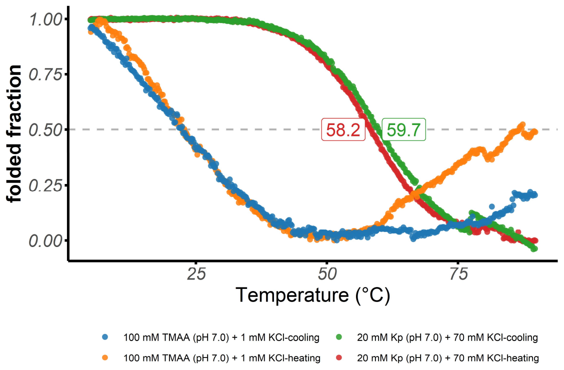

*Figure S160. Folded fraction of the HIV-PRO1 oligonucleotide as a function of temperature, determined by UV-melting (λ = 295 nm)*

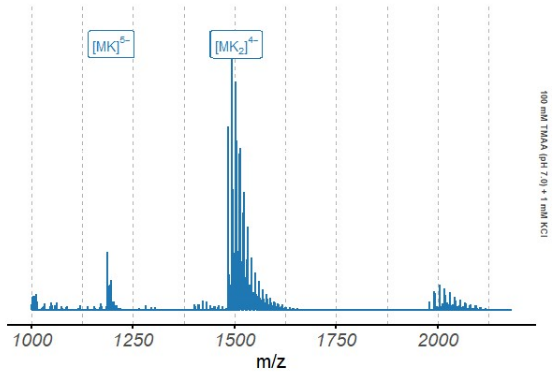

*Figure S161. Native ESI-MS spectra of the HIV-PRO1 oligonucleotide (10 μM)*

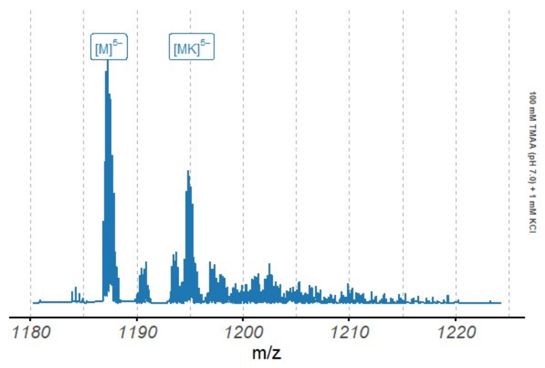

*Figure S162. Native ESI-MS spectra of the HIV-PRO1 oligonucleotide (10 μM), focused on the 5⁻ charge state*

#### 6GZN

*Table S27. Species information*

| Sequence (5' to 3') | $\epsilon_{260nm}$ $(M^{-1}cm^{-1})$ | DOI |
|---|---|---|
| GGGTAGGGAGCGGGAGAGGG | 211700 | 10.1002/anie.201809328 |

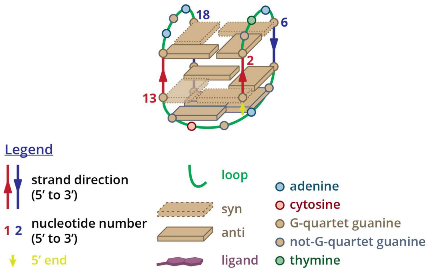

*Figure S163. Structure diagram of 6GZN*

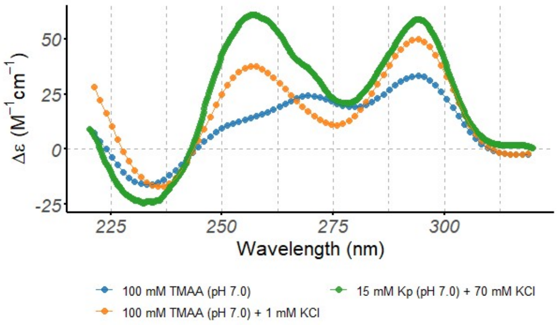

*Figure S164. Circular dichroism spectra of the 6GZN oligonucleotide (10 µM), acquired at 25°C in 0.4-cm path-length cuvettes*

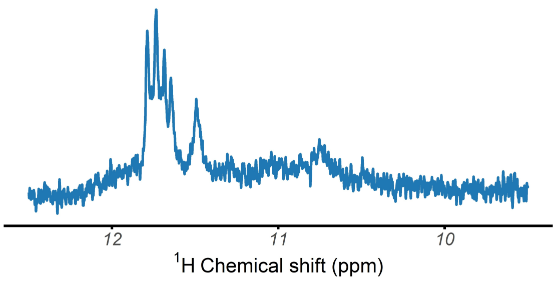

*Figure S165. $^{1}$H-NMR spectrum of the 6GZN oligonucleotide, acquired at 25°C in 100 mM TMAA (pH 7.0) + 1 mM KCl*

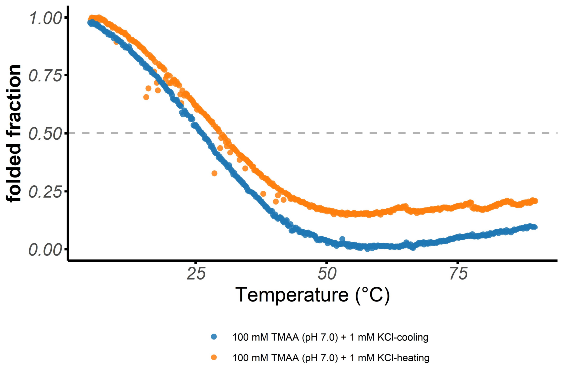

*Figure S166. Folded fraction of the 6GZN oligonucleotide as a function of temperature, determined by UV-melting (λ = 295 nm)*

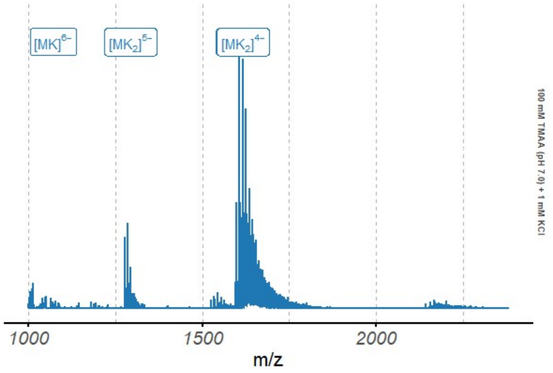

*Figure S167. Native ESI-MS spectra of the 6GZN oligonucleotide (10 µM)*

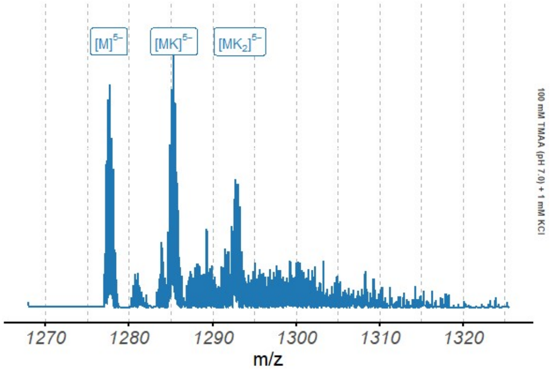

*Figure S168. Native ESI-MS spectra of the 6GZN oligonucleotide (10 µM), focused on the 5⁻ charge state*

### g4dbr

Eric Largy

2020-11-02

**References** **71**

*Copyright 2020 Eric Largy. Licensed under the GPL-3 license.*

### 1 General overview

#### 1.1 Intended and less-intended uses

*g4dbr* is an R package containing the Shiny app *g4db* that is dedicated to the creation, visualization, and reporting of curated circular dichroism (CD), $^{1}$H-NMR, UV-melting and native mass spectrometry (MS) data from oligonucleotides. Although specifically developed for G-quadruplex forming sequences deposited in the PDB, *g4dbr* can be used with any nucleic acid sequence.

Users can either employ the app to visualize a database generated by *g4db*, visualize data pasted into a templated Excel file (provided in the package), and create/edit/complete a *g4db* database from data supplied in said template.

The long-term goal is to provide tools for the robust deposition of raw experimental data, and processed data derived from them, while allowing for easy and versatile visualisation and reporting.

Raw data pasted in the supplied Excel template can be deposited, and visualized in several ways, which are open to other scientists without the need for proprietary software. The approach is two-fold:

##### 1.1.1 Templated .xlsx file deposition as is

Once pasted into the input template, the data can be deposited as is. It can then be explored natively in Excel or any open-source equivalent. The data is formatted in a non-ambiguous layout, provided it is properly labeled in the header cells.

The template is also amenable to software allowing header cell import/management, such as Origin, in which import scripts can be used.

Of course, the template can be natively imported in the *g4db* app. The advantages over Excel/Origin for this particular application are numerous in terms of both ease and speed of use (e.g. data filtering, automated figure plotting), and functionalities (e.g. peak labeling, normalization/calculation, selective data export). See the Main features section for more details.

Any data treatment and filtering performed within *g4db* is not saved into the input .xlsx file. To save this into a new or existing database file, the second approach must be used:

##### 1.1.2 Rdata file

*g4db* allows exporting selected datasets into an RData (.Rda) file where the data is consolidated and all calculation has already been performed. This leads to faster figure display, smaller file size, and is amenable to host very large datasets (where Excel is limited in row numbers, which is particularly problematic for mass spectrometry data).

The downside of this approach is that it cannot be handled outside of R. Note, however, that *g4db* is not required to open and use the data, it can be natively loaded in base R, which is free and open source. To do so, use the `load` function, for instance below for a demo database provided in the package:

```
load(system.file("extdata/demo_database.Rda", package = 'g4dbr'))
```

#### 1.2 Extended scope

*g4dbr* includes a number of functionalities that will be described here within the context of their intended use, but that can be utilized outside of this scope, *i.e.*

- automated or semi-automated data filtering, treatment and labeling,

- computation of molar extinction coefficient ($\lambda$ = 260 nm) of oligonucleotides (*epsilon.calculator*),
- UV-melting data treatment (*meltR*),
- MS data size reduction (*mass.diet*)
- Database selective data deletion (*database.eraser*)

#### 1.3 Main features

Below is a list of the main features of *g4dbr*.

- Visualization of CD, UV-melting, $^1$H-NMR and native MS data gathered in a database (.Rda format)
  - Collapsible and tabulated interface
  - Quick and user-friendly data filtering in tables and figures (*e.g.* oligonucleotide, buffer, cation, x-axis range,…)
    - Automated buffer list collection
    - Automated tune and replicate collection
  - Control over the database content, display, and reporting (see below)
- Robust database creation and edition
  - Data imported from a templated Excel file
  - Selective data importing (by *e.g.* technique/oligo/buffer/data range)
  - Duplicate detection/suppression
  - Automated deposition date and DOI link generation for traceability purpose
  - Replication management for MS and UV-melting data
  - Different tune management for MS data
- Automated data treatment
  - Conversion of CD to molar ellipticities
  - MS data normalization
  - $^1$H-NMR and MS peak labeling
  - UV-melting data normalization and conversion to folded fraction
  - UV-melting thermodynamic quantities determination
  - UV-melting Tm labeling
- Custom figures
  - Control over colors, size, and transparency of figures
  - Color palettes adapted to qualitative, sequential, and diverging data
  - Switch between overlaid and paneled figures for quick comparisons
  - Control over variables mapped in paneled figures
  - Automated colour mapping to non-paneled variables
  - Automated figure dimension change to accommodate multiple rows
- Automated report generation
  - Full or Supporting information dedicated reports
  - pdf, HTML and docx formats
  - All data, figure captions, figure sizing, file name, etc. generated dynamically without user input
- Open
  - Coded in R
  - Easy-to-export data tables (practical for standalone data treatment)
  - Import template easy to read in other software
  - Full code and experimental data hosted openly on GitHub

#### 1.4 Workflow

For raw data import, the data must be pasted into a templated Excel file, then read in the *importR* module of *g4db*. In this module, the data can be filtered, processed, and selected for writing into a database file (.Rda). The .Rda file can be opened in the *database* module for visualization and reporting purposes. It can also simply be loaded in base R for further processing or reporting steps that may not be possible in *g4db*.

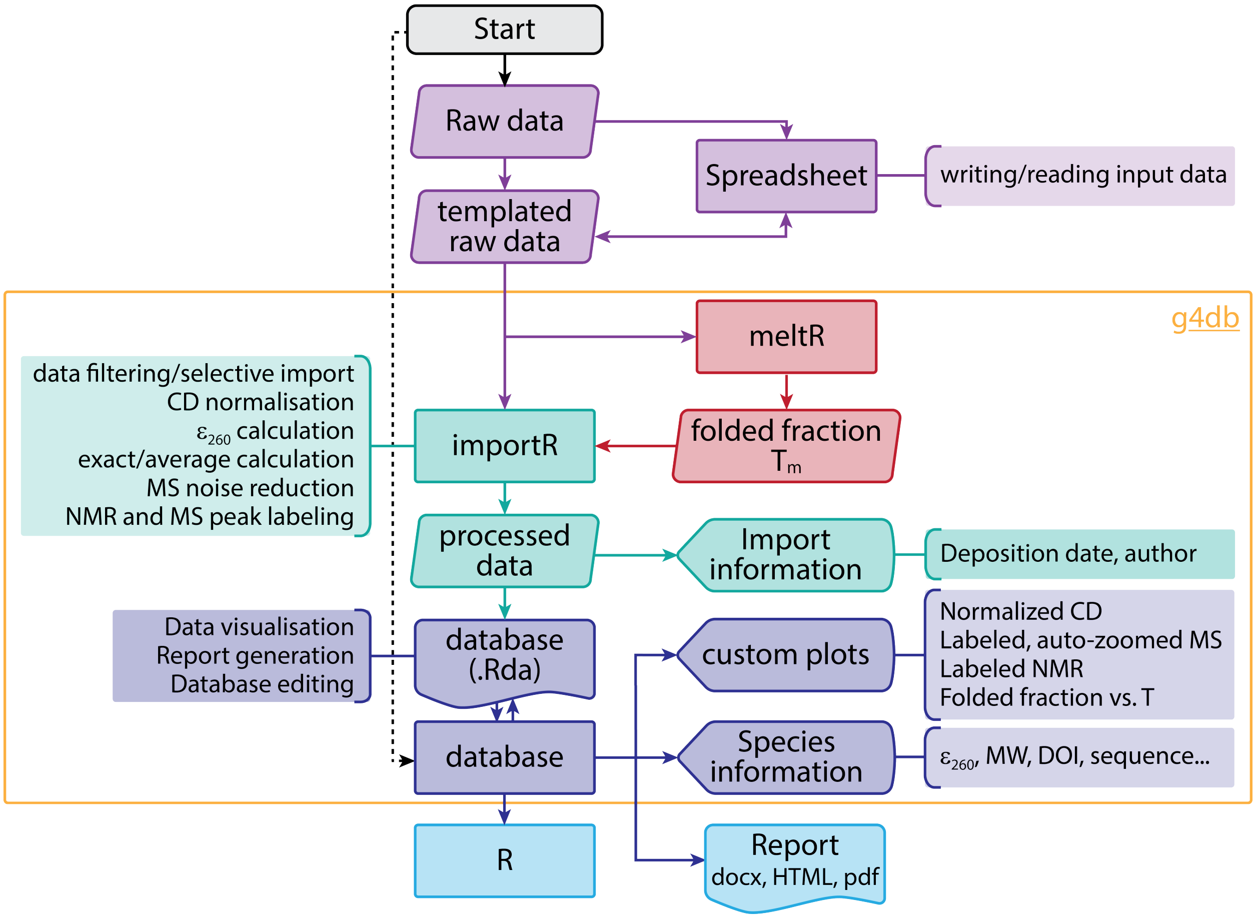

Figure S169: Application workflow

### 2 Installation and setup

#### 2.1 Installation

Install from Github using:

```
install.packages("devtools")
devtools::install_github('EricLarG4/g4dbr')
```

Alternatively, download the .zip archive from GitHub then run:

```
install.packages("devtools")
devtools::install_local("XXX/g4dbr-master.zip")
```

Where `XXX` is the file path to the zip archive.

##### 2.2 Setup

Load the package with:

```
library(g4dbr)
```

## 3 *g4db*

##### 3.1 Running the app

Only one function must be called to use all functionalities from *g4dbr*:

```
g4db()
```

This function opens a Shiny app in either the currently used IDE (e.g. RStudio), or a web browser.

Other functions used in *g4db* are packaged in *g4dbr*, and can be used as standalone tools. Refer to the Other functions and reference files section.

##### 3.2 Interface overview

The interface is divided in 3 tabs that can be selected at the top of the screen, and are used to accomplished specific tasks:

- *database*, to visualize, report, and remove data from a database file.
- *importR*, to visualize and process raw data, and export all or part of it to a database file,
- *meltR*, to visualize and treat UV-melting data, and export all or part of it to the a database (via *importR*).

The tabs make use of various sidebars, mainly to perform data importing, filtering, processing, exporting and reporting.

###### 3.2.1 Figures and tables

In the main area of the interface are the figures and tables, within collapsible and closable boxes, letting the user select what data to display.

All tables are sortable and filterable to assist in exploring rich data sets, and find specific data points rapidly. The data is presented in *long format*, which makes it easier to filter through, and to map variables into figures, because each variable is contained in its own column. Columns can be selectively hidden, and some of the less relevant ones are hidden by default.

Data presented in figures and tables reflects the values given to the different filters. On the contrary, filtering the tables does *not* alter the figures, it is only a mean of accessing and/or exporting a subset of the data.

All tables can be exported as .csv, .xlsx, or in the clipboard. All columns will be exported, regardless of their visibility in the app.

##### 3.2.2 Sidebars and panels

**3.2.2.1 Left sidebars and panels** Each tab has a sidebar on the left-hand side, which contains a number of tools for data importing, exporting, filtering, and formatting. This *left sidebar* is collapsible to release some space for figures and tables on smaller screens. Each tab has a specific and independent *left sidebar*, and the values from those *left sidebar* modifies the data for *all* the content of the tab (and almost always only this tab). Drop-down menus contain *select all/deselect all* buttons for quick data selection.

Given the amount of menus necessary for the *meltR* tab, a large portion is hosted in two collapsible and movable “hovering” panels.

The sidebar from the *database* and *importR* tabs, and a panel of *meltR* also contain a color palette selection menu, and submenu for certain palettes having variations (Figure S170). The available palettes include:

- The well known Brewer palettes that include qualitative, diverging, and sequential palettes,
- Some discrete palettes from D3.js, a JavaScript library for producing interactive data visualizations (imported from the `ggsci` package),
- Several palettes inspired by the colors used by scientific journals/publishers (NPG, AAAS, NEJM, Lancet, JAMA, JCO, etc.; imported from the `ggsci` package).

The selected colour palette is applied to all the figures of the tab, but does not affect other tabs.

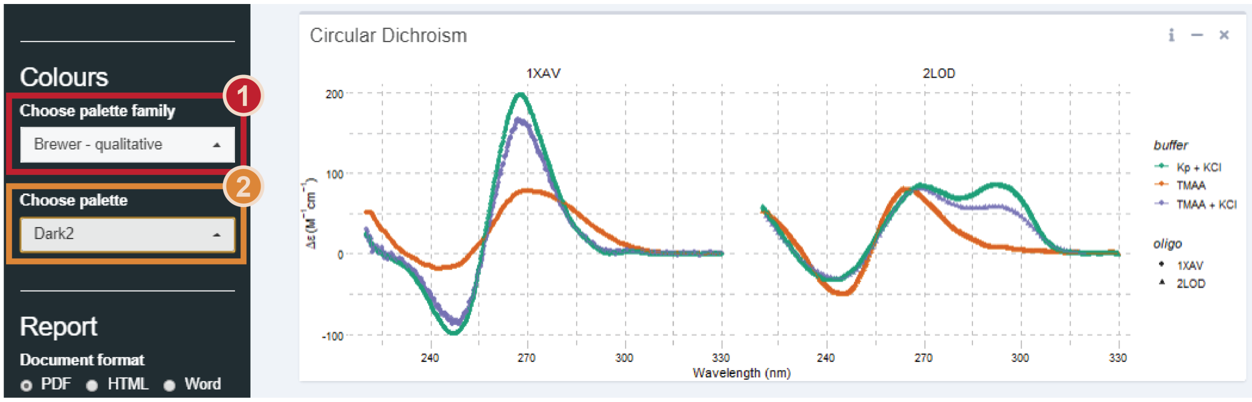

Figure S170: Colour palette selection. Some palette families (1) containing several palette variations (2)

**3.2.2.2 Right sidebars** Figure boxes feature a *right sidebar*. They contain filtering and data formatting filters that are applied *only* on the corresponding figure (contrary to the *left sidebars* that affect entire tabs). These sidebars are collapsible as well, and hidden by default.

#### 3.3 Consulting a database: the *database* tab

The database tab is dedicated to visualizing, exporting, and reporting on the data of a curated database file.

##### 3.3.1 Database input

The data from a given database must be gathered in a single .Rda file generated in the *importR* tab. It contains five dataframes: one dedicated to the general oligonucleotide information (`db.info`), and the four other ones to each analytical technique (`db.CD`, `db.NMR`, `db.MS`, `db.UV`).

*g4db* extracts automatically all the data, but it can also be loaded in the global environment (*i.e.* without using *g4db*) using `load()`. For instance, to load the demo database, run:

```r
load(system.file("extdata/demo_database.Rda", package = 'g4dbr'))
```

The global environment should now contain five dataframes that can be opened and worked with. When using `g4db()`, the data will be loaded in the package environment and will therefore not appear in the global environment.

##### 3.3.2 Database use

**3.3.2.1 Data loading** Upon opening the database file, the interface should be devoid of data. The first step is to import a database file:

1. Click on *Browse* in the *Load* section of the *left sidebar* (Figure S171),
2. Select a .Rda file that has been prepared in *importR*

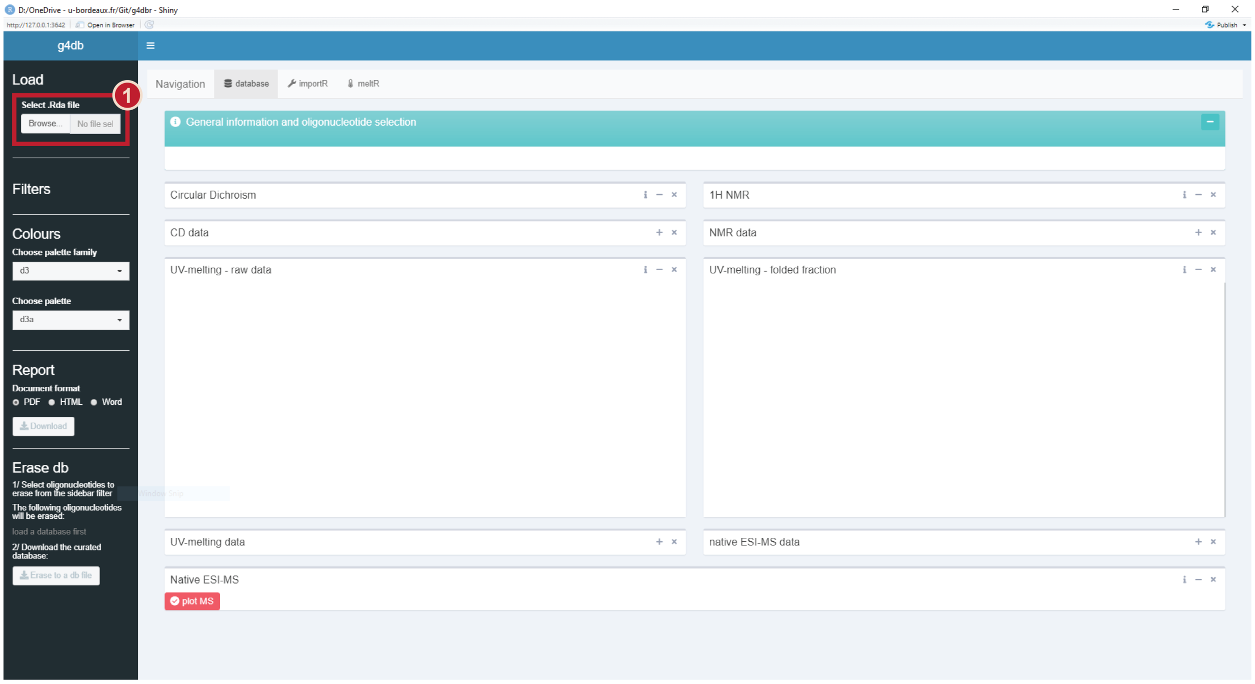

Figure S171: Empty database view

The *General information and oligonucleotide selection* table should now be populated by a list of the oligonucleotides for which the database file contains at least information data (Figure S172-1).

The content of this table is controlled by a drop-down menu in the *left sidebar*, and by the oligonucleotide column filter (in that order) (Figure S172-2). By default, all oligonucleotides are shown, but none are selected for analytical result display (to avoid wait times when the table content is changed).

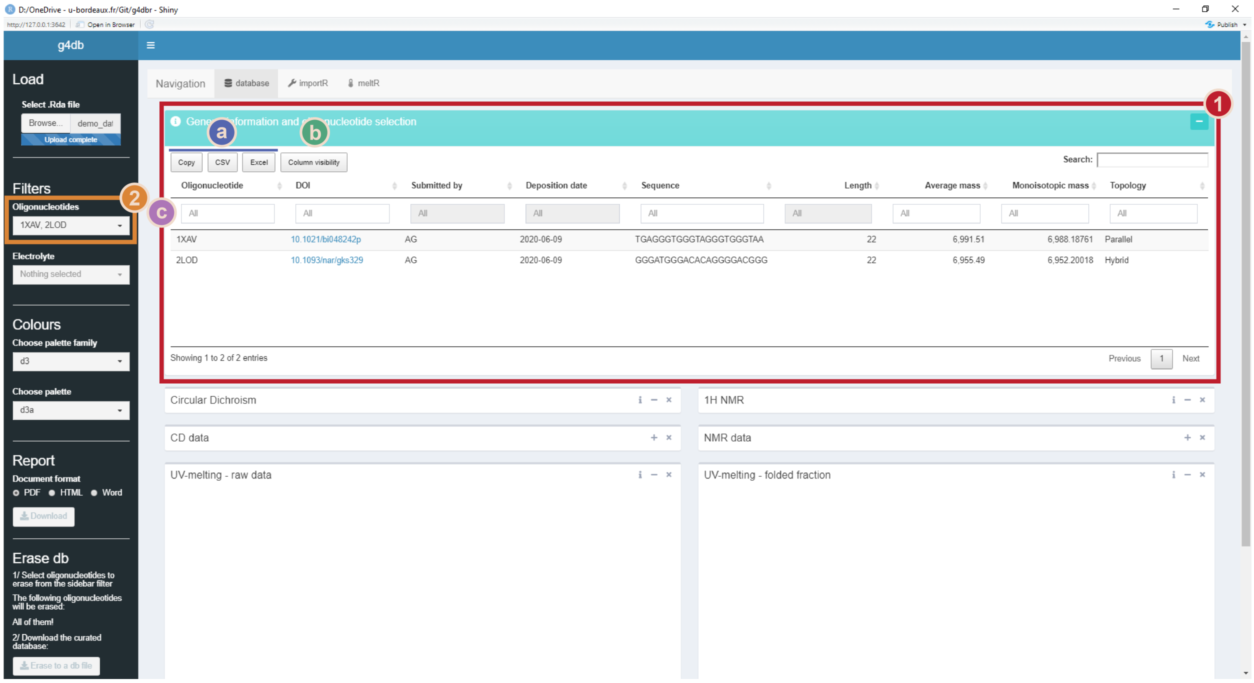

Figure S172: Demo database loaded in the database tab: the general oligonucleotide information should be displayed (1). The visible oligonucleotides can be filtered in the table or from the dropdown menu in the left sidebar (2). The table (1), and other tables in g4db, can be exported (a), their column visibility changed (b), and their content sorted, filtered or searched through (c)

**3.3.2.2 Data display** To start visualizing data, the *oligonucleotide(s)* of interest must be selected from the *General information* table, by clicking on one or several rows (Figure S173-1). Clicking again on a row deselects it.

The *CD*, *NMR* and *UV-melting* data should now be displayed (Figures S173-2 and S174-1). By default, the data acquired for all *buffer* conditions (i.e. all *cation* + *electrolyte*) are shown, but it can be restricted to only certain *buffers*, *electrolytes* or *cations*, using the menus from the *left sidebar* (Figure S173-3). Individual *cation* and *electrolyte* selections supersede the *buffer* selection. For instance, if the buffers “TMAA + KCl” and “Kp + KCl” are selected, but the “Kp” electrolyte is excluded, then only “TMAA + KCl” will effectively be selected.

Note that the *buffers*, *electrolytes* and *cations* are not a static list, but are automatically collected from the *CD* and *UV-melting* data. It is therefore important to keep their naming consistent across the entire database.

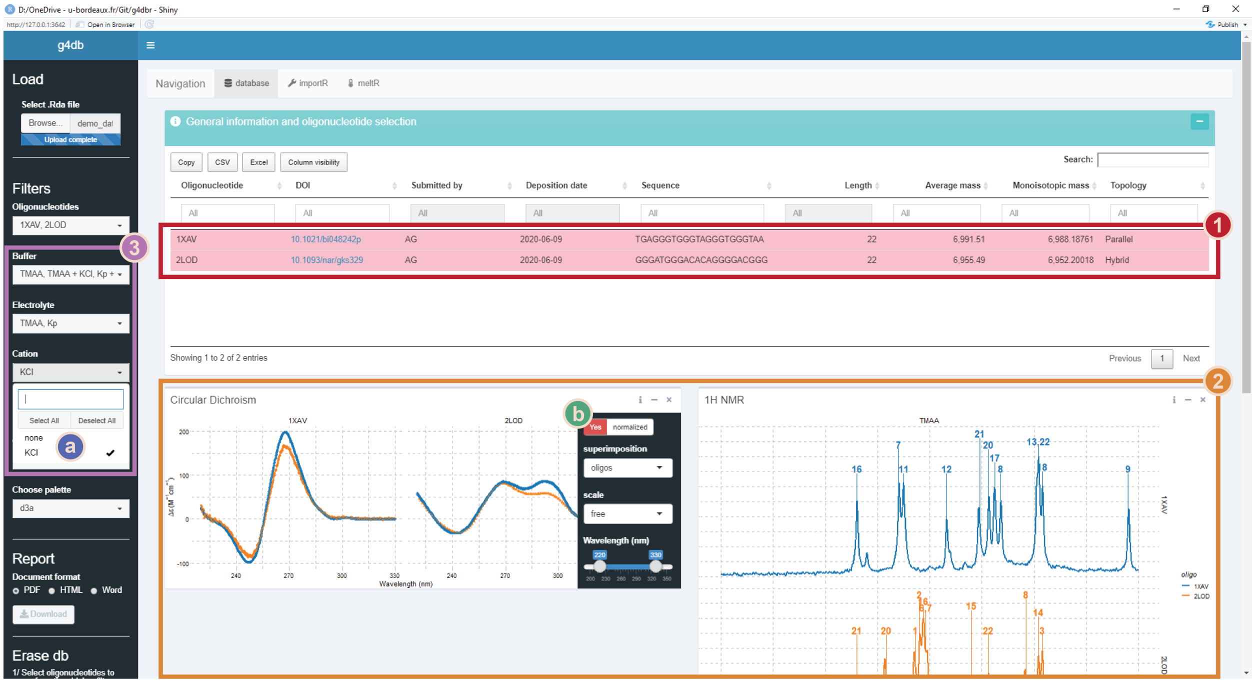

Figure S173: Database data display: both oligonucleotides have been selected (1). Their data is displayed (2) but was buffer-filtered (3): only KCl-containing solutions are selected (a). Using the right sidebar, the CD data was panelled by oligonucleotide (b)

The UV-melting data is displayed in two separate figures (S174-1): on the left is shown the raw data with the fit line, and on the right is processed data. Depending on whether the data was processed by non-linear fitting or not, the processed data will either be the *folded fraction* or the absorbance normalized in [0,1]. This allows to plot the data of highly stable or unstable species on the same figure as those for which the *Tm* could be determined.

To also display MS data (S174-2), the *Plot MS* button must be used S174-3. This avoid long refresh times when selecting oligonucleotides. Any change in the *oligonucleotide*, *buffer*, *tunes*, *replicates*, and *m/z* selections will only be effective if the figure is replotted.

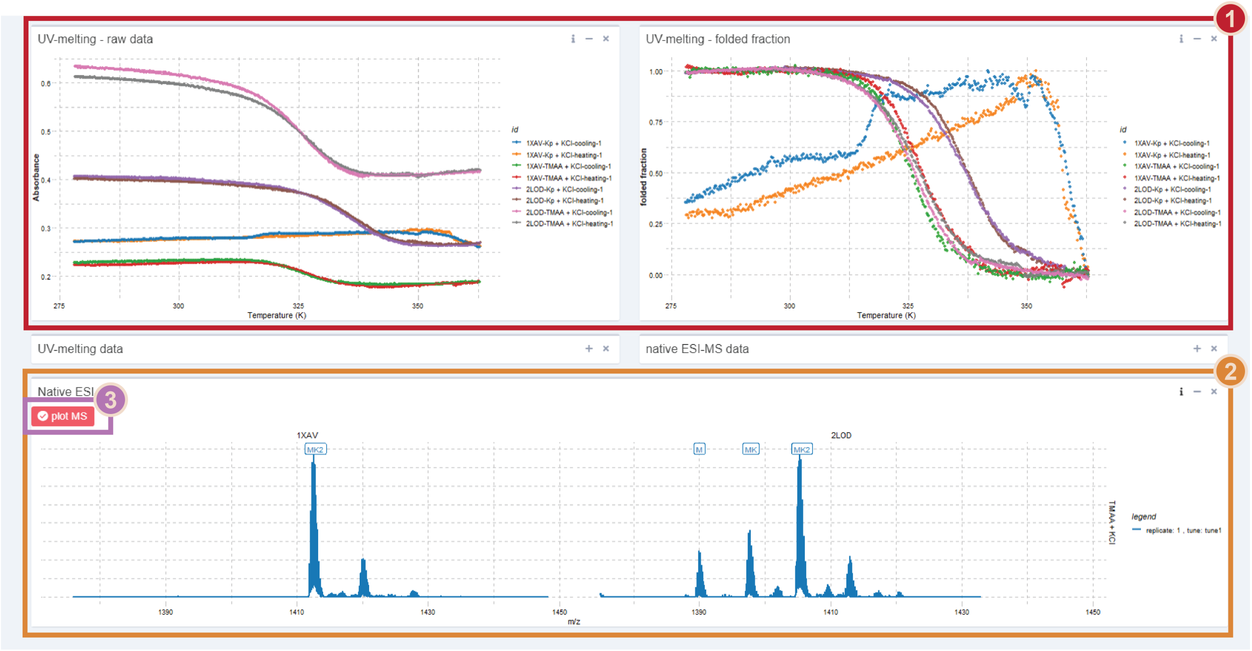

Figure S174: Database data display: UV-melting (1) and native MS (2) display. To display the MS data, the Plot MS button must be used (3)

For all these analytical methods, all data points are gathered in tables, collapsed by default. These data points can be sorted, filtered, and exported in .xlsx or .csv files, or copied in the clipboard (Figure S172). Again, filtering data in the tables does **not** affects the figures, only the *left* and *right sidebars* do.

##### 3.3.3 Data content and customization

**3.3.3.1 General information** This table gathers all the general information on the deposited oligonucleotides. By default, the following variables are displayed:

- `Oligonucleotide` name, preferably a PDB code where available
- `DOI`, with a hyperlink that is automatically generated upon importing with *importR*
- `Submitted by`, the initials or full name of the data submission author
- `Deposition date`, which is generated automatically by *g4db*
- `Sequence`, the 5' to 3' oligonucleotide sequence
- `Length`, the number of nucleotides, generated automatically by *g4db*
- `Average mass` and `Monoisotopic mass` of the oligonucleotide, generated automatically by *g4db*, and used for the native MS peak labelling
- `Extinction coefficient (260 nm)`, the molar extinction coefficient of the oligonucleotide (in $M^{-1}cm^{-1}$), calculated automatically by *g4db* (via the *epsilon.calculator*)
- `Topology`, a short user-supplied description of the oligonucleotide structure (e.g. *parallel quadruplex*)

The fields hidden by default (nucleotide and atom numbers) are not of direct interest to the general user, but can be displayed using the *column visibility* button.

Importantly, this table is used to select the oligonucleotide for which the analytical data should be displayed, as shown in Figure S173. It is possible to quickly filter through entries by e.g. topology or length, to select all oligonucleotides falling in a given category.

###### 3.3.3.2 Circular dichroism

The data is shown as points and lines, colored by *buffer*. The *oligonucleotides* are differentiated by point shape.

The *right sidebar* contain the following settings:

- *normalized* switch: choose to display molar ellipticities (as automatically calculated in *importR*; default) or raw data (i.e. in mdeg).
- *superimposition* dropdown menu: choose to display all data superimposed (default), grouped in panels by *oligonucleotide* or *buffer*, or not superimposed at all.
  - The figure size will automatically adjust with the number of panels
- *scale* dropdown menu: select whether all panels must have the same y-axis scale (*not free*) or can be rescaled to better fit their content (*free*)
- *Wavelength* slider: select the wavelength range to display (default: 220-330)
- *point size* and *line size* sliders: adjust the size of points and lines
- *transparency* slider: adjust the transparency of both points and lines

The data is gathered in the *CD data* table below, which can be sorted, filtered, and exported. The fields displayed by default are `Oligonucleotide`, `Buffer`, `Wavelength (nm)`, `CD (mdeg)`, and `Delta epsilon (M-1cm-1)`. The other fields hidden by default can be displayed using the *column visibility* button.

###### 3.3.3.3 $^1$H NMR

The data is shown as a line, colored by *oligonucleotide*, and is normalized so that all spectra will share the same y-axis range. By default, each spectrum is shown in its own panel. Peak numbers are shown above their peaks and linked by a segment.

The *right sidebar* contains some settings identical to the CD one (*superimposition*, *scale*, *line size*). In addition, it contains a *chemical shift (ppm)* slider to select the chemical shift range to display (default: 9.5-12.5 ppm).

The data is gathered in the *NMR data* table below, which can be sorted, filtered, and exported. The fields displayed by default are `Oligonucleotide`, `Buffer`, `Chemical shift (ppm)`, and `Intensity`. The other fields hidden by default can be displayed using the *column visibility* button.

###### 3.3.3.4 UV-melting

UV-melting data is plotted with points, and in the case of the raw data with an additional fit line.

The *right sidebar* contains some settings identical to some described above (*point size*, *line size*, *line transparency*). In addition, it contains a *Temperature (K)* slider to select the temperature range to display (default: 278-368 K).

The data is gathered in the *UV-melting data* table below, which can be sorted, filtered, and exported. The fields displayed by default are `Oligonucleotide`, `Buffer`, `ramp`, `T (K)`, `Folded fraction`, and `Absorbance`. The other fields hidden by default can be displayed using the *column visibility* button.

###### 3.3.3.5 Native mass spectrometry

There are two distinct plots to visualize MS data, i.e. one full scale and one charge-state focused, to better see the potassium adduct distribution.

In both cases, the data is shown as line, with labels to name the visible species (Figure S175). By default, spectra are paneled by *oligonucleotide* (columns) and *buffer* (rows), which should typically lead to a single spectrum per panel. Peak labels appear above their corresponding peak. The focused plot displays the 5- charge state by default, but this can be changed by the user.

Besided a *line size* slider, the *right sidebar* of the full-scale plot contains:

- *m/z* slider: select the *m/z* range to display (default: 800-2500 *m/z*).
- *Tunes* dropdown menu: select the *tunes* to display.

- *tunes* are collected automatically from the data

- *Replicates* dropdown menu: select the *replicates* to display
  - *replicates* are collected automatically from the data
- *Layout* dropdown menu: select a panel layout among all combinations of *oligonucleotide*, *tune*, *buffer*, and *replicate*
  - Six unique combinations can be selected, and the six other ones are accessed using the *transpose grid* switch
  - If more than one spectrum appears on a panel, the two variables that are not mapped by the layout are combined to be mapped as colours
- *labels* slider: choose whether to show (default) or hide labels

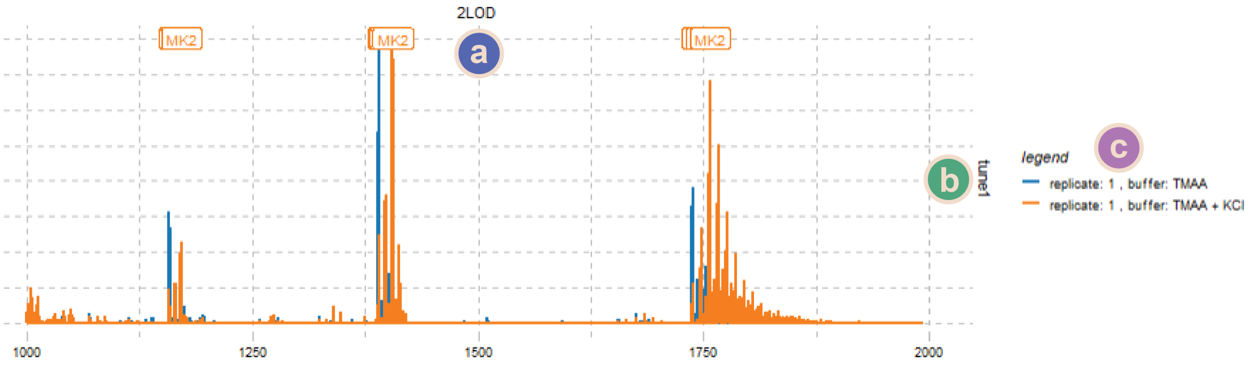

Figure S175: Detail of native MS data panelled with oligonucleotides in columns (a) and tunes in row (b). Because several spectra are superimposed, the remaining variables (replicate and buffer) are combined to map colors (c)

The charge-state focused plot *sidebar* only contains a *charge* selection menu.

The data is gathered in the *native ESI-MS data* table above, which can be sorted, filtered, and exported. The fields displayed by default are `Oligonucleotide`, `Buffer`, `Tune`, `Replicate`, `m/z`, `Normalized intensity`, and `Intensity`. The other fields hidden by default can be displayed using the *column visibility* button.

The table may take some time to load given the large number of data points.

##### 3.3.4 Reporting

**3.3.4.1 Report generation** Reports can be generated from the displayed data, either *full* (with traceability features, titles,...), or *SI* (with minimal information to avoid redundancy when reports are collated into a supporting information document), in Word, pdf, and HTML formats, in a few simple steps:

1. Select the *oligonucleotide(s)* for which the report must be generated,
2. Plot the MS data, if they are to be included in the report. If not, the section will not appear in the report,
3. Customize, where necessary, the figures (e.g. colours, scales),
4. Select the report type (*full* or *SI*) with the *Report type* switch,
5. Select a document format (Word, pdf, HTML), in the *left_sidebar* (*Report* section)
6. Click on the *Download* button and save the document.

###### 3.3.4.2 Word formatting

The Word format uses a template file to define its appearance (i.e. the styles). This template file can be changed by the user to generate reports directly with the desired appearance, to avoid additional work outside of *g4db*.

The template is located in the *markdown* folder of the *g4dbr* package. To locate the template, run:

```
system.file("rmarkdown/word-styles-reference.docx", package = "g4dbr")
```

Then, modify the **styles** as desired. Local text modifications will **not** be taken into account.

It is also advised to back up this file in another location, because any new install or update will overwrite it.

##### 3.3.5 Data deletion

It is possible to selectively remove data from the database, by oligonucleotide and analytical method, using the *database.eraser* function implemented within *g4db*.

Several oligonucleotides can be processed at once, if the same analytical methods to remove are selected. If all analytical methods are selected, the selected oligonucleotide entries will be entirely purged (including the general information).

In many cases, it is not good practice to ever delete data from a database. If the use of *g4db* lies within these cases do not use the data deletion tool as it **permanently deletes data**. Here, the data deletion tool was mostly provided as a mean to correct and update data *cleanly*, as the new data might not be written to the database if a duplicate record already exists. It is also a way to generate lighter, sub-databases for specific uses, by discarding all irrelevant entries.

By default, a new file will be generated, named `Modified database-YYYY-MM-DD.Rda`, where `YYYY-MM-DD` is the date of the day, so as to avoid accidental file overwriting.

To delete one or several entries:

1. Select the *oligonucleotide(s)* to delete from the dropdown menu in the *left_sidebar* (**not from the *general info table***),
2. Select the *methods* for which the data must be removed, by flipping the switches on,
3. Click on *Erase to a db file*
4. Save the file (with a different name than the one in use)
5. Optional: load the new database file for verification and further use

For more details on the *database.eraser* function, refer to the *Other functions and reference files* section.

#### 3.4 Importing data in the database: the *importR* tab

##### 3.4.1 Templated-Excel file

Before importing data into a database file using *g4db*, it is necessary to paste this data into a provided Excel template file. Once filled, this file doubles as a data repository that can be explored in other pieces of software. Note, however, that such files can become quite heavy (in particular with MS data), leading to very slow loading and saving times, and high memory use.

The Excel file is divided into seven tabs that contain raw data (*UV*, *CD*, *NMR*, *MS*), general oligonucleotide information (*info*), or peak labeling data (*NMR* and *MS labels*). It is essential to maintain consistency throughout the file to ensure that the data and labels are read and associated correctly: *oligonucleotide*, *electrolyte*, *cations*, *tunes* and *replicate* must be named identically across columns and tabs. If the data is to be appended to an existing database, the naming scheme must be extended to the new data. In particular, attention should be paid about capitalization (e.g. 'TMAA' vs 'tmaa' vs 'Tmaa') and typical name variants (e.g. 'Kp' vs. 'Kpi').

The template is installed with the package. Its location can be obtained by running:

```
system.file("extdata/demo_input.xlsx", package = 'g4dbr')
```

After adding data, do **not** save the file in this folder, as it would be overwritten by a package update, and deleted upon package removal.

###### 3.4.1.1 Info

The first tab gathers essential data on the entries to submit (Figure S176). Five fields must be filled, i.e.:

- `oligo`, the name of the oligonucleotide, preferably a PDB code where available,
- `sequence`, in the 5' to 3' direction, without spaces or dashes,
- `submitted_by` the initials or full name of the data submission author,
- `DOI` is the DOI of the paper linked to the PDB deposition. Paste the DOI only, and not a full link, which will be automatically generated by *importR*
- `Topology`, a short user-supplied description of the oligonucleotide structure (e.g. *parallel quadruplex*).

| oligo | sequence | submitted_by | DOI | Topology |
|---|---|---|---|---|
| 1XAV | TGAGGGTGGGTAGGGTGGGTAA | AG | 10.1021/bi048242p | Parallel |
| 2LOD | GGGATGGGACACAGGGGACGGG | AG | 10.1093/nar/gks329 | Hybrid |

Figure S176: Info template

All the other fields that can be seen in the corresponding tables in *g4db* are calculated automatically.

#### 3.4.1.2 CD

The *CD* data must be pasted in two columns, below the header, with the `wavelength` in the first column and the `ellipticity` in mdeg in the second column (Figure S177).

The `oligonucleotide`, `buffer` and `cation` names, the cuvette `path length` in cm, and the `oligonucleotide concentration` (in µM) must be supplied in the header rows.

For every new data set (new *oligonucleotide/buffer/cation* combination), the next two columns must be used and so forth. Even if the wavelength axis is the same, it must be specified again; this allows dealing with mismatched axes (see the right-hand side columns in Figure S177).

| *x* | *y* | *x* | *y* | *x* | *y* | *x* | *y* |
|---|---|---|---|---|---|---|---|
| **oligonucleotide** | 1XAV | **oligonucleotide** | 1XAV | **oligonucleotide** | 2LOD | **oligonucleotide** | 2LOD |
| **buffer** | TMAA | **buffer** | Kp | **buffer** | TMAA | **buffer** | TMAA |
| **cation** | KCl | **cation** | KCl | **cation** | | **cation** | KCl |
| **pathlength (cm)** | 0.4 | **pathlength (cm)** | 0.4 | **pathlength (cm)** | 0.4 | **pathlength (cm)** | 0.4 |
| **oligo concentration (** | 10 | **oligo concentration (** | 10 | **oligo concentration (** | 10 | **oligo concentration (** | 10 |
| **wavelength (nm)** | **CD (mdeg)** | **wavelength (nm)** | **CD (mdeg)** | **wavelength (nm)** | **CD (mdeg)** | **wavelength (nm)** | **CD (mdeg)** |
| 350 | -0.21646438 | 350 | -0.11269 | 350 | 0.076688654 | 220 | 6.868693931 |
| 349.8 | -0.101279683 | 349.8 | -0.11567 | 349.8 | 0.078271768 | 221 | 6.155448549 |
| 349.6 | 0.102572559 | 349.6 | -0.11092 | 349.6 | 0.079828496 | 222 | 5.440356201 |
| 349.4 | -0.045435356 | 349.4 | -0.10861 | 349.4 | 0.087084433 | 223 | 4.724366755 |

Figure S177: CD template. Four spectra are shown. Note that one of the x-axis is mismatched

**3.4.1.3 UV-melting** The *UV-melting* tab is the only one where three columns must be filled for each *oligonucleotide/buffer/cation* combination:

- `Temperature`, is the solution temperature, in °C or K (*importR* determines which automatically),
- `Absorbance`, is the absorbance of the solution, with or without blank subtraction (blank subtraction can be performed in *importR*)
- `Blank`, is the absorbance of the reference blank solution to subtract, where necessary.

Besides the `oligonucleotide`, `buffer` and `cation` names, the header contains a `replicate` field, to increment when several experiments for the same *oligonucleotide/buffer/cation* combination are submitted.

| *x* | *y* | *z* | *x* | *y* | *z* | *x* | *y* | *z* | *x* | *y* | *z* |
|---|---|---|---|---|---|---|---|---|---|---|---|
| **oligonucleotide** | 1XAV | | **oligonucleotide** | 1XAV | | **oligonucleotide** | 2LOD | | **oligonucleotide** | 2LOD | |
| **buffer** | TMAA | | **buffer** | Kp | | **buffer** | TMAA | | **buffer** | Kp | |
| **cation** | KCl | | **cation** | KCl | | **cation** | KCl | | **cation** | KCl | |
| **replicate** | 1 | | **replicate** | 1 | | **replicate** | 1 | | **replicate** | 1 | |
| ***Temperature*** | ***Absorbance*** | ***Blank*** | ***Temperature*** | ***Absorbance*** | ***Blank*** | ***Temperature*** | ***Absorbance*** | ***Blank*** | ***Temperature*** | ***Absorbance*** | ***Blank*** |
| 4.2 | 0.2228 | 0 | 4.2 | 0.2711 | 0 | 4.2 | 0.6109 | 0 | 4.2 | 0.4012 | 0 |
| 4.4 | 0.2225 | 0 | 4.4 | 0.2711 | 0 | 4.4 | 0.6105 | 0 | 4.4 | 0.4013 | 0 |
| 4.6 | 0.2227 | 0 | 4.6 | 0.2715 | 0 | 4.6 | 0.6091 | 0 | 4.6 | 0.4013 | 0 |
| 4.8 | 0.2227 | 0 | 4.8 | 0.2717 | 0 | 4.8 | 0.6112 | 0 | 4.8 | 0.4015 | 0 |
| 5 | 0.2228 | 0 | 5 | 0.2718 | 0 | 5.1 | 0.6127 | 0 | 5 | 0.4017 | 0 |
| 5.3 | 0.2233 | 0 | 5.3 | 0.2717 | 0 | 5.3 | 0.6119 | 0 | 5.3 | 0.4022 | 0 |
| 5.5 | 0.2226 | 0 | 5.5 | 0.2712 | 0 | 5.5 | 0.6119 | 0 | 5.5 | 0.4005 | 0 |

Figure S178: UV-melting template. Case where the data is already blank-subtracted

The melting data must be pasted as is, in particular if both cooling and heating ramps are recorded successively. *MeltR* uses the changes in temperature (increase or decrease) from successive rows to assess whether it deals with a heating or cooling ramps, and eventually dissociates both for further processing.

**3.4.1.4 NMR** The *$^{1}$H-NMR* template follows the same principle as the *CD* one: two columns (per *oligonucleotide/buffer/cation* combination) for the chemical shift and intensity, and three header rows for the `oligonucleotide`, `buffer` and `cation` names (Figure S179).

| *x* | *y* | *x* | *y* |
|---|---|---|---|
| **oligonucleotide** | 1XAV | **oligonucleotide** | 2LOD |
| **buffer** | TMAA | **buffer** | TMAA |
| **cation** | KCl | **cation** | KCl |
| **Chemical shift** | **Intensity** | **Chemical shift** | **Intensity** |
| 14.61938 | -5112 | 14.61865 | 22577 |
| 14.61908 | -5136 | 14.61835 | 22475 |
| 14.61877 | -5159 | 14.61804 | 22363 |
| 14.61847 | -5173 | 14.61774 | 22240 |
| 14.61817 | -5177 | 14.61744 | 22110 |
| 14.61787 | -5167 | 14.61713 | 21977 |
| 14.61756 | -5151 | 14.61683 | 21839 |
| 14.61726 | -5136 | 14.61653 | 21692 |
| 14.61696 | -5124 | 14.61623 | 21536 |

Figure S179: NMR template

**3.4.1.5 NMR labels** This tab is used to submit $^{1}$H NMR peak labelling information (Figure S180). The header structure is the same than in the NMR data tab. The first column must be filled with `peak` numbers, in any order, with the corresponding `chemical shifts` in the second column. The labels are handled as text, and therefore several numbers can be submitted for a single chemical shift value.

As a sidenote, it is possible to keep cells empty if a given peak number is in the list but there is no corresponding peak in the spectrum. This is practical when several spectra are being labelled and a common peak number list is used. Note that the peak list must be repeated for all spectra, even if they are identical.

| *x* | *y* | *x* | *y* |
|---|---|---|---|
| **oligonucleotide** | 1XAV | **oligonucleotide** | 2LOD |
| **buffer** | TMAA | **buffer** | TMAA |
| **cation** | KCl | **cation** | KCl |
| **Peaks** | **chemical shift** | **Peaks** | **chemical shift** |
| 9 | 10.55 | 3 | 10.96 |
| 18 | 10.96 | 14 | 10.98 |
| 13,22 | 10.98 | 8 | 11.04 |
| 8 | 11.16 | 22 | 11.22 |
| 17 | 11.19 | 15 | 11.3 |
| 20 | 11.22 | 6,7 | 11.52 |
| 21 | 11.26 | 16 | 11.53 |
| 12 | 11.42 | 2 | 11.55 |
| 11 | 11.625 | 1 | 11.57 |
| 7 | 11.65 | 20 | 11.71 |
| 16 | 11.85 | 21 | 11.85 |

Figure S180: NMR labels template. Note that both oligonucleotides have completely different labellings

Make sure to mirror the header from the NMR data tab, so that all spectra are labelled.

**3.4.1.6 MS** The MS template shares the same structure as *NMR* and *CD*, with `m/z` and the `intensity` as columns one and two (Figure S181). The intensity can be supplied normalized or not, it will eventually be normalized in *importR*. Two additional header rows must be filled:

- `tune`, a short name identifying the MS parameters. The name must be linked to said parameters along the database file (e.g. publication, readme file).
- `replicate`, a number to increment when several experiments for the same *oligonucleotide/buffer/cation/tune* combination are submitted.

| *x* | *y* | *x* | *y* | *x* | *y* | *x* | *y* |
|---|---|---|---|---|---|---|---|
| **oligonucleotide** | 1XAV | **oligonucleotide** | 1XAV | **oligonucleotide** | 2LOD | **oligonucleotide** | 2LOD |
| **buffer** | TMAA | **buffer** | TMAA | **buffer** | TMAA | **buffer** | TMAA |
| **cation** | | **cation** | KCl | **cation** | | **cation** | KCl |
| **tune** | tune1 | **tune** | tune1 | **tune** | tune1 | **tune** | tune1 |
| **replicate** | 1 | **replicate** | 1 | **replicate** | 1 | **replicate** | 1 |
| **m/z** | **Intensity** | **m/z** | **Intensity** | **m/z** | **Intensity** | **m/z** | **Intensity** |
| 299.9195 | 0 | 299.9195 | 167 | 299.9195 | 30 | 299.9195 | 150 |
| 299.9225 | 1 | 299.9225 | 163 | 299.9225 | 39 | 299.9225 | 148 |
| 299.9255 | 5 | 299.9255 | 145 | 299.9255 | 30 | 299.9255 | 141 |
| 299.9285 | 0 | 299.9285 | 155 | 299.9285 | 16 | 299.9285 | 179 |
| 299.9314 | 10 | 299.9314 | 223 | 299.9314 | 11 | 299.9314 | 174 |
| 299.9344 | 38 | 299.9344 | 288 | 299.9344 | 24 | 299.9344 | 123 |
| 299.9374 | 72 | 299.9374 | 273 | 299.9374 | 22 | 299.9374 | 97 |

Figure S181: MS template

It is advised to be relatively conservative with data-heavy spectra to cut on processing time in *importR*, e.g. irrelevant m/z ranges can be discarded. In case of doubt, everything can be kept at this stage and filtered later on in *importR*.

###### 3.4.1.7 MS labels

This tab is aimed at providing the database with the **nature** of the species to label in the MS spectrum, not their `m/z`. It therefore differs from the NMR label tab, where one must supply the `chemical shift` of each label.

The first column contains the charge state numbers, to label different charge states independently (Figure S182). The second column contains the name of the species to be labelled, which must be supplied using the following syntax: *M* for the non-adducted oligonucleotide, *MK* for a single-potassium-adduct species, *MK2* for a two-potassium-adduct species, and so forth (up to ten).

| *x* | *y* | *x* | *y* | *x* | *y* | *x* | *y* |
|---|---|---|---|---|---|---|---|
| **oligonucle** | 1XAV | **oligonucle** | 1XAV | **oligonucle** | 2LOD | **oligonucle** | 2LOD |
| **buffer** | TMAA | **buffer** | TMAA | **buffer** | TMAA | **buffer** | TMAA |
| **cation** | | **cation** | KCl | **cation** | | **cation** | KCl |
| **tune** | tune1 | **tune** | tune1 | **tune** | tune1 | **tune** | tune1 |
| **replicate** | 1 | **replicate** | 1 | **replicate** | 1 | **replicate** | 1 |
| **charge** | **label** | **m/z** | **Intensity** | **charge** | **label** | **m/z** | **Intensity** |
| 4 | M | 4 | MK2 | 4 | M | 4 | M |
| 5 | M | 5 | MK2 | 5 | M | 4 | MK |
| 6 | M | 6 | MK2 | 6 | M | 4 | MK2 |
| | | | | | | 5 | M |
| | | | | | | 5 | MK |
| | | | | | | 5 | MK2 |
| | | | | | | 6 | M |
| | | | | | | 6 | MK |
| | | | | | | 6 | MK2 |

Figure S182: MS labels template. Note the difference in labelling between oligonucleotides and buffer.

Make sure to mirror the header from the MS data tab, so that all spectra are labelled.

##### 3.4.2 Populating a database

Once the template file is ready, the data can be loaded in *g4db*, processed, filtered, and written into a new or existing database file. All of these steps can be performed in the *importR* tab, except for the *UV-melting* data treatment that is carried out in *meltR* (see the Importing UV-melting data: the *meltR* tab section).

Essentially, *importR* works just like *database*. The main window hosts the same data tables and figures than database (except *UV-melting* figures, which are in *meltR*, and the charge-focused MS plot), with the same functioning (data filtering, figure customization). In the same vein, the *left sidebar* also contains the filters and color palette selection menus. All these common features are described in the *Interface overview* and *Consulting a database: the database tab* sections, and will not be discussed below.

The key aspect of *importR* is that it is a *selective* database writing tool. In that context:

- **What you see is what you write** to the database. Any data point filtered out (whether by *oligonucleotide*, *buffer* compositon, x-axis range), will *not* be written in the database file.
- Duplicated data points (same technique, *oligonucleotide*, *buffer* composition, x-axis position,...) are discarded. For instance, resubmitting data with a wider x-axis range will have the effect of completing the

database (without doubling the already existing points), but resubmitting corrected data on the same range might not replace the initial data. It is therefore better to first remove the erroneous entry (see the Data deletion section).
- Individual oligonucleotides and analytical methods can be included or excluded from the database writing.

###### 3.4.2.1 Template file input

The data is imported by selecting a file via the *Browse…* button in the *left sidebar*.

###### 3.4.2.2 Data filtering and processing

Oligonucleotides are selected from the *General information* table. Further buffer composition filtering can be performed in the *left sidebar*.

The *CD* and *NMR* calculations (e.g. normalization, labeling) and plotting are automatically performed, without any user input. The MS data is processed and plotted when the *plot MS* button is clicked. Note that if the MS data is not plotted, it cannot be exported to a database.

Method-dependent *filtering* is performed in the corresponding *right sidebars*, as described for the *database* tab.

###### 3.4.2.3 Importing UV-melting data: the *meltR* tab

The processing of UV-melting data is performed in *meltR*, a distinct tab from *importR*, to avoid overcrowding the interface and allow its use outside of the database frame.

The data is sourced from the template file loaded in *importR*, and once the data is processed in *meltR* it can be sent back to *importR* to include in the database. Note that the filtering of temperature range and buffer composition must be performed directly in *meltR*.

The use of *meltR* itself is described below.

##### 3.4.3 Writing a database file

Once the data has been selected and properly filtered (including or not UV-melting data from *meltR*), it can be written into a database file in three simple steps:

1. Select a database file, either an existing one (to add new entries) or an empty one (to create a new database). This file can be opened in the *Export* section of the *left sidebar* of *importR*, or from the *database* tab. In either way, the data can be consulted in the *database* tab. An empty file is available in the package, and can be found by running:

```
system.file("extdata/empty_database.Rda", package = 'g4dbr')
```

2. Select the methods to write to the database file, using the switches. The MS and UV-melting data must be generated to be exported.
3. Click on *Write to db file*. By default, the file will be named following the `Database-YYYY-MM-DD.Rda` template. Rename where necessary. If the database in use was generated the same day than the deletion operation, there is a risk of it being overwritten: make sure to name the new file with a different name.
4. Optional: load the new/updated database to verify that the import worked correctly.

#### 3.5 Automated processing of UV-melting data: the *meltR tab*

##### 3.5.1 Principle

###### 3.5.1.1 Purpose

*meltR* is an automated UV-melting data processing software. It determines the melting temperatures ($T_m$), $\Delta G^0$, $\Delta H^0$ and $\Delta S^0$ by non linear fitting, and converts the absorbances into folded fractions.

Folded fractions are a good way to assess to which extent an oligonucleotide is structured (1: all molecules folded, 0: all molecules unfolded), visually observe the $T_m$ (folded fraction = 0.5), and normalize the data of different samples (and therefore different absorbances) to a common y-scale.[3]

For the non-linear fitting and the folded fraction calculation to work, the data must contain both a *lower* and *higher* baseline.[3] In other words, the oligonucleotide must not be too stable or too unstable. In such cases, *meltR* allows to normalize the data to [0;1] to at least bring all data to a common y-scale.

**3.5.1.2 Data modeling: General model** In a melting experiment, changes in the solution temperature lead to changes in the amount of folded (decreases with increasing temperatures) and unfolded species (increases with increasing temperatures). The model relies on the expression of the measured absorbance $A_T$ as the sum of the absorbances from the folded (*F*) and unfolded (*U*) forms, weighted by their abundance expressed from the folded fraction $\theta_T$.

$$A_T = A_T^F \times \theta_T + A_T^U \times (1 - \theta_T)$$

Herein, the absorbances measured at 295 nm were converted to molar extinction coefficient (in $M^{-1}cm^{-1}$) using $\varepsilon = A/lC$, where $l$ is a path length (in cm) and $C$ the oligonucleotide concentration (in M).

$$\varepsilon_T = \varepsilon_T^F \times \theta_T + \varepsilon_T^U \times (1 - \theta_T)$$

The folded fraction is defined by $\theta = \frac{[F]}{[F]+[U]}$. Assuming a simple two-state model $F \Leftrightarrow U$ with an equilibrium constant $K$, $\theta$ can be expressed as:

$$\theta = \frac{1}{1+K}$$

This leads to:

$$\varepsilon_T = \varepsilon_T^F \times \frac{1}{1+K} + \varepsilon_T^U \times \frac{K}{1+K}$$

$\varepsilon_T^F$ and $\varepsilon_T^U$ can be modeled as a linear function of the temperature, where $a$ is the slope and $b$ the intercept of these baselines:

$$\varepsilon_T = (a^F T + b^F) \times \frac{1}{1+K} + (a^U T + b^U) \times \frac{K}{1+K}$$

$K$ can be expressed by thermodynamic quantities of interest: $\Delta G^0$, $\Delta H^0$ and $\Delta S^0$.

$$-RTlnK = \Delta G^0 = \Delta H^0 - T\Delta S^0$$

Note that in meltR, potential changes in heat capacity changes in the evaluated temperature range are not taken into account to avoid over-paramaterization. At the melting temperature:

$$\Delta G_m^0 = \Delta H_m^0 - T\Delta S_m^0 = 0$$

Which leads to:

$$\Delta S_m^0 = \frac{\Delta H_m^0}{T_m}$$

And therefore:

$$\Delta G^0 = \Delta H_m^0(1 - \frac{T}{T_m})$$

Finally, K can be expressed as $exp(-\frac{\Delta H^0(1-\frac{T}{T_m})}{RT})$, yielding:

$$A_T = (a^F T + b^F) \times \frac{1}{1 + exp(-\frac{\Delta H^0(1-\frac{T}{T_m})}{RT})} + (a^U T + b^U) \times \frac{exp(-\frac{\Delta H^0(1-\frac{T}{T_m})}{RT})}{1 + exp(-\frac{\Delta H^0(1-\frac{T}{T_m})}{RT})}$$

###### 3.5.1.3 Data modeling: Implementation and derived values

In meltR, the absorbance is converted to molar extinction coefficients before fitting with the following model:

```
#code simplified for readibility
epsilon = (P3+P4*T)*1/(1+exp(-P1*(1-T/P2)/(8.31451*T))) +
  (P5+P6*T)*exp(-P1*(1-T/P2)/(8.31451*T))/(1+exp(-P1*(1-T/P2)/(8.31451*T)))
```

where `epsilon` is the molar extinction coefficient, `T` is the temperature (in Kelvin), `P1` is $\Delta H^0$, `P2` is the $T_m$, `P3`/`P5` and `P4`/`P6` are respectively the origins and slopes of the baselines. The optimized parameters are summarized in the *meltR* tab, and can be later consulted in the *database* tab.

The non-linear fitting is performed with the base function `nls()`. Below is a more detailed view of the fitting model, applied on a demo data melting curve of *1XAV* (Figure S183). Note that some user inputs have been hard-coded hereafter:

```
#libraries
library(tidyverse)
library(ggthemes)

#Experimental conditions
melt.c <- 10 #oligo concentration (micromolars)
melt.l <- 1 #cuvette of 1.0-cm path length

#loading the demo data
load(system.file('extdata/demo_database.Rda', package = 'g4dbr'))

#Selection of a melting curve from the demo data
data.to.fit <- db.UV %>%
  select(oligo, buffer, cation, rep, comment, ramp, id, T.K, abs.melt) %>%
  filter(oligo == '1XAV' & buffer == '100 mM TMAA (pH 7.0)' & ramp == 'cooling')

#Plot
data.to.fit %>%
  ggplot() +
  geom_point(aes(x = T.K, y = abs.melt), color = 'steelblue') +
  theme_pander() +
  xlab("T (K)") +
  ylab(expression(epsilon~(M^-1*cm^-1)))
```

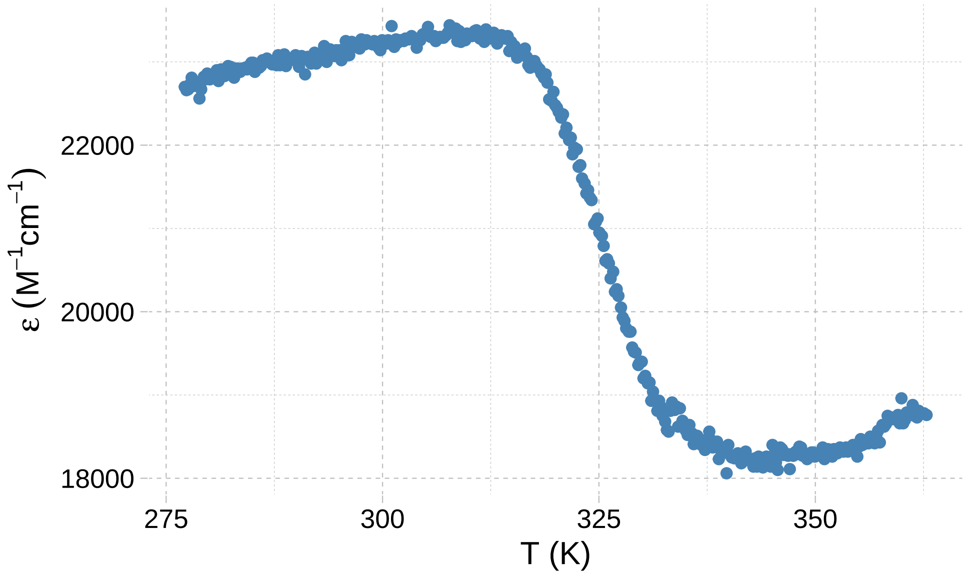

Figure S183: Melting curve of 1XAV (cooling ramp) in 100 mM TMAA + 1 mM KCl, from the demo database

```
#Fit initialization (automated in the application)
P1s <- 130000
P2s <- 325 #automatically extracted from the first derivative in the application
P3s <- 1/(melt.c/1E6 * melt.l) #denominator converts initial parameters to molar abs coeff.
P4s <- 0.30/(melt.c/1E6 * melt.l)
P5s <- 0/(melt.c/1E6 * melt.l)
P6s <- -0.2/(melt.c/1E6 * melt.l)

#Non-linear fitting using the base nls() function
ms <- nls(
  data=data.to.fit,
  data.to.fit$abs.melt~(P3+P4*data.to.fit$T.K)*1 /
    (1+exp(-P1*(1-data.to.fit$T.K/P2) / (8.31451*data.to.fit$T.K))) +
    (P5+P6*data.to.fit$T.K)*exp(-P1*(1-data.to.fit$T.K/P2) / (8.31451*data.to.fit$T.K))
  / (1+exp(-P1*(1-data.to.fit$T.K/P2) / (8.31451*data.to.fit$T.K))),
  start = list(P1 = P1s, P2 = P2s, P3=P3s, P4=P4s, P5=P5s, P6=P6s), #initial parameters
  nls.control(maxiter = 5000, #default value, hard-coded here but users can modify it
              warnOnly = T)
)

#Optimized parameters
fit.output <- data.frame(
  nb.data.pt = nobs(ms),
  RSS = sum(residuals(ms)^2),
  SE.residual = sigma(ms),
  P1 = as.vector(coef(ms))[1],
  P2 = as.vector(coef(ms))[2],
  P3 = as.vector(coef(ms))[3],
  P4 = as.vector(coef(ms))[4],
  P5 = as.vector(coef(ms))[5],
  P6 = as.vector(coef(ms))[6]
```

```
)

fit.output
#>   nb.data.pt     RSS SE.residual       P1       P2       P3       P4       P5       P6
#> 1        387 2555867    81.90429 218057.5 325.7146 16737.21 21.67566 8193.128 28.99342
```

Note that the residual sum of squares (RSS) and standard error of residuals (RMSE) are computed.

After the fitting is complete, a number of derived values are calculated. The $\Delta H$ř and $\Delta S$ř of the folding reaction are obtained from `P1` and `P2`.

```
#Temperature at which the free energy is calculated
temp = 293 #User input in the app

DeltaH = -as.vector(coef(ms))[1]
DeltaS = -as.vector(coef(ms))[1]/as.vector(coef(ms))[2]
DeltaG = DeltaH - temp * DeltaS

data.frame(DeltaH, DeltaS, DeltaG)
#>      DeltaH   DeltaS    DeltaG
#> 1 -218057.5 -669.474 -21901.57
```

The baselines (in $M^{-1}cm^{-1}$) are obtained with `P3+P4*T` and `P5+P6*T` (Figure S184):

```
data.to.fit %>%
  mutate(low.T.baseline = fit.output$P3+fit.output$P4*T.K, #low temperature baseline
         high.T.baseline = fit.output$P5+fit.output$P6*T.K) %>% #high temperature baseline
  ggplot() +
  geom_point(aes(x = T.K, y = abs.melt), color = 'steelblue') +
  geom_line(aes(x = T.K, y = low.T.baseline), color = "coral", size = 1) +
  geom_line(aes(x = T.K, y = high.T.baseline), color = "coral", size = 1) +
  theme_pander() +
  xlab("T (K)") +
  ylab(expression(epsilon~(M^-1*cm^-1)))
```

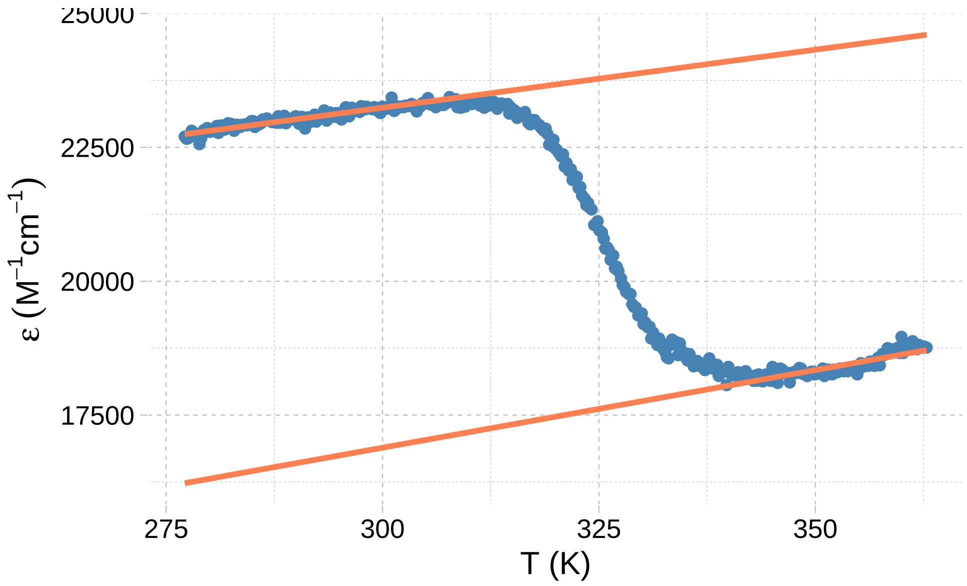

Figure S184: The baselines are not determined manually, but computed from the fitting parameters

The folded fraction (Figure S185) is calculated by deconvoluting the baseline contributions:

$$\theta = \frac{P6T + P5 - \varepsilon}{P6T + P5 - (P4T + P3)}$$

```
data.to.fit %>%
  mutate(folded.fraction.model = (fit.output$P5+fit.output$P6*T.K-abs.melt)/(fit.output$P5+fit.output$P6
  ggplot(aes(x = T.K, y = folded.fraction.model)) +
  geom_point(color = "steelblue") +
  theme_pander() +
  xlab("T (K)") +
  ylab(expression(epsilon~(M^-1*cm^-1)))
```

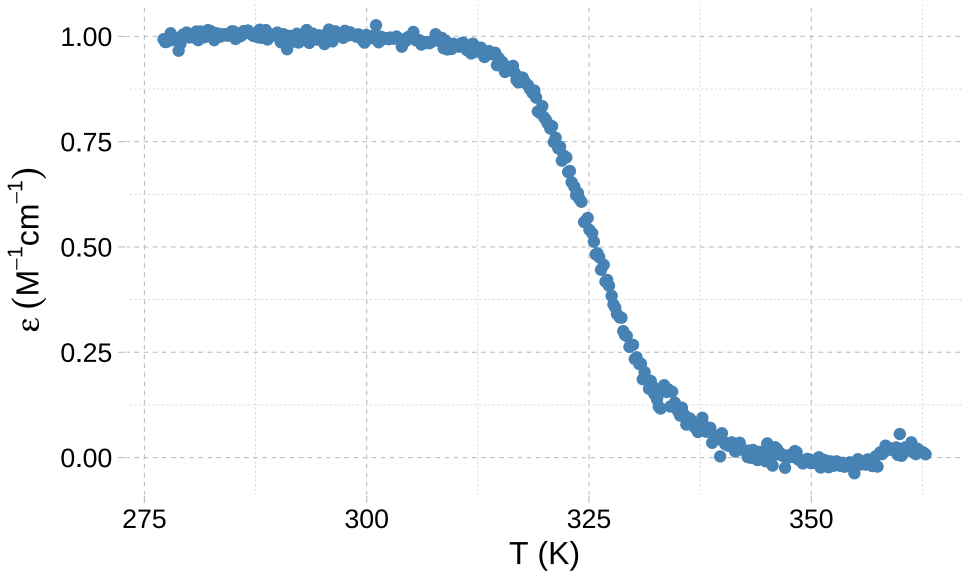

Figure S185: The folded fraction of 1XAV (cooling ramp) in 100 mM TMAA + 1 mM KCl

The modeled folded fraction (Figure S186) is also derived from the fit, using:

$$\theta_{model} = \frac{1}{1 + exp(-\frac{P1(1-\frac{T}{P2})}{RT})}$$

.

```
data.to.fit %>%
  mutate(folded.fraction =
           (1/(1+exp(-fit.output$P1*(1-T.K/fit.output$P2)/(8.31451*T.K))))) %>%
  ggplot(aes(x = T.K, y = folded.fraction)) +
  geom_point(color = "steelblue") +
  theme_pander() +
  xlab("T (K)") +
  ylab(expression(epsilon~(M^-1*cm^-1)))
```

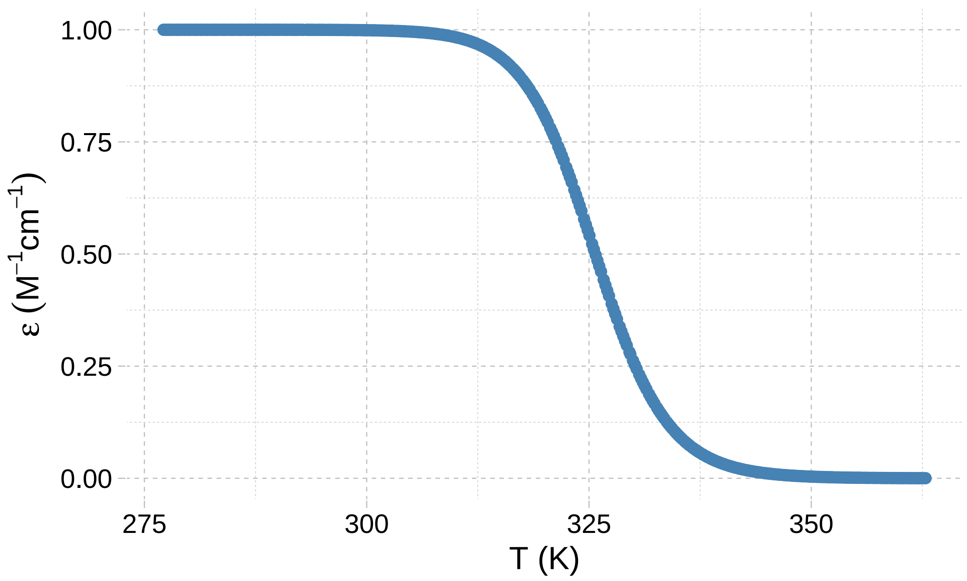

Figure S186: The modeled folded fraction of 1XAV (cooling ramp) in 100 mM TMAA + 1 mM KCl

**3.5.1.4 Workflow** The data is processed following this workflow:

1. Detection of the temperature unit, and conversion to Kelvin where necessary,
2. Generation of a unique `id` for each *oligonucleotides*, *ramps*, *buffers*, and *replicates* combinations. From then on, all data is processed by `id` (in particular cooling and heating ramps are processed separately).
3. Blank subtraction, if blank data is submitted (can be turned off),
4. Conversion of the absorbance data to molar extinction coefficient,
5. Determination and separation of the ramps (cooling and heating). The ramps are always processed separately.
6. Data selection from user input: *oligonucleotides*, *ramps*, *buffers*, *replicates*, or individual *id*.

The steps 7–9 are only carried out if the data can be fitted (presence of both lower and upper baselines):

7. Non linear fitting initialization
   a. `P2` (the *Tm*) is initialized as the maximum of the first derivative ($\frac{\Delta\varepsilon}{\Delta T}$)
   b. All other parameters initial values are hard-coded, and modulated by the oligonucleotide concentration and cell path length
   c. User modifications, where necessary
8. Non linear fitting (see model above),
9. Calculation of the folded fractions (from experimental data and from the model) and baselines (see equations above)

Step 10 is only carried out for non-fittable data:

10. The $\varepsilon$ values are normalized in the [0;1] range, to be displayed alongside folded fraction data (same y-scale).

##### 3.5.2 Data loading and filtering

The data must be loaded from the Excel template into *importR*. All of the UV-melting data is automatically imported into *meltR*, regardless of the oligonucleotides selected in *importR* (to facilitate the standalone use). However, only the processed data for the oligonucleotides selected in *importR* is sent back to that tab.

The *meltR* interface has a slightly different organization than *importR* and *database*: the filtering of data to process is carried out in the hovering *Filter* panel (Figure S187).

1. Where necessary, refine the temperature range (default: 276-363 K, or ~ 3-90 °C),
2. Select the oligonucleotides to process (default to all). It is possible to process several oligonucleotides at once. Remember however that, in the context of *g4db*, these different oligonucleotides need to be selected in *importR* to be sent to that tab.
3. Select the ramps (heating or cooling) to process (default: both). The nature of the ramps is determined automatically, and the ramps are processed separately.
4. Select the buffers to process (default: all),
5. Select the replicates to process (default: all)
6. If the steps 2–5 do not allow to specifically select the desired data, it is possible to directly filter the data by `id`.

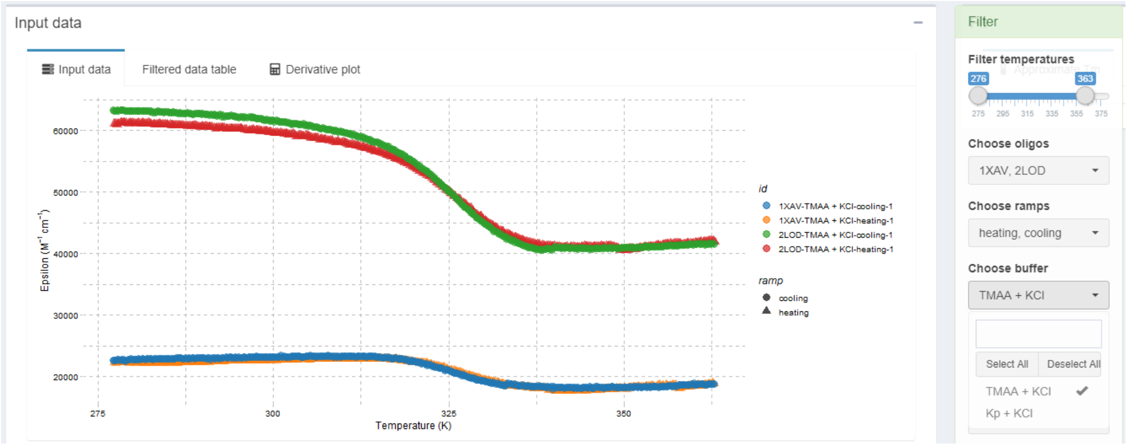

Figure S187: The UV-melting data from the demo input, where the Kp+KCl buffer was filtered off

The *Filter* panel can be minimized by clicking on the header.

##### 3.5.3 Data fitting

This section can be carried out only for data that can be fitted. For non-fittable data, skip this section.

1. Click on the *Plot derivative* button, located in the *left sidebar*.
   a. The *Input data* box will automatically switch to display $\frac{\Delta\varepsilon}{\Delta T}$ (Figure S188)
   b. The *Approximate Tm* table is filled with the maxima from the derivatives, in the *Fit* box.

   c. Artifactual points (e.g. caused by important local data variations) may lead to erroneous approximated *Tm*: increase the smooth window and click on the button again. If the results are still not satisfactory, continue anyway to step 2 (Figures S188 and S189).

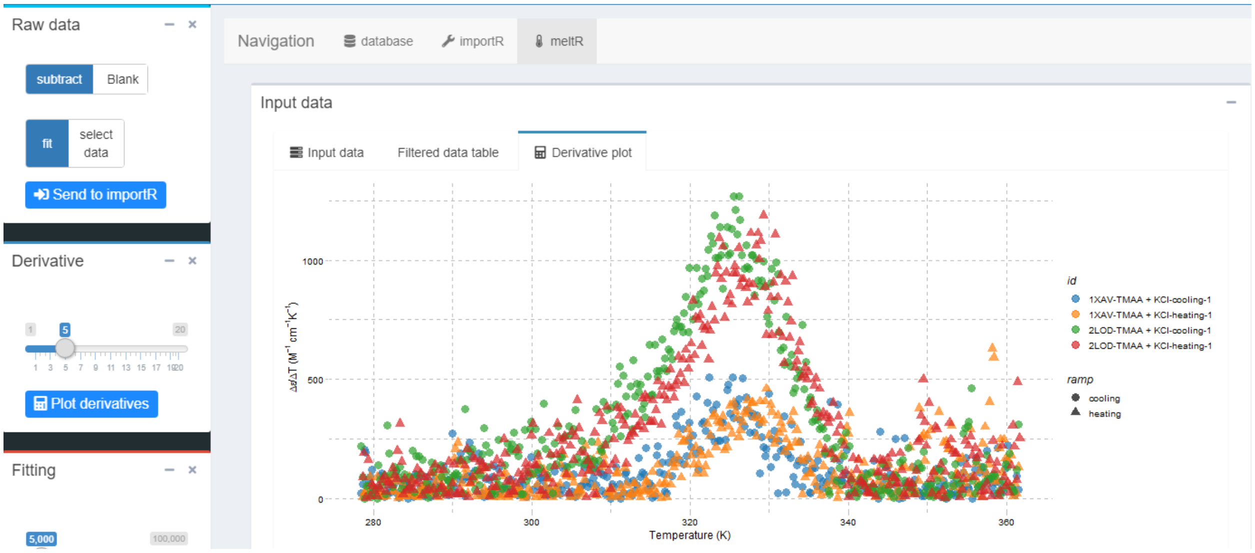

Figure S188: First derivative data was obtained by clicking on Plot derivatives. Note the presence of artifacts at high temperature that will cause an erroneous initialization to the Tm for 1XAV-TMAA + KCl-heating-1

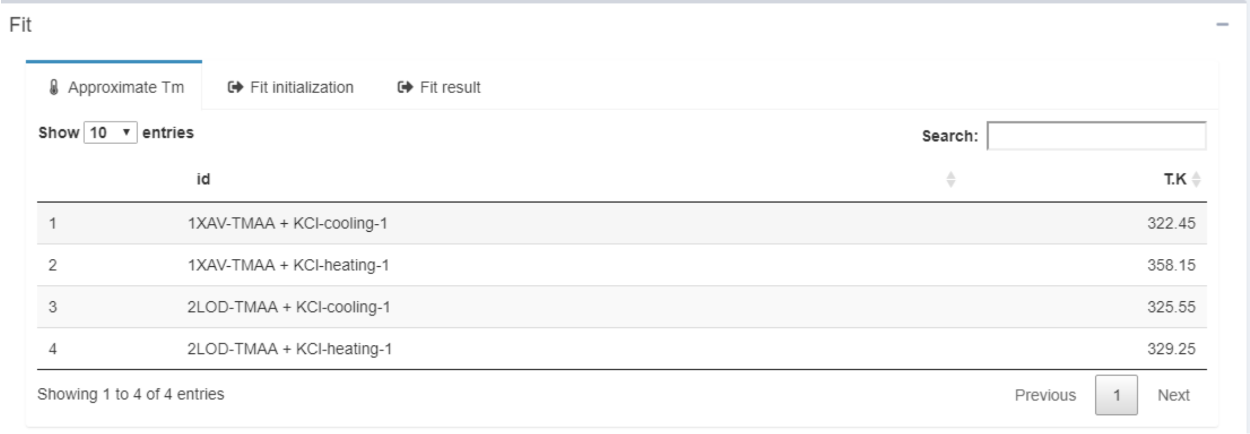

| | id | T.K |
|---|---|---|
| 1 | 1XAV-TMAA + KCl-cooling-1 | 322.45 |
| 2 | 1XAV-TMAA + KCl-heating-1 | 358.15 |
| 3 | 2LOD-TMAA + KCl-cooling-1 | 325.55 |
| 4 | 2LOD-TMAA + KCl-heating-1 | 329.25 |

Figure S189: Tm initialization from first derivative data. Here, the second entry is erroneous and must be corrected either by increasing the derivative smoothing, or manually at the next step

2. Click on the *Initialize fitting button*, located in the *left sidebar* (Figure S190).
   a. The *Fit* box will automatically switch to the *Fit initialization* table.
   b. If step 1. was not satisfactory, manually correct the `Tm.init` variable. Correctly initialized *Tm* are critical for the success of the fitting process. The other initial fitting parameter values can also be modified.
   c. If desired, change the legend; by default it is the `id`

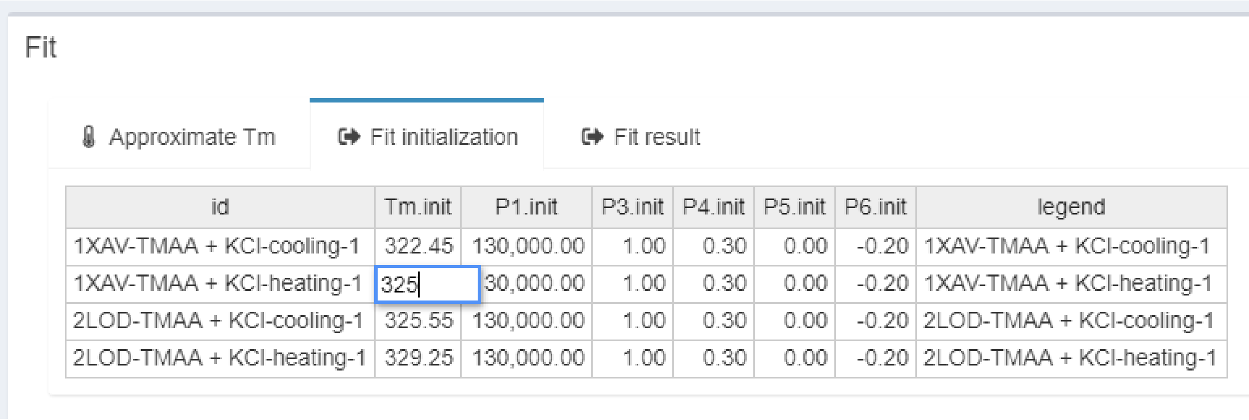

| id | Tm.init | P1.init | P3.init | P4.init | P5.init | P6.init | legend |
|---|---|---|---|---|---|---|---|
| 1XAV-TMAA + KCl-cooling-1 | 322.45 | 130,000.00 | 1.00 | 0.30 | 0.00 | -0.20 | 1XAV-TMAA + KCl-cooling-1 |
| 1XAV-TMAA + KCl-heating-1 | 325 | 30,000.00 | 1.00 | 0.30 | 0.00 | -0.20 | 1XAV-TMAA + KCl-heating-1 |
| 2LOD-TMAA + KCl-cooling-1 | 325.55 | 130,000.00 | 1.00 | 0.30 | 0.00 | -0.20 | 2LOD-TMAA + KCl-cooling-1 |
| 2LOD-TMAA + KCl-heating-1 | 329.25 | 130,000.00 | 1.00 | 0.30 | 0.00 | -0.20 | 2LOD-TMAA + KCl-heating-1 |

Figure S190: Fitting initialization. All parameters are initialized. Note that the Tm initialization is being manually corrected

3. Click on the *Launch fitting* button, and the data will be processed and the result displayed in several figures and tables (Figure S191).
   a. The *Fit* box will automatically switch to the fit result tab, showing the fit lines and calculated baselines. Baselines can be toggled off using the corresponding switch in the *left sidebar*.
   b. The folded fractions (and modeled folded fraction) are shown in the *Fit results* box
   c. The melting temperatures and other thermodynamic values are accessible in the *Melting temperatures* box. The temperature at which the Free energy is calculated can be adjusted from a slider in the *left sidebar*. The *Tm* values are also plotted in the *Plot* tab (box plot grouped by *oligonucleotide* and *buffers*, with distinctive colors per ramp).
   d. If the fit fails, it is likely that the data was not correctly initialized. Change the parameters, and click again on *Launch fitting*.
   e. Where necessary, the maximum number of iterations can be increased (slider in the *left sidebar*; default: 5000).

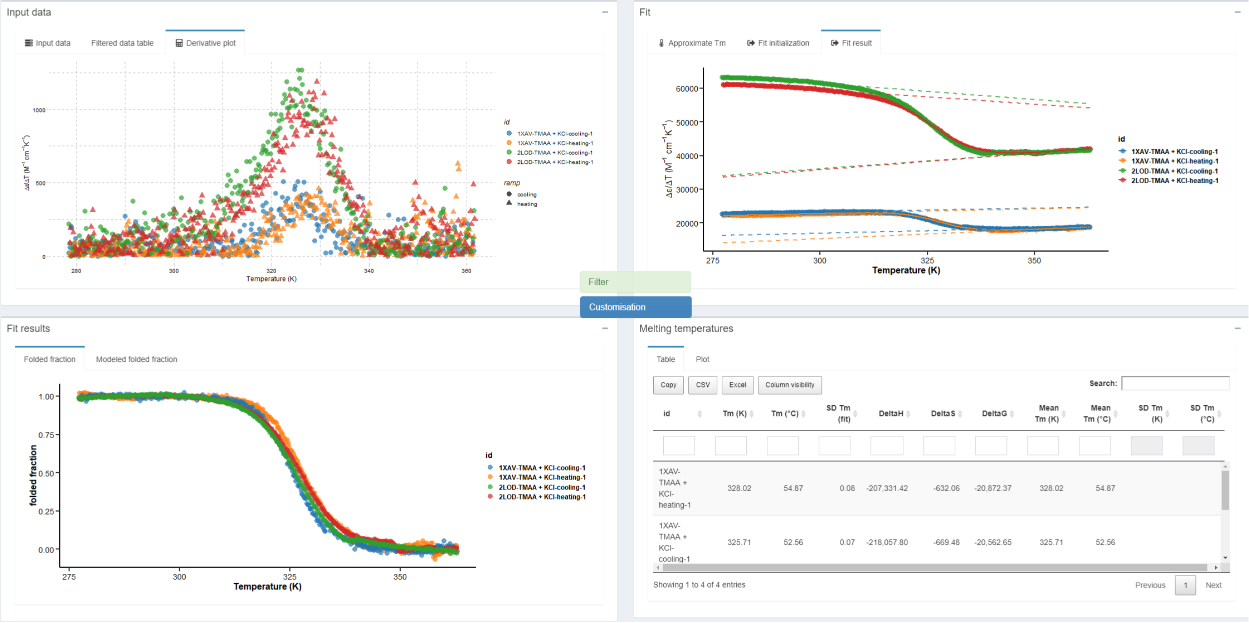

Figure S191: Fitting results: Fitted data (top right), folded fraction (bottom left), data table and Tm plot (bottom right)

###### 3.5.4 Sending data to *importR*

To send data to *importR* for database edition:

1. If not already done, select the oligonucleotides to import in *importR* from the *General information* table of that tab,
2. Select whether the data was fitted or not with the *select data* switch, in the *left sidebar*,
3. Click on the *send to importR* button,
4. In *importR*, verify that the data has correctly been sent into the *UV-melting data* box.

###### 3.5.5 Figure customization

The choice of colour palettes, lines and points size and transparency, can be made from the hovering *Customisation* panel. The panel can be minimized by clicking on the header.

### 4 Other functions and reference files

#### 4.1 epsilon.calculator

###### 4.1.1 Principle

The oligonucleotide molar extinction coefficients at 260 nm are calculated using the nearest-neighbor model in its traditional format,[1,2] where $\varepsilon_i$ is the molar extinction coefficient (in M$^{-1}$cm$^{-1}$) of the nucleotide in position $i$ (in the 5'to 3' direction), $\varepsilon_{i,i+1}$ is the extinction coefficients for doublets of nucleotides in positions $i$ and $i+1$, and $N_b$ is the number of nucleotides in the oligonucleotide.

$$\varepsilon_{260nm} = \sum_{i=1}^{N_b-1} \varepsilon_{i,i+1} - \sum_{i=2}^{N_b-1} \varepsilon_i$$

To that effect, it uses `epsilondb`, a database of reference $\varepsilon_{260nm}$ contributions from the individual nucleobases, and couples of nucleobases (neighboring effects):

```
epsilondb
#> # A tibble: 4 x 6
#>   base  epsilon Acorr Ccorr Gcorr Tcorr
#>   <chr>   <dbl> <dbl> <dbl> <dbl> <dbl>
#> 1 A       15400 27400 21200 25000 22800
#> 2 C        7400 21200 14600 18000 15200
#> 3 G       11500 25200 17600 21600 20000
#> 4 T        8700 23400 16200 19000 16800
```

`epsilondb` is contained within the `[installpath]/data/Rdata.rds` file after the package is built. The value may be modified from the `[installpath]/extdata/referencedb.xlsx` but requires to rebuild the package.

###### 4.1.2 Code

The code of `epsilon.calculator` is contained in `R/EpsilonCalc.R`

First, the list of nucleobases and their nearest 3' neighbor are extracted from the user-supplied sequence (here 5'-GCAT-3'):

```r
library(stringr)
library(tidyverse)

#sequence provided by the user
sequence <- 'GCAT'

#initialization of result data frame
epsilon.calc <- data.frame()
buffer <- data.frame()
result <- data.frame()

#extraction of individual bases and their 3' nearest neighbor
for (i in 1:str_length(sequence)) {
  buffer <- data.frame(position = i,
                       nucleo = substr(sequence, i, i),
                       nn = substr(sequence, i+1, i+1)
  )
  epsilon.calc <- rbind(epsilon.calc, buffer)
}

epsilon.calc
#>   position nucleo nn
#> 1        1      G  C
#> 2        2      C  A
#> 3        3      A  T
#> 4        4      T
```

Their contribution are then attributed by matching their one letter code to the database, and both the 5' and 3' ends have their individual contributions set to zero.

```r
#attribution of individual and nearest neighbor contributions
  epsilon.calc <- epsilon.calc %>%
    mutate(
      indiv.base.cont = case_when( #individual
        nucleo == 'G' ~ epsilondb$epsilon[epsilondb$base == 'G'],
        nucleo == 'C' ~ epsilondb$epsilon[epsilondb$base == 'C'],
        nucleo == 'T' ~ epsilondb$epsilon[epsilondb$base == 'T'],
        nucleo == 'A' ~ epsilondb$epsilon[epsilondb$base == 'A']
      ),
      nn.cont = case_when( #nearest neighbor
        nucleo == 'G' ~ case_when(
          nn == 'G' ~ epsilondb$Gcorr[epsilondb$base == 'G'],
          nn == 'C' ~ epsilondb$Ccorr[epsilondb$base == 'G'],
          nn == 'T' ~ epsilondb$Tcorr[epsilondb$base == 'G'],
          nn == 'A' ~ epsilondb$Acorr[epsilondb$base == 'G']
        ),
        nucleo == 'C' ~ case_when(
          nn == 'G' ~ epsilondb$Gcorr[epsilondb$base == 'C'],
          nn == 'C' ~ epsilondb$Ccorr[epsilondb$base == 'C'],
          nn == 'T' ~ epsilondb$Tcorr[epsilondb$base == 'C'],
          nn == 'A' ~ epsilondb$Acorr[epsilondb$base == 'C']
        ),
        nucleo == 'T' ~ case_when(
```

```
          nn == 'G' ~ epsilondb$Gcorr[epsilondb$base == 'T'],
          nn == 'C' ~ epsilondb$Ccorr[epsilondb$base == 'T'],
          nn == 'T' ~ epsilondb$Tcorr[epsilondb$base == 'T'],
          nn == 'A' ~ epsilondb$Acorr[epsilondb$base == 'T']
        ),
        nucleo == 'A' ~ case_when(
          nn == 'G' ~ epsilondb$Gcorr[epsilondb$base == 'A'],
          nn == 'C' ~ epsilondb$Ccorr[epsilondb$base == 'A'],
          nn == 'T' ~ epsilondb$Tcorr[epsilondb$base == 'A'],
          nn == 'A' ~ epsilondb$Acorr[epsilondb$base == 'A']
        )
      )
    )

  #attributes 0 to the first nucleobase individual contribution
  epsilon.calc$indiv.base.cont[1] = 0
  #attributes 0 to the last nucleobase individual contribution
  epsilon.calc$indiv.base.cont[str_length(sequence)] = 0

epsilon.calc
#>   position nucleo nn indiv.base.cont nn.cont
#> 1        1      G  C               0   17600
#> 2        2      C  A            7400   21200
#> 3        3      A  T           15400   22800
#> 4        4      T                  0      NA
```

Finally, the sum of individual contributions are subtracted from the nearest neighbor contributions:

```
#sum of indiv cont subtracted from sum of nn cont.
result <- sum(epsilon.calc$nn.cont, na.rm = T) - sum(epsilon.calc$indiv.base.cont, na.rm = T)
result
#> [1] 38800
```

###### 4.1.3 Use

`epsilon.calculator` computes the molar extinction coefficient at 260 nm of oligonucleotides from their sequences. So far, it only works for DNA oligonucleotides, using the four canonical nucleotides.

Below is an example for a single sequence:

```
epsilon.calculator("GGGTTAGGGTTAGGGTTAGGG")
#> [1] 215000
```

The sequence must be provided as a string, and **must** be written with upper case letters (to allow the implementation of RNA calculation in the future):

```
epsilon.calculator("gggttagggttagggttaggg")
#> [1] 0
```

`epsilon.calculator` can be applied on a list of sequence (here, `oligo.list`) using the base function `lapply`:

```r
oligo.list <- c('oligo name 1' = 'GGGTTAGGGTTAGGGTTAGGG', 'oligo name 2' = 'TGGGGT',
                'oligo name 3' = 'GCAT', 'oligo name 4' = 'TACG')

epsilon.list <- lapply(oligo.list, epsilon.calculator)

epsilon.list
#> $`oligo name 1`
#> [1] 215000
#>
#> $`oligo name 2`
#> [1] 57800
#>
#> $`oligo name 3`
#> [1] 38800
#>
#> $`oligo name 4`
#> [1] 39800
```

or on a data frame (here, `df`) to directly associate the results to other variables, as is performed within `g4db`.

```r
df <- data.frame(
  oligo = c('name 1', 'name 2', 'name 3', 'name 4'),
  something = c('a', 'b', 'c', 'd'),
  sequence = c('GGGTTAGGGTTAGGGTTAGGG', 'TGGGGT', 'GCAT', 'TACG')
)

df$epsilon <- lapply(df$sequence, epsilon.calculator)

df
#>    oligo something              sequence epsilon
#> 1 name 1         a GGGTTAGGGTTAGGGTTAGGG  215000
#> 2 name 2         b                TGGGGT   57800
#> 3 name 3         c                  GCAT   38800
#> 4 name 4         d                  TACG   39800
```

#### 4.2 mass.diet

##### 4.2.1 Principle

The *importR* tab includes an optional mass spectrometric data reduction step, performed by the `mass.diet` function. It applies two different filters:

- An *m/z* filter, which exclude all data points above or below a user-supplied *m/z* range,
- An *intensity* filter, which excludes data points whose intensity is below a threshold. This intensity threshold is calculated as the mean *intensity* of a user-supplied *m/z baseline* range of length *n*, multiplied by a user-supplied *coefficient.*

$$threshold = \frac{\sum_{baseline_{start}}^{baseline_{end}} intensity}{n} \times coefficient$$

When submitting several spectra, the intensity thresholds are computed for each individual spectrum to avoid issues with different signal-to-noise ratios.

##### 4.2.2 Code

The code of `mass.diet` is contained in `R/massdiet.R`.

`mass.diet` requires that the data is formatted as a dataframe with the following columns:

- `mz`, the m/z axis,
- `int`, the intensity,
- `oligo`, the oligonucleotide names,
- `buffer.id`, the buffer name,
- `tune`, the MS tune name,
- `rep`, the replicate number

The last four columns are used as grouping variables to calculate individual intensity thresholds.

The data is processed in three simple steps. First the *m/z* range filter is applied, then the intensity threshold is calculated for each spectrum from the average noise in the defined baseline, and finally the intensity thresholds are applied to their respective spectrum. If the user lets the coefficient to its default value, i.e. 0, no intensity filtering will happen.

```r
mass.diet <- function(fat.mass, base.start, base.end, range.start, range.end, baseline.int){

  library(tidyverse)

  #m/z range filtering----
  losing.mass <- fat.mass %>%
    filter(mz > min(range.start)) %>%
    filter(mz < max(range.end))

  #intensity filtering----
  #intensity threshold determination
  if (baseline.int > 0) { #filters by intensity if the coefficient is not 0
    baseline.filter <- losing.mass %>%
      group_by(oligo, buffer.id, tune, rep) %>% #grouping by individual spectra
      filter(mz < base.end) %>% #selection of baseline range
      filter(mz > base.start) %>%
      #intensity threshold (mean noise times the multiplier)
      summarise(basemean = mean(int)*baseline.int)

    #removal of noise
    fit.mass <- losing.mass %>% #joins threshold to m/z filtered data
      left_join(baseline.filter, by = c("oligo", "buffer.id", "tune", 'rep')) %>%
      group_by(oligo, buffer.id, tune, rep) %>% #group by spectrum
      filter(int > basemean) %>% #filters
      select(-c(basemean)) #removes threshold column
  } else {
    #does nothing if coefficient at 0
    fit.mass <- losing.mass
  }
  return(fit.mass)
}
```

##### 4.2.3 Use

`mass.diet` can be used outside of *g4db*, provided the input data contains the above-mentioned columns.

Here, we will use the data from the demo input file. In *g4db* it is loaded as follows:

```r
library(readxl)
library(hablar)

wide.input <- read_excel(system.file("extdata/demo_input.xlsx", package = 'g4dbr'),
                         sheet = "MS")

#extract descriptors
descriptors <- wide.input %>%
  slice(1:6)

#extract data
wide.input <- wide.input %>%
  slice(-1:-6)

data.collector <- data.frame()

for (i in 1:ncol(wide.input)-1) {
  if (i %% 2 != 0) { #runs on uneven columns only
    buffer <- wide.input %>%
      select(i, i+1) %>% #select every couple columns
      mutate(descriptors[[1, i+1]], #adds columns for descriptors
             descriptors[[2, i+1]],
             descriptors[[3, i+1]],
             descriptors[[4, i+1]],
             descriptors[[5, i+1]]) %>%
      magrittr::set_colnames(
        c('mz', 'int', 'oligo', 'buffer', 'cation', 'tune', 'rep')
        ) %>%
      mutate(buffer.id = ifelse(is.na(cation), buffer, paste(buffer, '+', cation))) %>%
      convert(num('mz', 'int')) #converts some columns to numeric type
    #binds data
    data.collector <- rbind(data.collector, buffer,
                            make.row.names = F)
  }
}

wide.input <- data.frame() #empty for memory
buffer <- data.frame() #same

data.collector
#> # A tibble: 1,268,904 x 8
#>        mz   int oligo buffer cation tune  rep   buffer.id
#>     <dbl> <dbl> <chr> <chr>  <chr>  <chr> <chr> <chr>
#>  1   300.     0 1XAV  TMAA   <NA>   tune1 1     TMAA
#>  2   300.     1 1XAV  TMAA   <NA>   tune1 1     TMAA
#>  3   300.     5 1XAV  TMAA   <NA>   tune1 1     TMAA
#>  4   300.     0 1XAV  TMAA   <NA>   tune1 1     TMAA
#>  5   300.    10 1XAV  TMAA   <NA>   tune1 1     TMAA
#>  6   300.    38 1XAV  TMAA   <NA>   tune1 1     TMAA
#>  7   300.    72 1XAV  TMAA   <NA>   tune1 1     TMAA
#>  8   300.    72 1XAV  TMAA   <NA>   tune1 1     TMAA
#>  9   300.    53 1XAV  TMAA   <NA>   tune1 1     TMAA
```

```
#> 10  300.     33 1XAV  TMAA   <NA>   tune1 1     TMAA
#> # ... with 1,268,894 more rows
```

`mass.diet` is applied as shown below, by specifying the *m/z* range with `range.start` and `range.end`, the baseline for noise with `base.start` and `base.end`, and the coefficient with `baseline.int`.

```
reduced.data <- mass.diet(fat.mass = data.collector, base.start = 1250, base.end = 1350,
                          range.start = 1000, range.end = 2000, baseline.int = 2)

reduced.data
#> # A tibble: 98,998 x 8
#> # Groups:   oligo, buffer.id, tune, rep [4]
#>        mz   int oligo buffer cation tune  rep   buffer.id
#>     <dbl> <dbl> <chr> <chr>  <chr>  <chr> <chr> <chr>
#>  1 1000.   326 1XAV  TMAA   <NA>   tune1 1     TMAA
#>  2 1000.   374 1XAV  TMAA   <NA>   tune1 1     TMAA
#>  3 1000.   378 1XAV  TMAA   <NA>   tune1 1     TMAA
#>  4 1000.   358 1XAV  TMAA   <NA>   tune1 1     TMAA
#>  5 1000.   432 1XAV  TMAA   <NA>   tune1 1     TMAA
#>  6 1000.   692 1XAV  TMAA   <NA>   tune1 1     TMAA
#>  7 1000.  1057 1XAV  TMAA   <NA>   tune1 1     TMAA
#>  8 1000.  1426 1XAV  TMAA   <NA>   tune1 1     TMAA
#>  9 1000.  1751 1XAV  TMAA   <NA>   tune1 1     TMAA
#> 10 1000.  1817 1XAV  TMAA   <NA>   tune1 1     TMAA
#> # ... with 98,988 more rows
```

Here, the 1250-1350 *m/z* region was picked for the baseline with a *coefficient* of 2, and the *m/z* was restricted to 1000-2000. This reduced the number of data points to 7% of its original value (from 1,268,904 to 98,998). That being said, `mass.diet` should be used conservatively and the size-reduced data *must be inspected* visually for excess removal.

Below are the four mass spectra from the demo file after running `mass.diet`.

```
library(ggthemes)

reduced.data %>%
  #normalization
  group_by(oligo, buffer.id) %>%
  mutate(int.min = min(int), int.max = max(int)) %>%
  mutate(norm.int = (int - int.min)/(int.max - int.min)) %>%
  #plot
  ggplot(aes(x = mz, y = norm.int, color = paste(oligo, buffer.id))) +
  geom_line() +
  xlab("m/z") +
  ylab("intensity") +
  facet_grid(buffer.id~oligo) +
  theme_pander() +
  theme(legend.position = 'none')
```

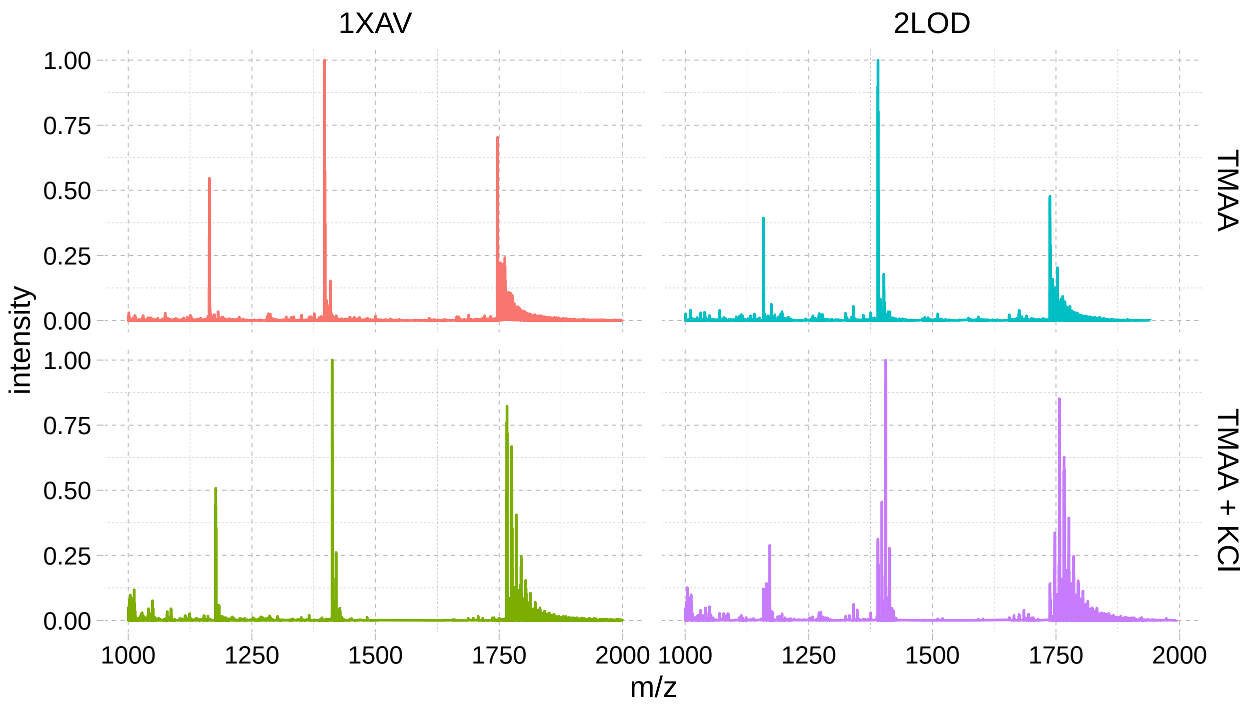

Figure S192: Normalized native MS spectra from the demo input file data reduced to a fraction of its original size using ‘mass.diet‘

#### 4.3 database.eraser

##### 4.3.1 Principle

The `database.eraser` function reads a user-specified database, remove the data for the indicated *oligonucleotides* and analytical *methods*, and returns a list of dataframe (one dataframe per *method*). Specifically, the `erase.db` function, which filters off the data of the indicated `oligos`, is applied method per method, and only on those specified by logical values `erase.CD`, `erase.MS`, `erase.NMR` and `erase.UV`. This way, it maintains the data frames structures even if all data is removed, which allows to reuse the file in *g4db*.

##### 4.3.2 Code

```
database.eraser <- function(db.to.erase = NULL, remove.oligos = NULL,
                            erase.CD, erase.NMR, erase.MS, erase.UV){

  #operator definition
  '%notin%' <- Negate('%in%')

  #data to remove
  remove.oligos <- remove.oligos

  #if all exp data is removed, remove the oligo info as well
  if (erase.CD == TRUE & erase.NMR == TRUE & erase.MS == TRUE & erase.UV == TRUE) {
    erase.info <- TRUE
```

```
  } else {
    erase.info <- FALSE
  }

  #file loading
  load(file = db.to.erase)

  #erasing function
  erase.db <- function(dataset = NULL, remove.oligos){

    dataset <- dataset %>%
      filter(oligo %notin% remove.oligos)

    return(dataset)
  }

  #Data removal (per method, if selected for removal)
  if (erase.CD == TRUE) {
    db.CD <- as.data.frame(erase.db(dataset = db.CD, remove.oligos))
  }

  if (erase.info == TRUE) {
    db.info <- as.data.frame(erase.db(db.info, remove.oligos))
  }

  if (erase.MS == TRUE) {
    db.MS <- as.data.frame(erase.db(db.MS, remove.oligos))
  }

  if (erase.UV == TRUE) {
    db.UV <- as.data.frame(erase.db(db.UV, remove.oligos))
  }

  if (erase.NMR == TRUE) {
    db.NMR <- as.data.frame(erase.db(db.NMR, remove.oligos))
  }

  #Rest of data collected back in a list
  db.collection <- list('db.info' = db.info,
                        'db.CD' = db.CD,
                        'db.NMR' = db.NMR,
                        'db.MS' = db.MS,
                        'db.UV' = db.UV)

  return(db.collection)

}
```

###### 4.3.3 Use

Below is an example for the demo database, for which the MS and NMR data will be removed for both entries.

```
modified.db <- database.eraser(db.to.erase = system.file('extdata/demo_database.Rda', package = 'g4dbr'
                remove.oligos = c('1XAV', '2LOD'),
                erase.CD = FALSE, erase.NMR = TRUE, erase.MS = TRUE, erase.UV = FALSE)
```

Both entries are still present in the database:

```
head(modified.db[["db.info"]])
#>   oligo                                                                 DOI submitted_by  depo.date
#> 1  1XAV   <a href=http://dx.doi.org/10.1021/bi048242p>10.1021/bi048242p</a>           AG 2020-06-26
#> 2  2LOD <a href=http://dx.doi.org/10.1093/nar/gks329>10.1093/nar/gks329</a>           AG 2020-06-26
```

And the UV and CD data are still present:

```
db.CD <- modified.db[["db.CD"]]
db.UV <- modified.db[["db.UV"]]
head(db.UV)
#>   T.unk abs.raw abs.blk oligo            buffer    cation rep melt.l melt.c                        co
#> 1   4.2  0.2711       0  1XAV 25 mM Kp (pH 7.0) 70 mM KCl   1      1     10 25 mM Kp (pH 7.0) + 70 m
#> 2   4.4  0.2711       0  1XAV 25 mM Kp (pH 7.0) 70 mM KCl   1      1     10 25 mM Kp (pH 7.0) + 70 m
#> 3   4.6  0.2715       0  1XAV 25 mM Kp (pH 7.0) 70 mM KCl   1      1     10 25 mM Kp (pH 7.0) + 70 m
#> 4   4.8  0.2717       0  1XAV 25 mM Kp (pH 7.0) 70 mM KCl   1      1     10 25 mM Kp (pH 7.0) + 70 m
#> 5   5.0  0.2718       0  1XAV 25 mM Kp (pH 7.0) 70 mM KCl   1      1     10 25 mM Kp (pH 7.0) + 70 m
#> 6   5.3  0.2717       0  1XAV 25 mM Kp (pH 7.0) 70 mM KCl   1      1     10 25 mM Kp (pH 7.0) + 70 m
head(db.CD)
#>      wl            CD oligo              buffer cation   l con            buffer.id delta.epsilon
#> 1 329.8  0.0274670185  1XAV 100 mM TMAA (pH 7.0)   none 0.4  10 100 mM TMAA (pH 7.0)   0.208209661
#> 2 329.6  0.0096042216  1XAV 100 mM TMAA (pH 7.0)   none 0.4  10 100 mM TMAA (pH 7.0)   0.072803378
#> 3 329.4 -0.0002250792  1XAV 100 mM TMAA (pH 7.0)   none 0.4  10 100 mM TMAA (pH 7.0)  -0.001706179 -
#> 4 329.2 -0.0025197889  1XAV 100 mM TMAA (pH 7.0)   none 0.4  10 100 mM TMAA (pH 7.0)  -0.019100886 -
#> 5 329.0 -0.0108839050  1XAV 100 mM TMAA (pH 7.0)   none 0.4  10 100 mM TMAA (pH 7.0)  -0.082503828 -
#> 6 328.8 -0.0103693931  1XAV 100 mM TMAA (pH 7.0)   none 0.4  10 100 mM TMAA (pH 7.0)  -0.078603647 -
```

But the MS and NMR data have been removed for both oligonucleotides. Note that the dataframe structure is conserved:

```
db.NMR <- modified.db[["db.NMR"]]
db.MS <- modified.db[["db.MS"]]
head(db.MS)
#>         mz int oligo              buffer   cation   tune rep                         buffer.id int.mi
#> 1 1250.003 103 oligo 100 mM TMAA (pH 7.0) 1 mM KCl tune99  99 100 mM TMAA (pH 7.0) + 1 mM KCl
head(db.CD)
#>      wl            CD oligo              buffer cation   l con            buffer.id delta.epsilon
#> 1 329.8  0.0274670185  1XAV 100 mM TMAA (pH 7.0)   none 0.4  10 100 mM TMAA (pH 7.0)   0.208209661
#> 2 329.6  0.0096042216  1XAV 100 mM TMAA (pH 7.0)   none 0.4  10 100 mM TMAA (pH 7.0)   0.072803378
#> 3 329.4 -0.0002250792  1XAV 100 mM TMAA (pH 7.0)   none 0.4  10 100 mM TMAA (pH 7.0)  -0.001706179 -
#> 4 329.2 -0.0025197889  1XAV 100 mM TMAA (pH 7.0)   none 0.4  10 100 mM TMAA (pH 7.0)  -0.019100886 -
#> 5 329.0 -0.0108839050  1XAV 100 mM TMAA (pH 7.0)   none 0.4  10 100 mM TMAA (pH 7.0)  -0.082503828 -
#> 6 328.8 -0.0103693931  1XAV 100 mM TMAA (pH 7.0)   none 0.4  10 100 mM TMAA (pH 7.0)  -0.078603647 -
```

To save the modified database, use the `save` function:

```
db.info <- modified.db[["db.info"]]

save(db.info,
     db.CD,
     db.NMR,
     db.MS,
     db.UV,
     file = 'filepath/filename.rda')
```
